# ADTnorm: Robust Integration of Single-cell Protein Measurement across CITE-seq Datasets

**DOI:** 10.1101/2022.04.29.489989

**Authors:** Ye Zheng, Daniel P. Caron, Ju Yeong Kim, Seong-Hwan Jun, Yuan Tian, Mair Florian, Kenneth D. Stuart, Peter A. Sims, Raphael Gottardo

## Abstract

CITE-seq enables paired measurement of surface protein and mRNA expression in single cells using antibodies conjugated to oligonucleotide tags. Due to the high copy number of surface protein molecules, sequencing antibody-derived tags (ADTs) allows for robust protein detection, improving cell-type identification. However, variability in antibody staining leads to batch effects in the ADT expression, obscuring biological variation, reducing interpretability, and obstructing cross-study analyses. Here, we present ADTnorm (https://github.com/yezhengSTAT/ADTnorm), a normalization and integration method designed explicitly for ADT abundance. Benchmarking against 14 existing scaling and normalization methods, we show that ADTnorm accurately aligns populations with negative- and positive-expression of surface protein markers across 13 public datasets, effectively removing technical variation across batches and improving cell-type separation. ADTnorm enables efficient integration of public CITE-seq datasets, each with unique experimental designs, paving the way for atlas-level analyses. Beyond normalization, ADTnorm includes built-in utilities to aid in automated threshold-gating as well as assessment of antibody staining quality for titration optimization and antibody panel selection. Applying ADTnorm to a published COVID-19 CITE-seq dataset allowed for identifying previously undetected disease-associated markers, illustrating a broad utility in biological applications.

## Main

Recent advances in single-cell multimodal profiling, such as Cellular Indexing of Transcriptomes and Epitopes by sequencing (CITE-seq), have enabled the paired profiling of gene expression alongside surface protein expression^1–4^. This paired multimodal profiling of single cells has allowed researchers to achieve more precise cell-type annotation (e.g., of immune cells)^5, 6^, study the relationship between transcriptomic state and surface phenotype^7–9^, and readily adapt results to flow cytometry for validation^1, 4^. Given its extraordinary potential, there is increasing application of CITE-seq for atlas construction^10–12^ and in large cohort disease-related studies^13–15^. To effectively leverage the data being generated, there is a pressing need for computational tools for CITE-seq data integration across studies.

Surface proteome profiling by CITE-seq gives rise to specific data characteristics and sources of technical noise inherent to antibody staining. Owing to the high copy number of surface proteins and efficient molecular capture of antibody-derived tags (ADTs), protein expression is considerably less sparse than other single-cell modalities such as mRNA expression or genome-wide chromatin accessibility. Consequently, the protein expression captured by CITE-seq often closely matches the information-rich multi-peak density distributions observed in flow cytometry^1^ (Supplementary Fig. 1A). Density distributions of protein expression of CITE-seq data frequently exhibit a negative peak, representing background signal arising from non-specifically bound or unbound (free-floating) antibody^16^, and one or more positive peak(s) representing cells expressing the target protein. Similar to fluorescence-based techniques, the signal-to-noise ratio between the negative- and positive-expression peak(s) is highly sensitive to antibody staining conditions, including antibody concentrations^17^, staining volumes and time^18^, and antibody panel composition^19^. Because of these unique considerations, the normalization and integration approaches devised for other single-cell modalities may not be directly translatable, highlighting the need for methodologies tailored to the intricacies of protein data.

Recent normalization algorithms designed for CITE-seq data, similar to established scRNA-seq approaches^20, 21^, have primarily focused on modeling sequencing bias and ambient expression to remove background signals. Centered log-ratio (CLR) normalization was initially proposed for CITE-seq^1^, using library size to account for variable sequencing depth and cell size. However, unlike scRNA-seq, which offers relatively unbiased transcriptional profiling, CITE-seq protein panels target only a handful of manually selected proteins, typically between 10 and 300. Therefore, the overall ADT library size is highly sensitive to panel composition, can be easily skewed by high expression of a few subset-specific proteins, and unreliably reflects sequencing depth or cell size. More sophisticated algorithms, including totalVI^22^, DSB^16^, and DecontPro^23^, attempt to model ambient contamination and remove or re-center the background-signal to zero. However, these negative-expression peaks in ADT abundance mirror expression distributions by conventional cytometry and are essential for reliable threshold-gating of cells for cell-type annotation^24^. Improper or incomplete removal of background ADT expression can make it difficult to distinguish between negative-, mid-, and high-expression peaks, for example, in the trimodal expression of the surface marker CD4, a T cell lineage marker. Consequently, normalization of the negative peak in CITE-seq should emphasize its essential role in cell-type identification rather than its artificial removal.

Instead of individually modeling each source of noise, we constructed a non-parametric strategy, ADTnorm, building on methods originally conceived for cytometry data^25^ to remove the batch effects through strategic peak identification and alignment. ADTnorm uses a curve registration algorithm^26^ to identify protein density landmarks, including the negative and positive peaks, and relies on local minima to detect the valleys separating adjacent peaks. Employing a functional data analysis approach^27^, ADTnorm normalizes protein expression by aligning the landmarks across datasets (Fig. 1A and Methods), effectively simulating a scenario where all data are derived from the same experiment with equivalent background and antibody staining quality. ADTnorm is implemented as an R package (https://github.com/yezhengSTAT/ADTnorm) with an interactive graphical user interface to simplify landmark adjustments (Supplementary Fig. 1B) and a Python wrapper (https://github.com/donnafarberlab/ADTnormPy) available to facilitate ADT-norm’s integration into existing CITE-seq analysis workflows (Supplementary Note).

Leveraging 13 public CITE-seq datasets (Supplementary Table 1), we benchmarked the integration performance of ADTnorm against 14 methods from three broad groups: (1) scaling methods commonly applied to cytometry and single-cell data, including Arcsinh transformation, CLR^1^, log-transformation of count per million (logCPM), and a hybrid approach combining Arcsinh and CLR transformations (Arcsinh + CLR); (2) popular single-cell batch effect removal tools, including Harmony^28^ implemented on the raw counts, arcsinh-transformed, CLR-transformed or logCPM-transformed data, fastMNN^29^, and CytofRUV^30^; and (3) methods tailored to CITE-seq normalization, including DSB^16^, decontPro^23^, totalVI^22^, and sciPENN^8^. Across 13 datasets, ADTnorm effectively reduced batch variability, such that negative and positive populations for each surface protein marker could be consistently identified across studies (Fig. 1B, protein density distributions in Supplementary Note). UMAP embeddings of the normalized ADT expression revealed effective batch integration by ADTnorm while preserving cell type separation at both broad and refined annotation levels, treating either the study-level or individual samples as batches (Supplementary Figs. 2-3). ADTnorm applied using default landmark detection (default) or manually adjusted landmark detection (customized; Methods) outperformed other tools in balancing cell-type separation with cross-study batch effect removal as quantified by Silhouette scores, Adjust Rand Index (ARI), and the Local Inverse Simposon’s Index (LISI) (Methods; Fig. 1C and Supplementary Fig. 4A-C). Furthermore, ADTnorm can facilitate the seamless integration of new datasets without reprocessing existing ones by aligning landmarks to predetermined locations (Supplementary Note). It can also incorporate users’ prior knowledge about a batch’s cell type composition. For example, because the *Buus 2021 T* cell dataset is composed of only T cells, ADTnorm is adjusted to align the singular peak in CD3 as positive-expression (Fig. 1B and Supplementary Note). ADTnorm is also highly scalable, with a fast processing speed and low memory consumption compared to other methods (Supplementary Fig. 4D-E). Also, ADTnorm is designed to process protein markers independently, allowing adaption to parallel processing.

**Figure 1.**
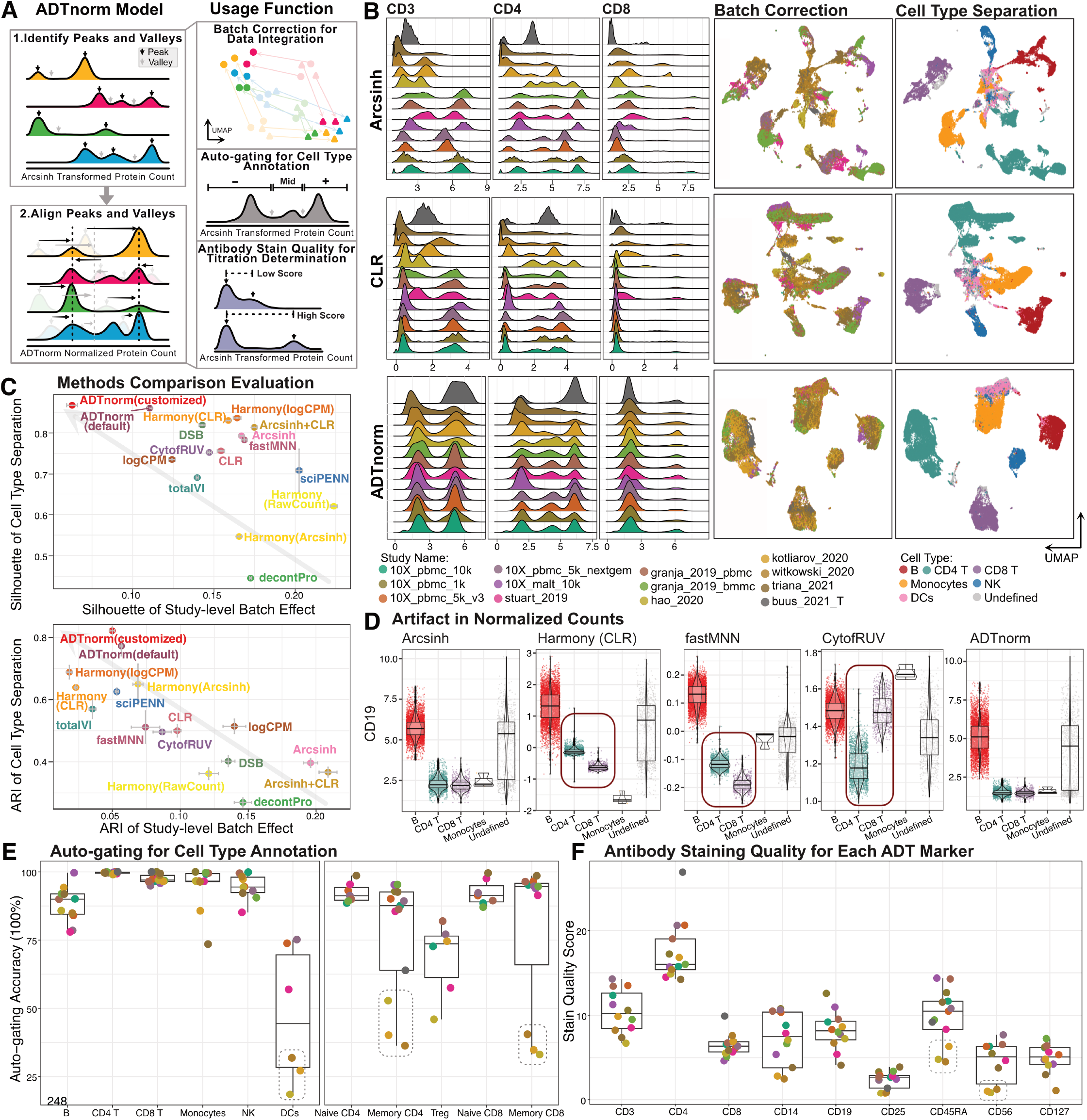
ADTnorm normalization model, function and performance. **A**. ADT-norm takes in the ADT expression matrix after routine quality control steps. The normalization procedure starts with the identification of landmarks (peaks and valleys) in the expression density distribution for each protein marker of each batch. Then, detected peaks and valleys are aligned across batches through functional data analysis. The landmark alignment normalization integrates CITE-seq data from different sources. The detected peaks and valleys can also be used for automatic threshold-gating (auto-gating) and antibody staining quality evaluation, which can guide the selection of CITE-seq antibodies and staining concentrations. **B**. Comparison of ADT expression distribution across studies of three T cell lineage markers (CD3, CD4 and CD8) after normalization by Arcsinh, CLR or ADTnorm. UMAP embeddings generated after normalization of 9 shared ADT markers were used across all 13 studies colored by study or cell type. ADTnorm was provided “study” as the batch key. Cell type annotations were defined by manual threshold-gating by two immunologists on each sample separately, independent of the normalization work in this paper (Methods). The corresponding manual gating strategy is summarized in Supplementary Table 2. **C**. Study-level batch correction and broad-level cell type separation quantified by Silhouette score and Adjusted Rand Index (ARI) across various ADT transformation and normalization approaches (Methods). ADTnorm was applied using default parameters or customized landmark alignment adjustments. Gray arrows indicate the direction of improved integration performance, i.e., minimized batch effect and maximized cell type separation. The vertical and horizontal error bars represent the standard deviations of 20 bootstrap samples for each normalization method. **D**. Violin plots displaying CD19 expression for each cell in the 10X malt 10k dataset following normalization under the severe imbalanced setting (Methods). Abnormal artifacts introduced to specific cell types during the normalization are highlighted by red squares. **E**. Average auto-gating accuracy across cell types (x-axis) and studies (colors). **F**. Averaged stain quality quantification across protein markers (x-axis) and studies (colors). The central boxes of **D**-**F** represent the interquartile range (IQR), which contains the middle 50% of the data. The line inside the box indicates the median. The whiskers extend to the smallest and largest values within 1.5 times the IQR from the lower and upper quartiles, respectively. **B, E** and **F** share the same color legend for studies.

We next explored the downstream impact of protein normalization on joint embeddings of RNA and protein data. Following batch-correction of ADT expression by the above methods and batch-correction of the RNA expression using reciprocal PCA^10^, we computed the multimodal embedding using the weighted nearest neighbor (WNN) algorithm^10^ (Supplementary Fig. 5A and Methods). As totalVI and sciPENN already incorporate gene expression into their protein normalization process, we omitted them from the WNN integration comparison. As expected, methods with sub-optimal removal of ADT batch effects resulted in skewed WNN integration (Supplementary Fig. 6). ADTnorm markedly minimized batch influences and achieved superior accuracy in segregating cell types as quantified by ARI (Supplementary Fig. 5B), underscoring its utility in post-normalization multimodal integration.

As surface protein expression varies across cell types, batch correction may be sensitive to variable subset composition across batches. To evaluate the resilience of normalization methods, we subsampled specific cell subsets from a few batches, devising three scenarios featuring increasingly skewed cell-type compositions (Methods). Careful examination revealed that Harmony, fastMNN, and CytofRUV were highly sensitive to compositional differences, producing unexpected and inaccurate results. For example, CD19 is a highly specific B cell-lineage marker. However, in some batches, Harmony- and fastMNN-normalized CD19 expression was significantly higher in CD4 T cells than in CD8 T cells, and CytofRUV-normalized CD19 expression in CD8 T cells was comparable to that in B cells, patterns not supported by biological expectations (Fig. 1D and Supplementary Fig. 7). Similar discrepancies were noted with DSB, totalVI, and sciPENN across other vital lineage markers (Supplementary Figs. 8-9). ADTnorm distinguishes itself by meticulously preserving the ranking of protein expression across cells within each batch, thereby reducing the risk of biologically irrelevant anomalies.

Beyond its primary role in batch correction, ADTnorm leverages intermediate landmark detection results to perform automated threshold-gating (auto-gating) for cell type annotation and to assess staining quality to aid in the optimization of CITE-seq experiments (Methods). Valley landmarks identified during ADTnorm normalization can be used to perform automated cell type annotation using predefined gating rules (Supplementary Table 2; Supplementary Fig. 10A-C). While ADTnorm auto-gating showcased high accuracy for a majority of the studies, achieving between 80-100% for comprehensive and nuanced cell type distinctions, autogating was underperformed for dendritic cells, memory CD4 T and memory CD8 T cells in the *Hao 2020, Kotliarov 2020*, and *Witkowski 2020* datasets (Fig. 1E). Auto-gating accuracy is likely influenced by the marker staining quality. Hence, we introduced a stain quality score, inspired by fluorescent stain index^31^, to detect protein markers with poor signal-to-noise separation (Methods; Supplementary Fig. 1C). Low-quality scores are suggestive of under-optimized staining conditions, which need careful evaluation or potential exclusion from downstream analyses. Leveraging ADTnorm to assess staining quality revealed that CD56 and CD45RA, which are markers used for gating dendritic and memory T cells, featured less distinct peak separation in batches with poor auto-gating performance (Fig. 1F and Supplementary Fig. 10D).

To effectively stain for surface protein, antibody concentrations must be carefully tuned for each sample type. Sufficient antibodies are essential for positive-expression signal(s) to overcome background, but an overabundance of antibodies can obscure rare or low-expression markers by increasing background noise and can increase experimental costs. Although down-stream analysis can often tolerate suboptimal staining conditions, variable staining quality is a major source of batch artifacts across samples and laboratories. To explore whether our stain quality score is sensitive enough for titration optimization and to evaluate ADTnorm’s ability to mitigate these batch effects, we utilized a titration CITE-seq study that analyzed 124 antibodies on human peripheral blood mononuclear cells (PBMCs)^17^. This study categorized antibody titration into four levels, including the manufacturer’s recommended concentration (1x) and adjustments to 1/25x, 1/5x, and double (2x) the recommended concentration. As anticipated, the higher concentrations (1x and 2x) typically yielded more distinct separation between negative and positive cell populations, whereas lower concentrations led to greater overlap between negative and positive populations or failed to identify any positive population (Fig. 2A and Supplementary Note). These trends were reflected in the stain quality scores, where markers with reduced separation at low antibody concentrations exhibited lower scores (Fig. 2B). Notably, conventional normalization methods were unable to successfully integrate expression across titration batches (Supplementary Fig. 11 and Supplementary Note), but ADTnorm could effectively align negative and positive populations across concentrations, thus rescuing cell type discrimination for many protein markers profiled using sub-optimal staining conditions (Supplementary Fig. 12) and minimizing batch effects (Fig. 2A). For markers at low titrations that exhibited no positive population, ADTnorm could only align the negative populations (Supplementary Fig. 13A). In these cases, excessively low stain quality scores could alert researchers of protein markers that consistently show poor discrimination, suggesting a potential need for revising antibody titration or selection (Fig. 2B and Supplementary Fig. 13B). We also assessed the influence of antibody titration on ADTnorm’s auto-gating accuracy, finding that auto-gating accuracy remains stable as long as lineage markers had detectable positive staining (Supplementary Fig. 14).

**Figure 2.**
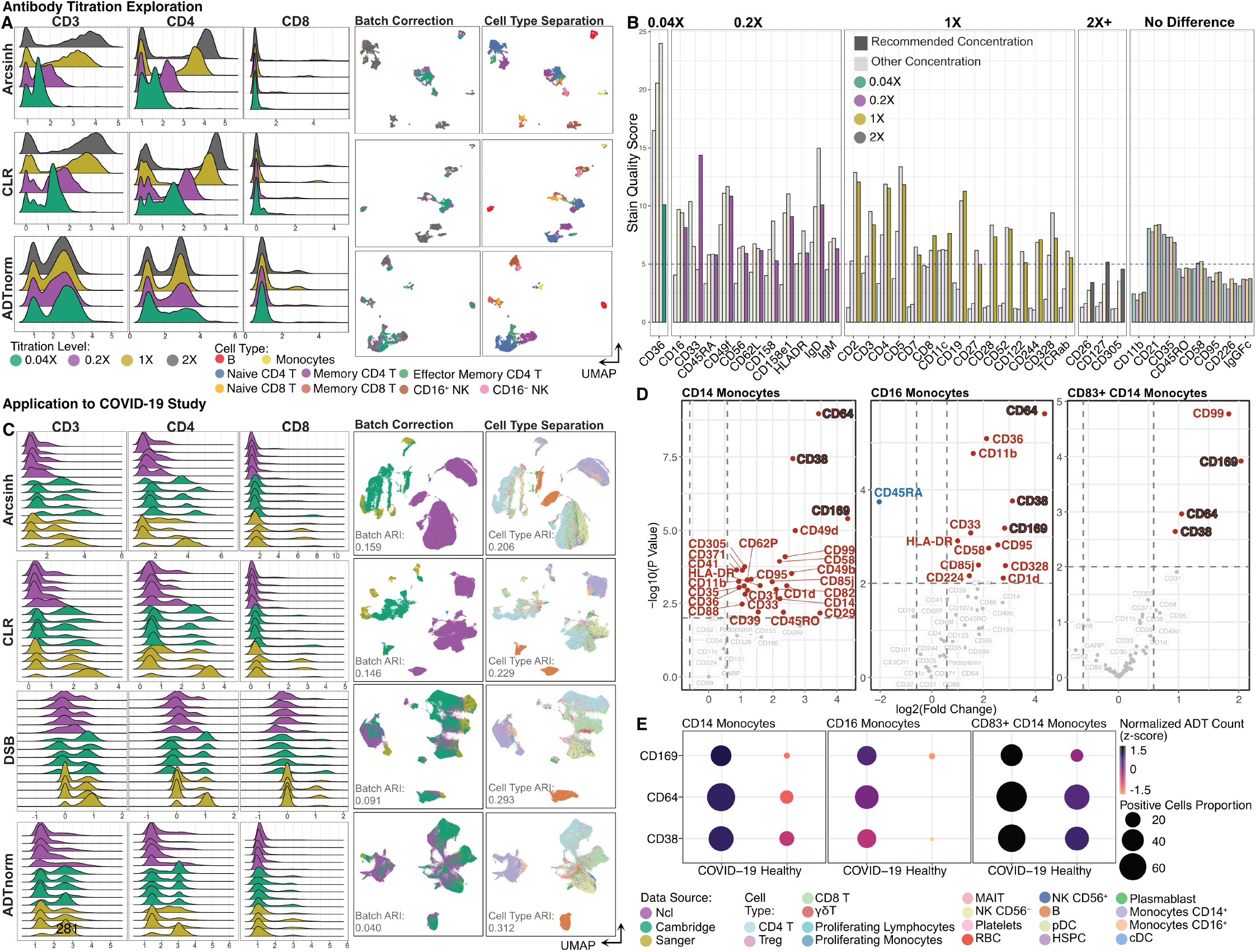
ADTnorm application to antibody titration determination and COVID-19 related disease study. **A**. ADT expression distributions of three T cell lineage markers (CD3, CD4 and CD8) across samples stained at 1/25, 1/5, 1 and 2 times the commercially recommended antibody concentration following normalization by Arc-sinh, CLR or ADTnorm. UMAP displayed the batch correction across the four antibody concentrations and cell-type separation using 124 ADT markers provided by the original paper^17^. **B**. Stain quality score is utilized to determine the positive population and negative population separation power (Methods). The lowest titration with sufficient separation of positive and negative cells (dashed line indicates stain quality score of 5) is highlighted for each protein marker with increased saturation. **C**. Data integration across three research institutes where CITE-seq was generated. UMAP shows the batch correction across three research institutes and cell type separation compared across Arcsinh, CLR, DSB and ADTnorm. DSB is the normalization method used in the original paper^13^. UMAPs were constructed on 192 ADT markers colored by research institute or cell type. **D**. Volcano plots displaying results of differential proportion of the positive cells for each protein marker between healthy donors and COVID-19 patients. The differential detection analysis was done for CD14^+^ Monocytes, CD16^+^ Monocytes and CD83^+^ CD14^+^ Monocytes, respectively. Cell type labels are from the original publication^13^ of the COVID-19 data. **E**. Dot plot displays consistently differentially expressed protein markers, i.e., CD38, CD64 and CD169, across three monocyte subsets. Points are colored by the average normalized ADT expression and the dot size is relative to the proportion of cells with positive-expression in healthy donors or COVID-19 patients.

We next explored whether ADTnorm could facilitate the analysis of consortium efforts. Three UK medical centers profiled 192 protein markers using CITE-seq to study COVID-19 immune response across a diverse cohort of over 100 healthy donors and COVID-19 patients^13^. Staining quality was highly variable across the participating medical centers (Fig. 2C). Specifically, samples from Newcastle (Ncl) exhibited reduced separation between negative and positive peaks, whereas samples from Cambridge and Sanger displayed robust separation and a higher frequency of detectable positive signals (Supplementary Figs. 15-16). These batch effects could not be effectively mitigated by other tools (Fig. 2C, Supplementary Fig. 17 and Supplementary Note). ADTnorm effectively reduced technical artifacts (Fig. 2C), resulting in improved cell type separation, both at the broad and refined annotation levels and also in the joint RNA and ADT embedding (Supplementary Fig. 18).

Leveraging ADTnorm’s integration and auto-gating, we next aimed to identify whether the expression of specific surface markers could be associated with COVID-19 disease (Supplementary Fig. 16E). Previous studies have identified compositional changes in the immune compartment associated with disease, including increases in the frequency of specific monocyte subsets in the PBMC compartment of mild, moderate, and severe COVID-19 patients (as noted in Fig. 1c of the original publication^13^). Other studies have identified biomarkers on blood monocytes associated with COVID-19 and type-I interferon signaling, including CD38^32,33^, CD64^34,35^ and CD169^36,37^. We sought to identify whether these trends could be attributed to changing subset frequencies within the monocyte compartment or to the upregulation of these markers across multiple subsets of monocytes. We analyzed the percent-positivity of these and other markers on CD14^+^, CD16^+^, and CD83^+^CD14^+^ monocytes, and observed upregulation of these markers among COVID-19 patients compared to healthy donors across multiple monocytes states (Fig. 2D-E and Supplementary Fig. 19A). Such upregulation mirrors the trends observed in scRNA-seq (Supplementary Fig. 19B). The normalization employed in the original publication, DSB, did not accurately represent these trends, masking positive expression of CD169 (Supplementary Fig. 20), failing to identify upregulation of CD169 with COVID-19 in any monocyte subset, and reducing signal of CD38 and CD64 in CD16 monocytes (Supplementary Fig. 19C). This demonstrates the utility of ADTnorm in isolating biologically relevant changes and uncovering previously concealed insights in surface protein expression.

In summary, ADTnorm offers a fast, precise, and scalable solution for normalizing protein expression data, effectively minimizing batch artifacts within studies and enabling integration across studies. ADTnorm is designed for high adaptability, allowing for normalization at various batch levels, supporting missing data, and incorporating prior cell type knowledge. By addressing protein batch effects, ADTnorm also improves multimodal aggregation of RNA and protein modalities, enhancing cell type discrimination and improving interpretability. Unlike other normalization methods that may introduce abnormal expression artifacts, ADTnorm maintains the ranked order of cells within batches for expression of each protein marker and delivers consistent performance across datasets with uneven cell type compositions. Additionally, its auto-gating feature offers an expedited avenue for cell-type annotation. The integrated stain quality scoring system alerts researchers to suboptimal staining and assesses experiment quality, aiding in the calibration of antibody titration for pilot studies tailored to specific tissue systems. Among positive-expressing populations, ADTnorm’s landmark registration approach homogenizes variations in enrichment strength across samples. While it is possible that these variations represent biological differences, that interpretation is confounded by many sources of technical noise, including antibody concentrations, staining conditions, and sequencing artifacts. Notably, ADTnorm also preserves information about the proportion of positive-expressing events in each batch, offering valuable insights into disease status, as exemplified in the COVID-19 case study. This feature underscores the potential of ADTnorm to transcend mere normalization, contributing to the identification of disease-associated protein markers.

Due to ADTnorm’s high adaptability, we expect its utility may also extend beyond CITE-seq, allowing for the harmonization of protein expression across multiple technologies (e.g., flow cytometry, CyTOF, and CITE-seq together). Its application is also primed for expansion to multimodal assays by leveraging the normalized protein data as a bridge for cross-modality integration, such as scCUT&Tag-pro^38^, ASAP-seq^39^ and PHAGE-ATAC^40^, which profile surface proteins alongside epigenomic or chromatin accessibility features. ADTnorm stands as a pivotal tool in the evolving landscape of genomic research, facilitating comprehensive analyses across a broad spectrum of biological conditions and technological platforms.

## Acknowledgments

This work was supported by the National Institutes of Health grant, HG012797, to Y.Z. and Chan Zuckerberg Initiative award, DI-0000000345, to R.G.. D.P.C. was supported by the Columbia University Graduate Training Program in Microbiology and Immunology (T32AI106711). P.A.S. was supported by U19AI128949. We also acknowledge the Scientific Computing Infrastructure at Fred Hutchinson Cancer Center funded by ORIP grant S10OD028685, the J. Orin Edson Foundation, the Translational Data Science Integrated Research Center of the Fred Hutchinson Cancer Center, and NIH U19AI128914. We also appreciate the timely and in-depth discussion with Drs. Helen Lindsay, Bernat Bramon Mora and Antonin Thiebaut from the University of Lausanne.

## Author Contributions

R.G. and Y.Z. conceived the project. Y.Z., D.P.C. and R.G. designed the research and developed the method. Y.Z., J.Y.K. and D.P.C. developed the software and organized the usage manual and tutorial. Y.Z. and S.H.J. designed the auto-gating strategy. Y.T. and M.F. manually gated the protein data to provide a gold standard for the cell type annotation. R.G., P.A.S. and K.D.S provided feedback and suggestions as the project progressed. All authors contributed to the preparation of the manuscript.

## Methods

### Data source and pre-processing

Public CITE-seq datasets were downloaded through URLs summarized in Supplementary Table 1. Datasets are identified by the first author’s last name or by “10X” for data obtained from the 10X genomics websites. ADTnorm was applied to CITE-seq protein expression data after quality checks and cell filtering. Empty droplets, cell aggregates, and apoptotic cells were removed from each dataset based on total UMI counts and the percentage of mitochondrial gene expression using the *PerCellQCMetrics* and *isOutlier* functions using default parameter values from the *scuttle* R package^41^.

### ADTnorm normalization and integration pipeline

#### Landmark Detection

ADTnorm first Arcsinh-transforms raw ADT counts, then identifies landmarks (peaks and valleys) in the density distribution of protein expression. Peaks are defined as local maxima within high-density regions (Supplementary Fig. 1A), and a curve registration algorithm^26^ is employed to identify all detectable peak locations. Between each pair of peaks, ADTnorm identifies valleys as local minima. In scenarios where only one peak is detected or in cases involving a shoulder peak (Supplementary Fig. 1C), valley detection depends on the density slope transitioning from the negative peak to the distribution’s right tail or shoulder peak. Peak and valley detection accuracy relies on precise kernel density estimation for each sample, making selecting a practical bandwidth crucial. The search for an appropriate bandwidth begins with a relatively large value, typically starting at 3. If no or only one peak is detected with this broader bandwidth, the search continues with narrower settings (e.g., 2 and 1). For markers generally exhibiting multiple peaks, like CD4, a narrower bandwidth is applied. Users can input prior information into the ADTnorm software to assist in selecting the optimal bandwidth for constructing the ADT density distribution.

CITE-seq ADT counts are discrete, unlike the continuous data from flow cytometry, with negative peaks often close to zero. Although the Arcsinh transformation effectively compresses large ADT counts into a more manageable range similar to log transformation, it remains nearly linear for counts near zero. Therefore, Arcsinh transformation potentially results in artificial peaks at this low range due to the discrete values. To eliminate suspicious negative peaks, ADTnorm merges peaks detected below a certain small threshold (*neg candidate thres* defined by users in *ADTnorm* function) near zero or applies a larger bandwidth to smooth these areas. Additionally, if the quality control and filtering steps are insufficiently rigorous, leaving empty droplets, a minor enriched peak might appear near zero before the true negative peak. ADTnorm is designed to recognize and disregard such spurious peaks. Conversely, doublets might create false positive peak landmarks outside the typical range. ADTnorm uses the mean absolute deviation (MAD, *mad* function in the *stats* R package with default values) to assess whether a positive peak landmark is an outlier, excluding it from peak alignment procedures. Similarly, outlier valley landmarks that substantially deviate from the typical range of valley values across samples within the same batch are identified by MAD and adjusted to the average valley locations of neighboring samples, i.e., samples with higher protein expression distribution similarity. Such similarity distance between pair of samples is quantified using the earth mover’s distance (EMD, *calculate emd gene* function in the *EMDomics* R package with default values)^42^ based on the ADT count density distribution for each protein marker.

#### Landmark Alignment

ADTnorm leverages identified peaks and valleys in ADT density distributions to mitigate technical variations across batches, studies, platforms, and other experimental inconsistencies by aligning these landmarks across samples. This landmark alignment strategy is inspired by methodologies like guassNorm and fdaNorm^25^, initially developed for flow cytometry data. Specifically, ADTnorm utilizes functional data analysis, employing a warping function^27^ to perform a one-to-one transformation of ADT expression that uniformly adjusts the ADT density distribution in a monotone fashion. Mathematically, the kernel density estimate for each sample *i* is represented by a B-spline interpoland *x*_*i*_. The peak(s) and valley(s) detected for each sample serve as landmarks and the landmark locations are denoted by *t*_*ij*_ where *j* = 1, .., *m. m* is 2 meaning there is only one peak and one valley and *m* is 3 indicating that this sample has two peaks and one valley. To align the peaks and valleys across sample, *x*_*i*_ is transformed by a strictly monotone and invertible function *h*_*i*_ known as a warping function for sample *i*, such that *h*_*i*_(*T*_*start*_) = *T*_*start*_ where *T*_*start*_ is the starting point of the ADT expression value range and *h*_*i*_(*T*_*end*_) = *T*_*end*_ where *T*_*end*_ is the endinng point of the ADT expression value range. Also, *h*_*i*_(*t*_0*j*_) = *t*_*ij*_ for *j* = 1,…, *m*, representing the transformation of the density curves *x*_*i*_ so that the corresponding landmark *j* align to a fixed location *t*_0*j*_. By default, *t*_0*j*_ is set to the mean value of *t*_*ij*_ across samples, but users can pre-defined the target landmark alignment locations. To obtain the optimal estimation of *h*_*i*_, the target function is set to minimizing *∫* ||*y*(*t*) *-xh*(*t*)||^2^*dt* + *λ ∫ ω*^2^(*t*)*dt* where y is a fixed function in the same class as *x*_*i*_ and *ω*(*t*) measures the relative curvature of h. This penalty on the relative curvature ensures that the transformation function is both smooth and monotone.

Note that ADTnorm also allows users to input prior information to more properly align positive peaks across samples. For instance, in batches exclusively involving T cells (e.g., buus 2021 T), a single positive peak for CD3 protein markers is expected. By providing a list of such batches and markers, ADTnorm can precisely align the detected peak to the positive peaks in other samples, ensuring consistent and accurate peak alignment (Supplementary Note). This functionality underscores ADTnorm’s adaptability and effectiveness in handling various experimental conditions and study designs. ADTnorm can be applied to integrate batch effects across studies (Supplementary Fig. 2) or batch effects between individual samples (e.g., each donor is a batch) within studies (Supplementary Fig. 3, Supplementary Note). Furthermore, by ignoring missing values, ADTnorm can be used to integrate ADT expression for markers profiled in some but not all batches, a capability not shared by all normalization methods (Supplementary Note).

### Default and customized ADTnorm normalization settings

In the benchmark analysis with 14 existing methods, ADTnorm normalized the 13 public datasets using default landmark detection (default) or GUI-assisted manually adjusted landmark detection (customized). The default setting applied the default parameter values of the *ADTnorm* R function, which can handle general protein expression normalization scenarios. *ADTnorm* R function offers adjustable parameters to refine landmark detection and provides intermediate density plot visualizations, allowing users to verify the reasonableness of detected peaks and valleys. A detailed tutorial (Supplementary Note and at https://yezhengstat.github.io/ADTnorm/articles/ADTnorm-tutorial.html) is available to facilitate ADTnorm’s usage, offering guidance on software utilization and parameter adjustment to adjust to different protein expression characteristics. Additionally, a GUI implemented using the R shiny function (Supplementary Fig. 1B) is available to help users manually fine-tune landmark locations for tailored protein normalization. The customized setting used in the benchmark analysis relied on manually fine-tuning the peak and valley landmarks to ensure the optimal landmark alignment.

### Weighted nearest neighbor integration of the RNA and protein

Multimodal embeddings were evaluated to test ADT integration performance of ADTnorm and existing methods. The RNA components are integrated using the *Seurat* reciprocal PCA (RPCA) strategy. Specifically, the raw gene expression data are first normalized by log-transformation of count per million (log CPM), and the top 5000 feature genes are selected by the “*vst*” method. Then, the normalized RNA data are scaled using the top features, followed by principal component analysis (PCA) for each study, respectively. Integration anchors are obtained by *FindIntegrationAnchors* function of *Seurat* using the RPCA reduction method. We confirmed the RNA component integration performance by visualizing in UMAP and color-coded by batch, disease status and cell types in Supplementary Fig. 5A. The weighted nearest neighbor (WNN) strategy^10^ from *Seurat* is leveraged to further integrate the harmonized RNA and normalized protein components. Specifically, the *FindMultiModalNeighbors* function from *Seurat* is used to construct the WNN graph based on the top 30 PCs of the RNA component and the top 15 PCs of the protein component. We use default values for all other parameters in the above-mentioned across-modality integration pipeline.

### Robustness evaluation on normalization methods by the imbalanced cell type constitution

To assess the robustness of normalization methods, we filtered the 13 public datasets to create three subsets of the data with different cell-type compositions. In the default integration setting, which we used to illustrate the ADTnorm model and performance, one dataset out of 13 public datasets, i.e., buus 2021 T, was filtered to only contains one sample of 666 T cells. The other 12 datasets profile total PBMCs. This default setting creates a mild imbalanced scenario for data integration. To test integration performance with moderately or severely imbalanced subset compositions across batches, we kept only T cells in the hao 2020 and triana 2021 studies (24 samples and nine samples, respectively), or only CD8 T cells in the triana 2021 study and only T cells from hao 2020 and buus 2021 studies. We evaluated the normalized expression for the CD19 and CD4 across major cell types on the 10X pbmc 10k and 10X malt 10k datasets, which contain one sample per study and the full data from the original studies were kept.

### Stain quality score

To determine the optimal concentration of antibody to stain specific protein markers, we proposed a stain quality score designed for ADT data. The stain quality score is inspired by the stain index widely used to optimize the quality and effectiveness of fluorescent staining of cells in flow cytometry^43^. Stain index is defined as the ratio of the separation between the positive and negative peaks divided by two times the standard deviation of the negative population.

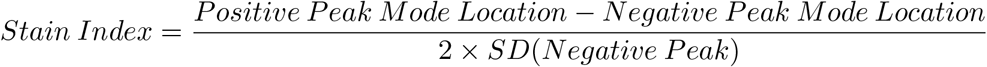

To extend the stain index to capture separation in more diverse data distribution patterns beyond bimodal expression, such as multiple peaks, shoulder peaks or heavy right tail (Supplementary Fig. 1C), we designed the stain quality score as follows:

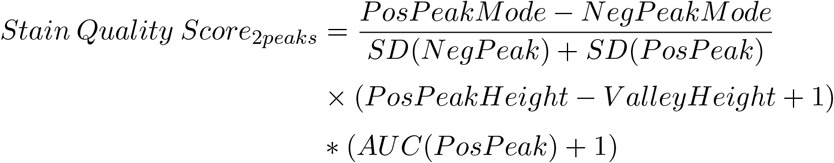

*AUC*(*PosPeak*) means the area under the curve of the positive peak in the corresponding density distribution. Therefore, the stain quality for protein markers with two peaks is positively correlated with the peak mode distance, the sharpness of the positive peak and the proportion of the positive population, and negatively correlated with the total standard deviation in the negative and positive populations.

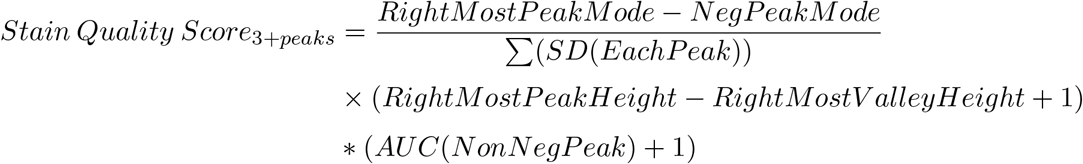

For protein markers with three or more peaks, the stain quality score is positively correlated with the landmark distance between the right-most peak and the negative peak, the sharpness of the most positive peak and the proportion of non-negative populations. The score is negatively correlated with the sum of the standard deviation of each peak.

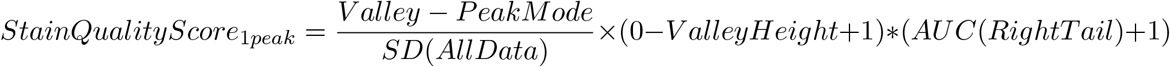

Due to the missing positive peak, for markers with one detected peak, we use the distance between peak and valley as the lower bound of the distance between any positive population and the negative peak mode. We continue to penalize the score for one peak by setting the *PosPeakHeight* to be 0. The area under the curve of the right tail beyond the valley is used to distinguish markers that only have a negative population and markers with a heavy right tail or even shoulder peak. In other words, although the independent positive peak failed to be detected, the positive population is still present.

Stain quality scores are comparable across markers with different peak numbers and generally give higher scores to markers with more peaks. For markers with the same number of identified peaks, better separation of positive and negative populations (longer distance between peak modes) and sharper peaks (lower standard deviation) leads to higher stain quality scores. Markers with two identified peaks score higher than those exhibiting only a shoulder peak. Distributions with only one identified peak, and a heavy right tail will have a lower score, and distributions with only one peak and no right tail will be given the lowest score. Supplementary Fig. 1C provided the diagram illustrating the peak patterns and associated stain quality score order.

### Computational environment for evaluating runtime and memory

Software performance assessments (Supplementary Fig. 4D-E) were conducted on a dedicated server at Fred Hutchinson Cancer Center in terms of running time and memory consumption. The server was equipped with an Intel(R) Xeon(R) Gold 6254 CPU @3.10GHz, featuring 18 cores and 36 threads, and 754GB RAM. For GPU-accelerated tasks, an NVIDIA-SMI GPU with 12GB of VRAM was utilized. The computational environment was hosted on Ubuntu 18.04.6 LTS, with kernel version 4.15.0-213-generic. The software was compiled and run using GCC version 8.3.0 and CUDA toolkit 12.2. Evaluations were performed under minimal system load to ensure consistent and reproducible results.

## Data availability

The raw data used in the paper were downloaded from multiple sources depending on the original studies. Supplementary Table 1 summarized the data source and accession. The corresponding processed data for the 13 public studies were uploaded as demo data to be part of the ADTnorm software repository (https://github.com/yezhengSTAT/ADTnorm/tree/main/data).

## Code availability

ADTnorm package is implemented in R is accompanied by a Python wrapper of the R function. The source codes and detailed instructions for running ADTnorm are publicly available at https://github.com/yezhengSTAT/ADTnorm for R package and https://github.com/donnafarberlab/ADTnormPy for the Python wrapper.

---

**Supplementary Table 1:**
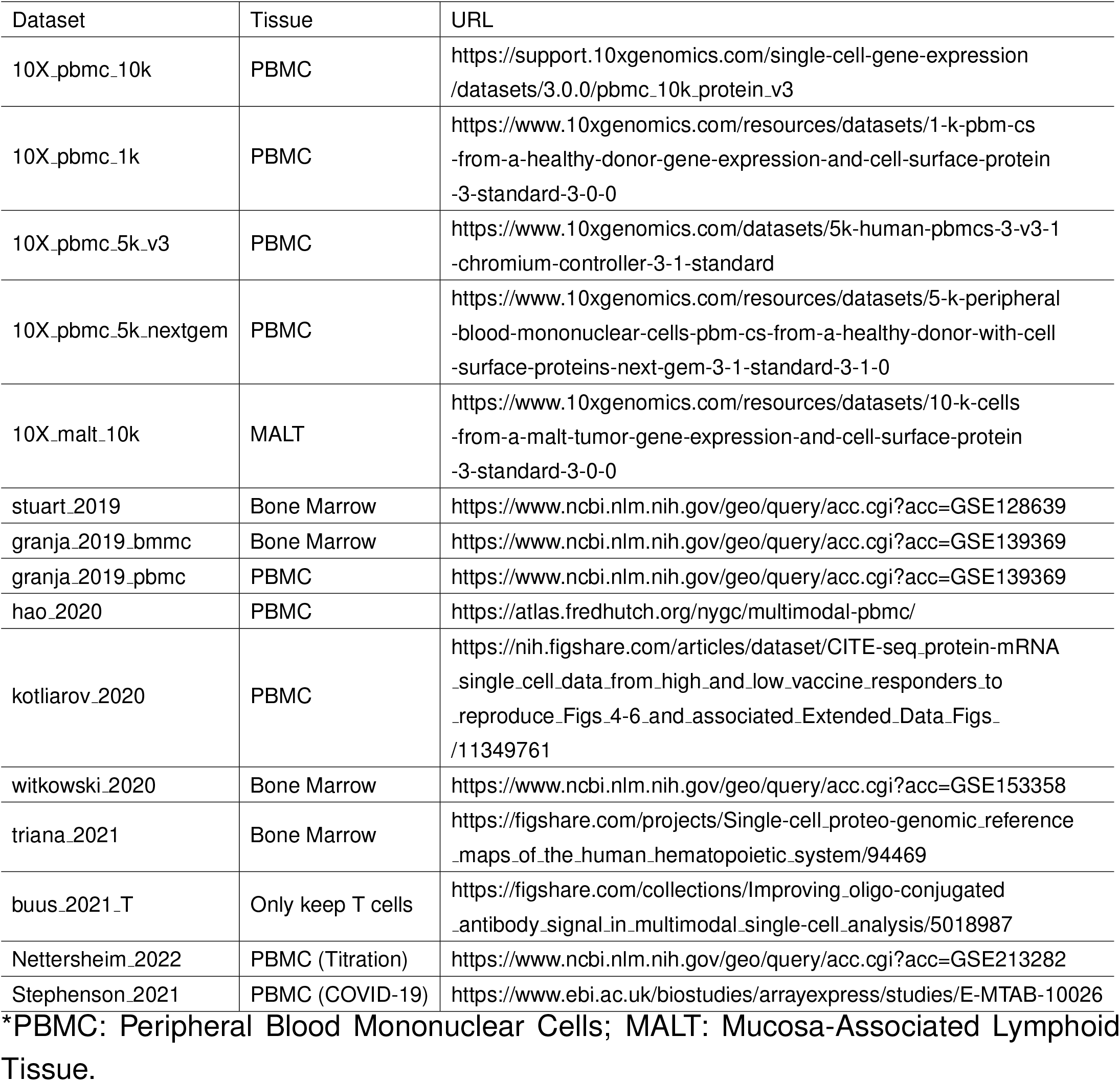
Public CITE-seq data summary.

**Supplementary Table 2:** Manual gating strategy.

| Cell Type | Gating Strategy |
| --- | --- |
| B | CD3<sup>-</sup>CD19<sup>+</sup> |
| CD4 T | CD3<sup>+</sup>CD19<sup>-</sup>CD4<sup>+</sup>CD8<sup>-</sup> |
| CD8 T | CD3<sup>+</sup>CD19<sup>-</sup>CD4<sup>-</sup>CD8<sup>+</sup> |
| DC | CD3<sup>-</sup>CD19<sup>-</sup>CD20<sup>-</sup>CD14<sup>-</sup>HLA-DR<sup>+</sup>CD56<sup>-</sup>CD16<sup>-</sup> |
| NK | CD3<sup>-</sup>CD19<sup>-</sup>CD20<sup>-</sup>CD14<sup>-</sup>HLA-DR<sup>-</sup>CD56<sup>+</sup> |
| Monocytes | CD3<sup>-</sup>CD19<sup>-</sup>CD20<sup>-</sup>CD14<sup>+</sup> |
| naïve B | CD3<sup>-</sup>CD19<sup>+</sup>CD27<sup>-</sup> |
| memory B | CD3<sup>-</sup>CD19<sup>+</sup>CD27<sup>+</sup> |
| naive CD4 | CD3<sup>+</sup>CD19<sup>-</sup>CD4<sup>+</sup>CD8<sup>-</sup>CD25<sup>-</sup>CD45RA<sup>+</sup>CD45RO<sup>-</sup> |
| memory CD4 | CD3<sup>+</sup>CD19<sup>-</sup>CD4<sup>+</sup>CD8<sup>-</sup>CD25<sup>-</sup>CD45RA<sup>-</sup>CD45RO<sup>+</sup> |
| Treg | CD3<sup>+</sup>CD19<sup>-</sup>CD4<sup>+</sup>CD8<sup>-</sup>CD25<sup>+</sup>CD127<sup>-</sup> |
| naive CD8 | CD3<sup>+</sup>CD19<sup>-</sup>CD4<sup>-</sup>CD8<sup>+</sup>CD45RA<sup>+</sup>CD45RO<sup>-</sup> |
| memory CD8 | CD3<sup>+</sup>CD19<sup>-</sup>CD4<sup>-</sup>CD8<sup>+</sup>CD45RA<sup>-</sup>CD45RO<sup>+</sup> |
| plasmacytoid DC | CD3<sup>-</sup>CD19<sup>-</sup>CD20<sup>-</sup>CD14<sup>-</sup>HLA-DR<sup>+</sup>CD56<sup>-</sup>CD16<sup>-</sup>CD123<sup>+</sup>CD11c<sup>-</sup> |
| myeloid DC | CD3<sup>-</sup>CD19<sup>-</sup>CD20<sup>-</sup>CD14<sup>-</sup>HLA-DR<sup>+</sup>CD56<sup>-</sup>CD16<sup>-</sup>CD123<sup>-</sup>CD11c<sup>+</sup> |
| CD16<sup>+</sup> NK | CD3<sup>-</sup>CD19<sup>-</sup>CD20<sup>-</sup>CD14<sup>-</sup>HLA-DR<sup>-</sup>CD56<sup>+</sup>CD16<sup>+</sup> |
| CD16<sup>-</sup> NK | CD3<sup>-</sup>CD19<sup>-</sup>CD20<sup>-</sup>CD14<sup>-</sup>HLA-DR<sup>-</sup>CD56<sup>+</sup>CD16<sup>-</sup> |
| classical monocyte | CD3<sup>-</sup>CD19<sup>-</sup>CD20<sup>-</sup>CD14<sup>+</sup>CD16<sup>-</sup> |
| intermediate monocyte | CD3<sup>-</sup>CD19<sup>-</sup>CD20<sup>-</sup>CD14<sup>+</sup>CD16<sup>+</sup> |
| non-classical CD16<sup>+</sup> monocyte | CD3<sup>-</sup>CD19<sup>-</sup>CD20<sup>-</sup>CD14<sup>-</sup>HLA-DR<sup>+</sup>CD56<sup>-</sup>CD16<sup>+</sup> |

