## Supplementary Figures for "ADTnorm: Robust Integration of Single-cell Protein Measurement across CITE-seq Datasets"

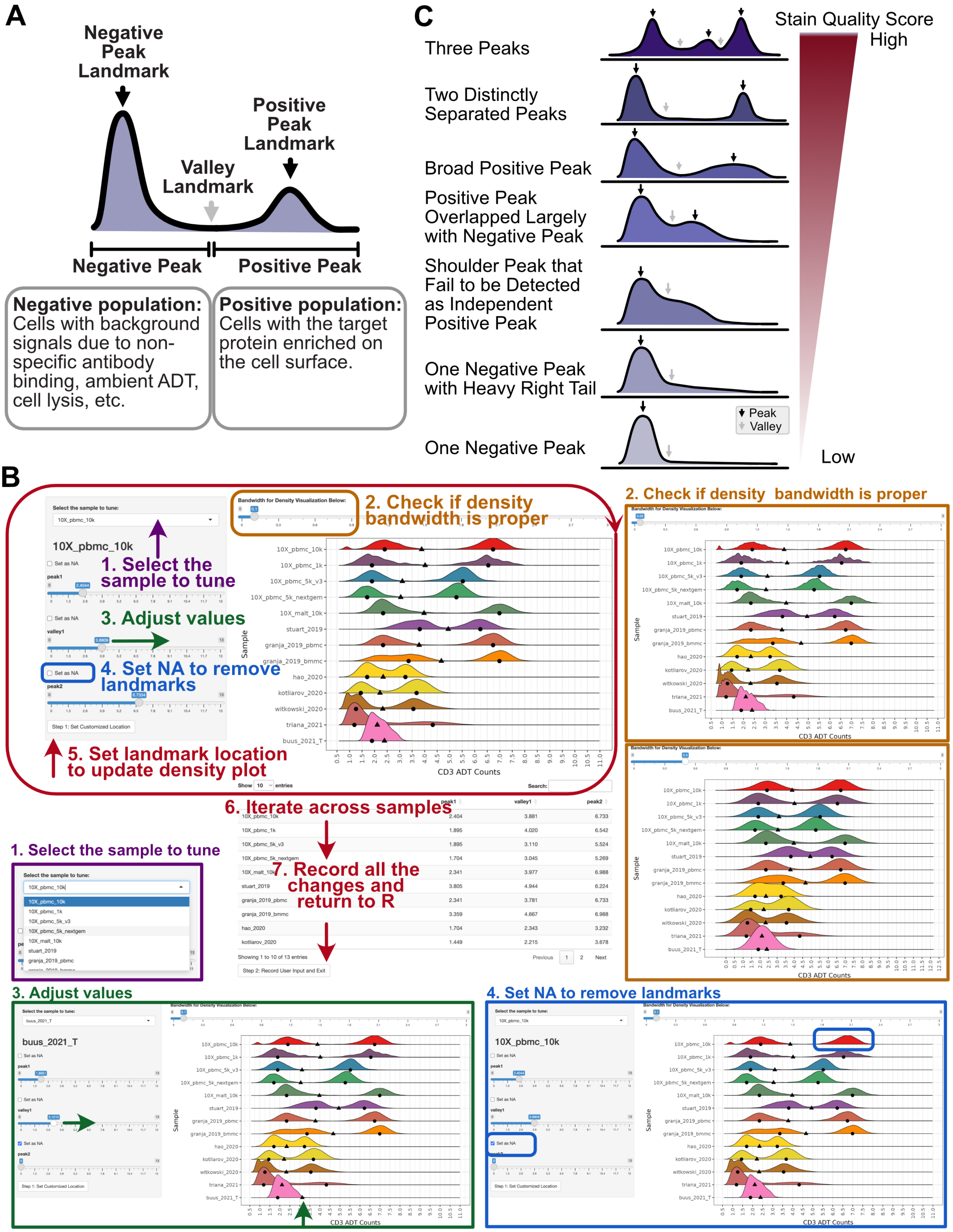


Supplementary Figure 1 Protein landmark definitions and the R shiny interac-

**tive graphical user interface to manually adjust landmark locations. A.** Negative and positive peak landmarks are the modes of negative and positive populations. Valley landmark is the local minimal between negative peak and positive peak. The negative peak indicates the population of cells that do not have the target protein enriched on the cell surface. Instead, those cells only have the background signals due to non-specific antibody binding, ambient ADT, etc. Positive peak captures cells with the target protein enriched on the cell surface. **B.** Peak and valley landmark locations can be manually tuned using R shiny interactive graphical user interface following the procedures: 1. Use the drop-down menu to select the sample to tune. 2. Adjust the density bandwidth to better visualize the ADT count density distributions. 3. Adjust the peak or valley values or remove the valley or peak by setting it to NA (4). 5. Update the landmark locations on the density plot for checking. 6. Tune all the samples that need further adjustment. 7. Finish the process by recording all the changes and return to the R terminal for downstream normalization processing. **C.** Peak separation pattern and the corresponding stain quality score order.

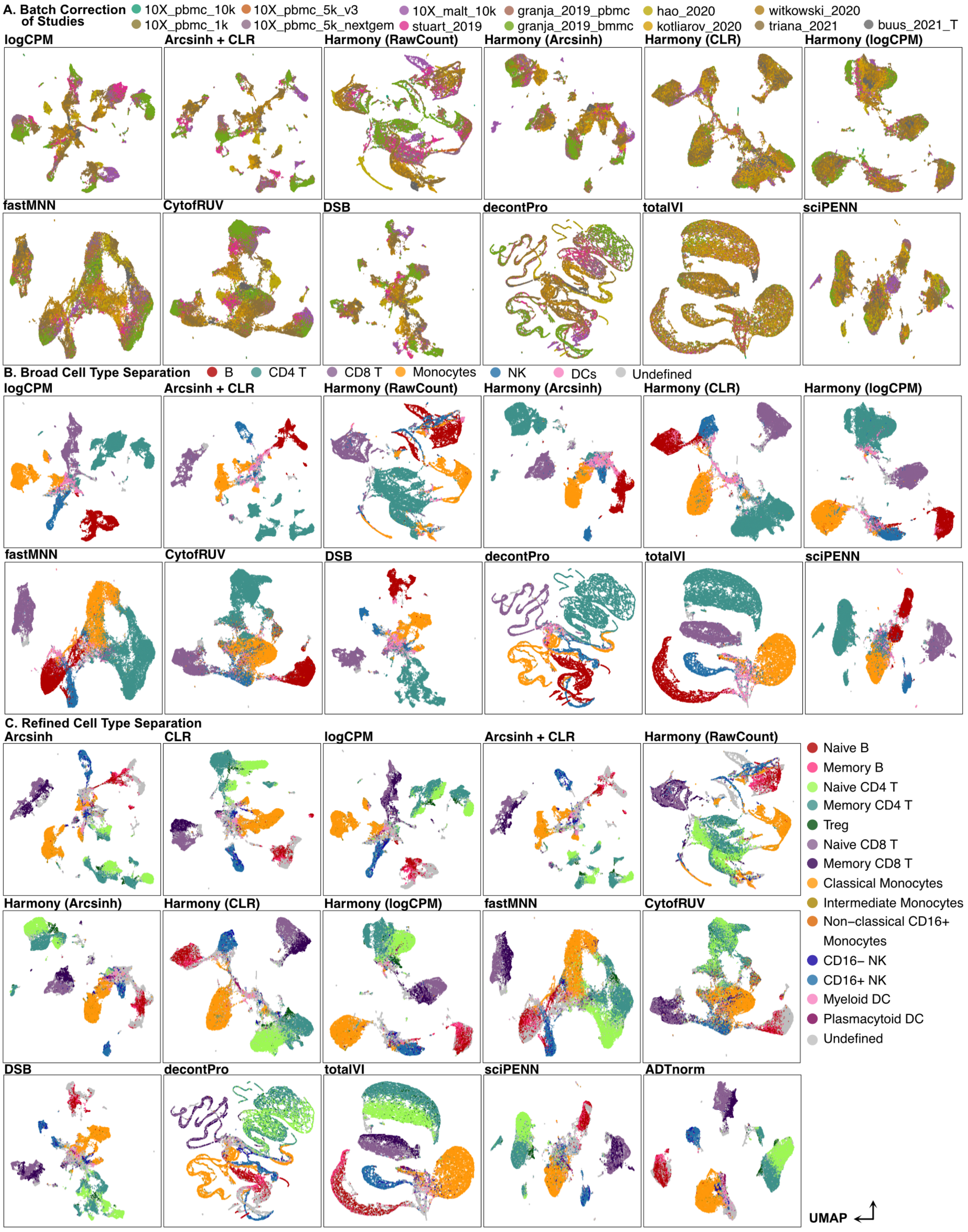
A. Batch Correction of Studies
10X_pbmc_10k
10X_pbmc_5k_v3
10X_malt_10k
granja_2019_pbmc
hao_2020
witkowski_2020
10X_pbmc_1k
10X_pbmc_5k_nextgem
stuart_2019
granja_2019_bmmc
kotliarov_2020
triana_2021
buus_2021_T
logCPM
Arcsinh + CLR
Harmony (RawCount)
Harmony (Arcsinh)
Harmony (CLR)
Harmony (logCPM)
fastMNN
CytofRUV
DSB
decontPro
totalVI
sciPENN
B. Broad Cell Type Separation
B
CD4 T
CD8 T
Monocytes
NK
DCs
Undefined
logCPM
Arcsinh + CLR
Harmony (RawCount)
Harmony (Arcsinh)
Harmony (CLR)
Harmony (logCPM)
fastMNN
CytofRUV
DSB
decontPro
totalVI
sciPENN
C. Refined Cell Type Separation
Arcsinh
CLR
logCPM
Arcsinh + CLR
Harmony (RawCount)
Harmony (Arcsinh)
Harmony (CLR)
Harmony (logCPM)
fastMNN
CytofRUV
DSB
decontPro
totalVI
sciPENN
ADTnorm
Naive B
Memory B
Naive CD4 T
Memory CD4 T
Treg
Naive CD8 T
Memory CD8 T
Classical Monocytes
Intermediate Monocytes
Non−classical CD16+ Monocytes
CD16− NK
CD16+ NK
Myeloid DC
Plasmacytoid DC
Undefined
UMAP

**Supplementary Figure 2 UMAP comparison of normalization methods using each study as one batch.** UMAP embeddings were generated after normalization of 9 shared ADT markers were used across all 13 studies to investigate the batch effect removal (**A**), broad cell type separation (**B**) and refined cell type separation (**C**).

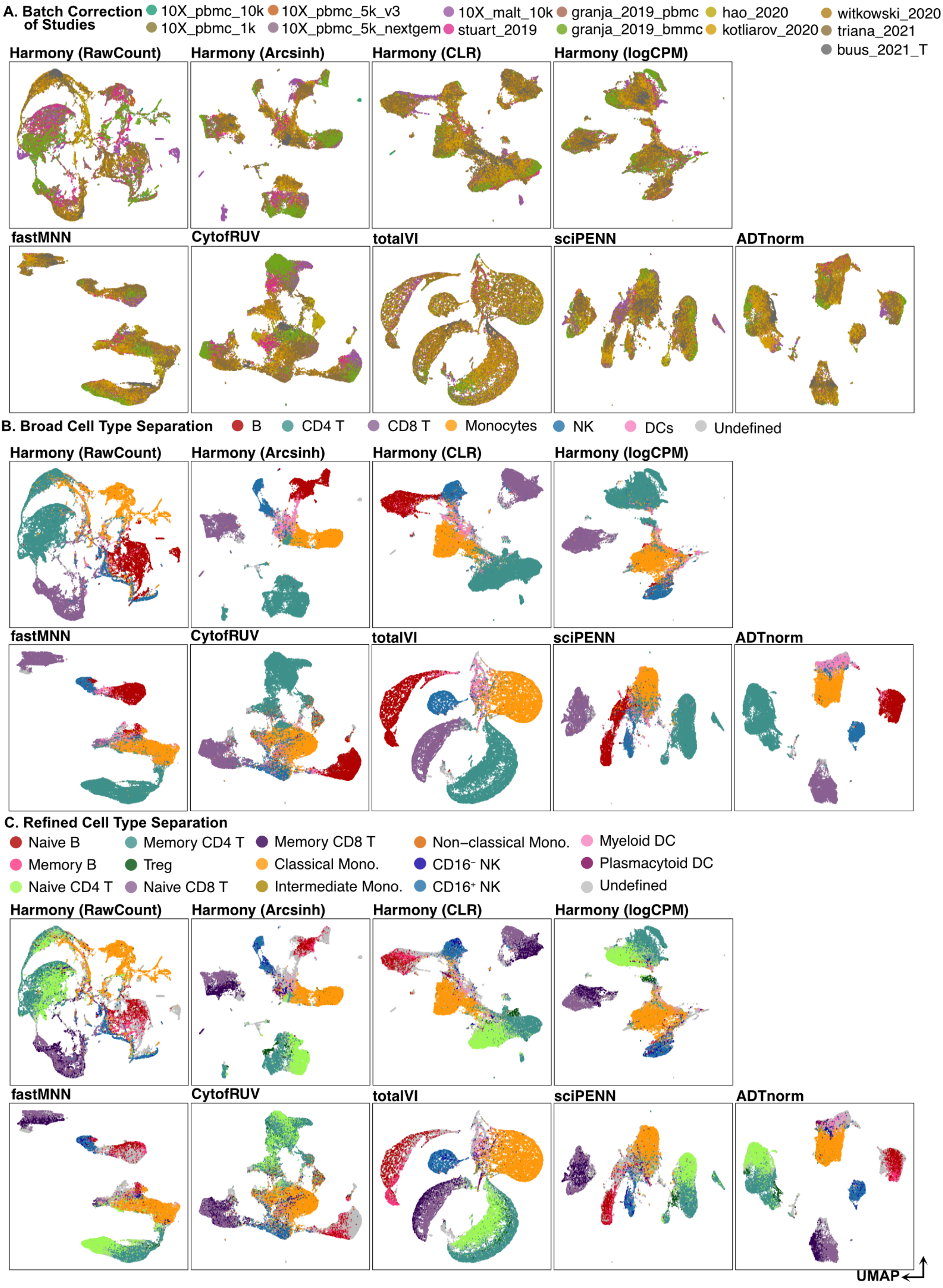

A. Batch Correction of Studies
10X_pbmc_10k
10X_pbmc_1k
10X_pbmc_5k_v3
10X_pbmc_5k_nextgem
10X_malt_10k
stuart_2019
granja_2019_pbmc
granja_2019_bmmc
hao_2020
kotliarov_2020
witkowski_2020
triana_2021
buus_2021_T
Harmony (RawCount)
Harmony (Arcsinh)
Harmony (CLR)
Harmony (logCPM)
fastMNN
CytofRUV
totalVI
sciPENN
ADTnorm
B. Broad Cell Type Separation
B
CD4 T
CD8 T
Monocytes
NK
DCs
Undefined
C. Refined Cell Type Separation
Naive B
Memory B
Naive CD4 T
Memory CD4 T
Treg
Naive CD8 T
Memory CD8 T
Classical Mono.
Intermediate Mono.
Non−classical Mono.
CD16⁻ NK
CD16⁺ NK
Myeloid DC
Plasmacytoid DC
Undefined
UMAP

**Supplementary Figure 3 UMAP comparison of normalization methods using each sample as one batch.** Each sample (i.e., donor or patient) from the 13 studies is considered as one batch. UMAP embeddings were generated after normalization of 9 shared ADT markers to investigate the batch effect removal (**A**), broad cell type separation (**B**), and refined cell type separation (**C**). Methods that lead to the same normalization results as ones using the study as batch key are omitted.

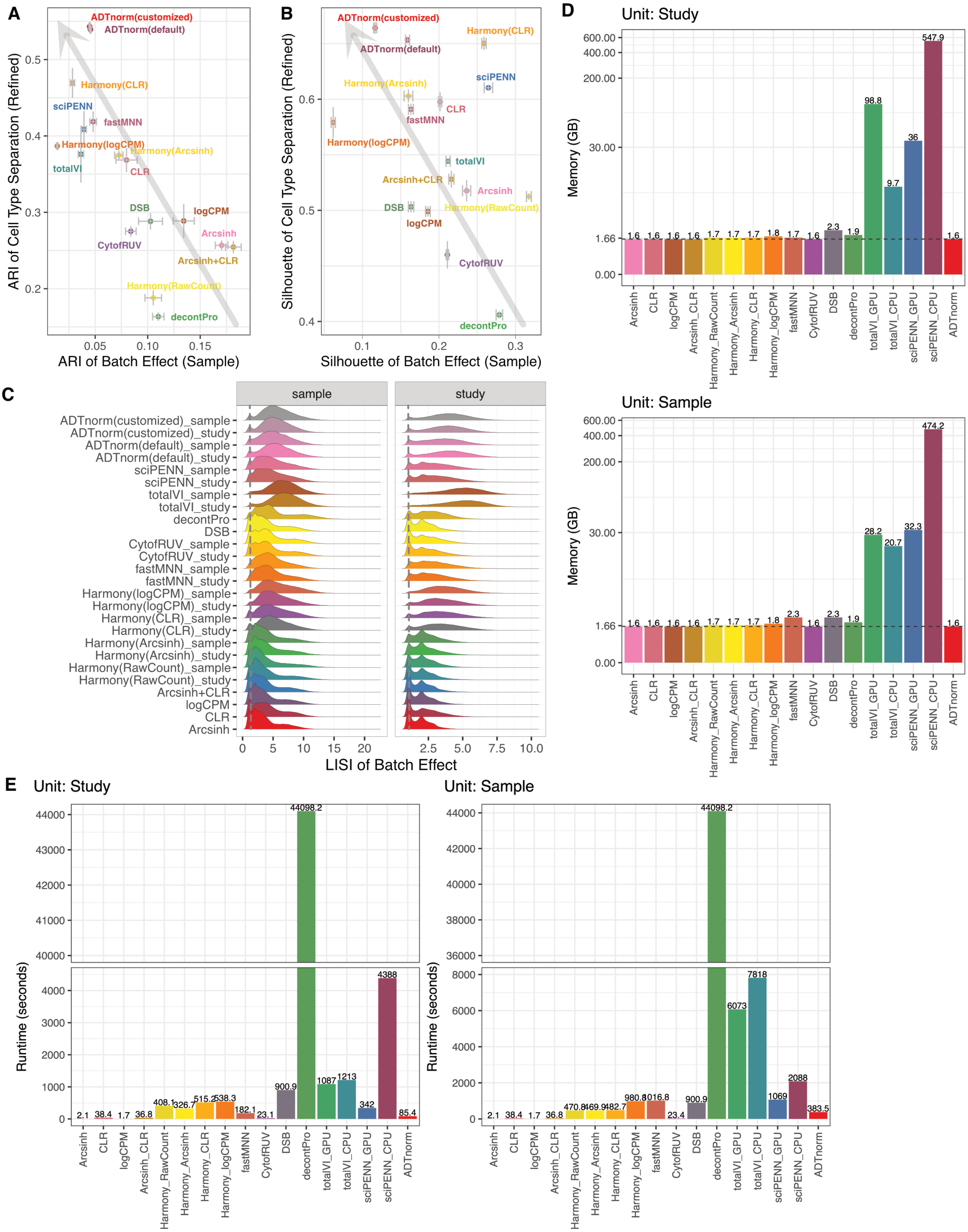


**Supplementary Figure 4  Evaluation of the batch correction and cell type sepa-**

**ration performance.** Silhouette score (**A**) and adjusted rand index (ARI, **B**) are used to evaluate the batch correction and cell type separation performance of ADTnorm and 14 other methods in comparison. Each sample per study is considered as a batch, and the cell type is annotated in a refined manner. Methods that lead to fewer batch effects and better cell type separation are located around the upper-right corner, which is indicated by the grey arrows. **C.** Local Inverse Simpson's Index (LISI) is used to quantify the local batch integration performance. The dashed line is x = 1. High enrichment around x = 1 represents that, within a local neighborhood, there are only cells from the same batches, indicating strong batch effects in the data. Larger LISI means that more batches are present within a local neighborhood, hence showing a better mixture of different batches. Each row corresponds to a method and each column corresponds to the way batch is defined, considering each sample as a batch (left) or each study as a batch (right). **D.** Memory usage and **E.** running time evaluation across methods to normalize and integrate CITE-seq data from 13 public studies. Memory and running time are evaluated under two scenarios: normalize using each study as one batch or normalize using each sample as one batch.

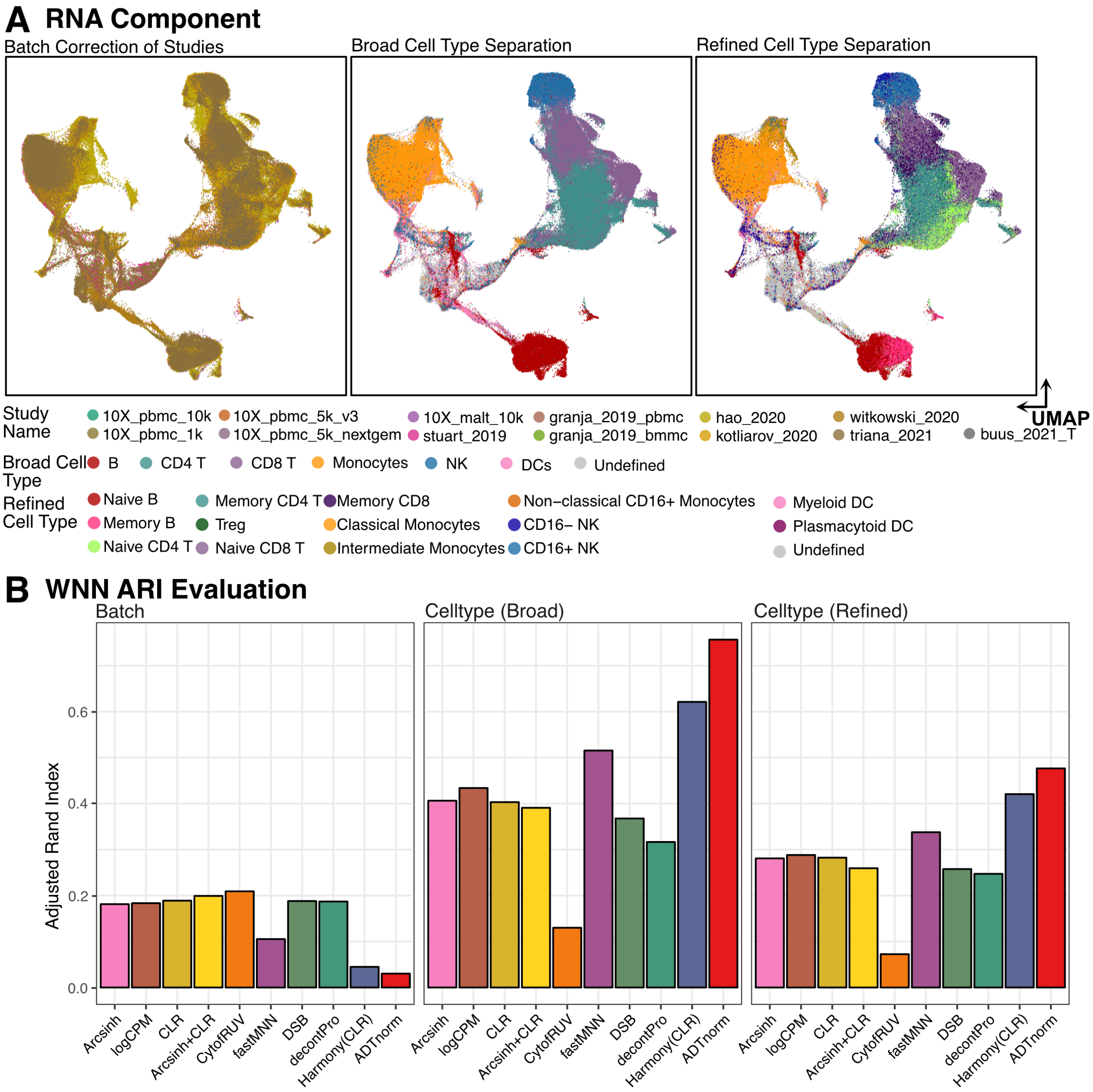


**Supplementary Figure 5 RNA component integration and the evaluation after weighted nearest neighbor integration of RNA and protein. A.** RNA component of CITE-seq data is integrated by reciprocal PCA algorithm across studies. Colors correspond to batch, disease status, and cell types, as indicated in the panel title. **B.** Batch correction and cell type separation at broad and refined levels are evaluated by adjusted rand index. Comparison is across methods that normalize ADT counts without borrowing the gene expression information.

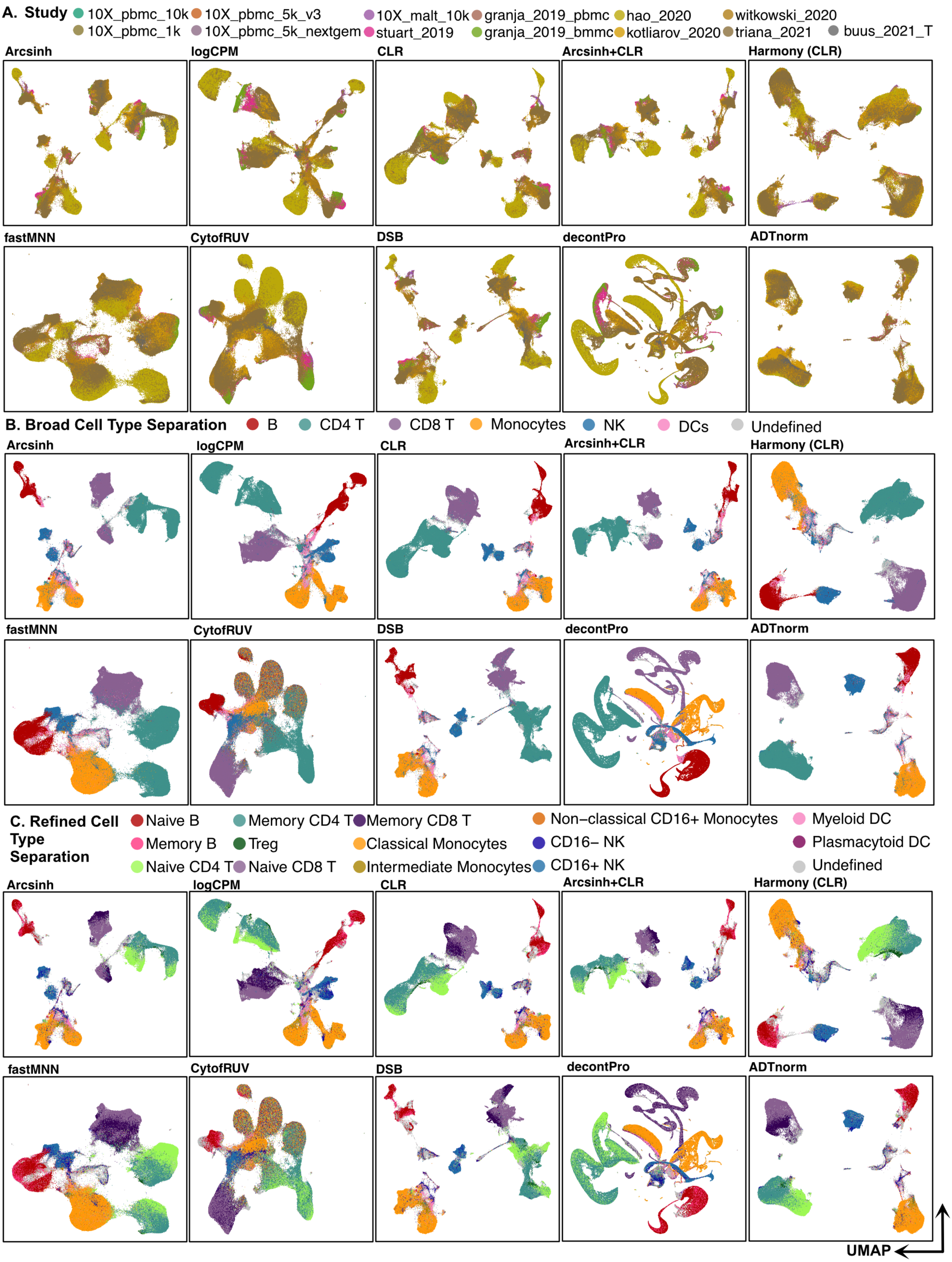

A. Study
10X_pbmc_10k
10X_pbmc_1k
10X_pbmc_5k_v3
10X_pbmc_5k_nextgem
10X_malt_10k
stuart_2019
granja_2019_pbmc
granja_2019_bmmc
hao_2020
kotliarov_2020
witkowski_2020
triana_2021
buus_2021_T
Arcsinh
logCPM
CLR
Arcsinh+CLR
Harmony (CLR)
fastMNN
CytofRUV
DSB
decontPro
ADTnorm
B. Broad Cell Type Separation
B
CD4 T
CD8 T
Monocytes
NK
DCs
Undefined
C. Refined Cell Type Separation
Naive B
Memory B
Naive CD4 T
Memory CD4 T
Treg
Naive CD8 T
Memory CD8 T
Classical Monocytes
Intermediate Monocytes
Non−classical CD16+ Monocytes
CD16− NK
CD16+ NK
Myeloid DC
Plasmacytoid DC
Undefined
UMAP

**Supplementary Figure 6 The batch correction after weighted nearest neighbor integration of RNA and normalized ADT counts by ADTnorm and other methods in comparison.** The RNA component is first aggregated by reciprocal PCA across studies, and the protein component is normalized by each method for batch correction using each study as one batch. The weighted nearest neighbor algorithm is used to integrate RNA and protein and UMAPs are colored by study (**A**), broad cell type (**B**) and refined cell type (**C**).

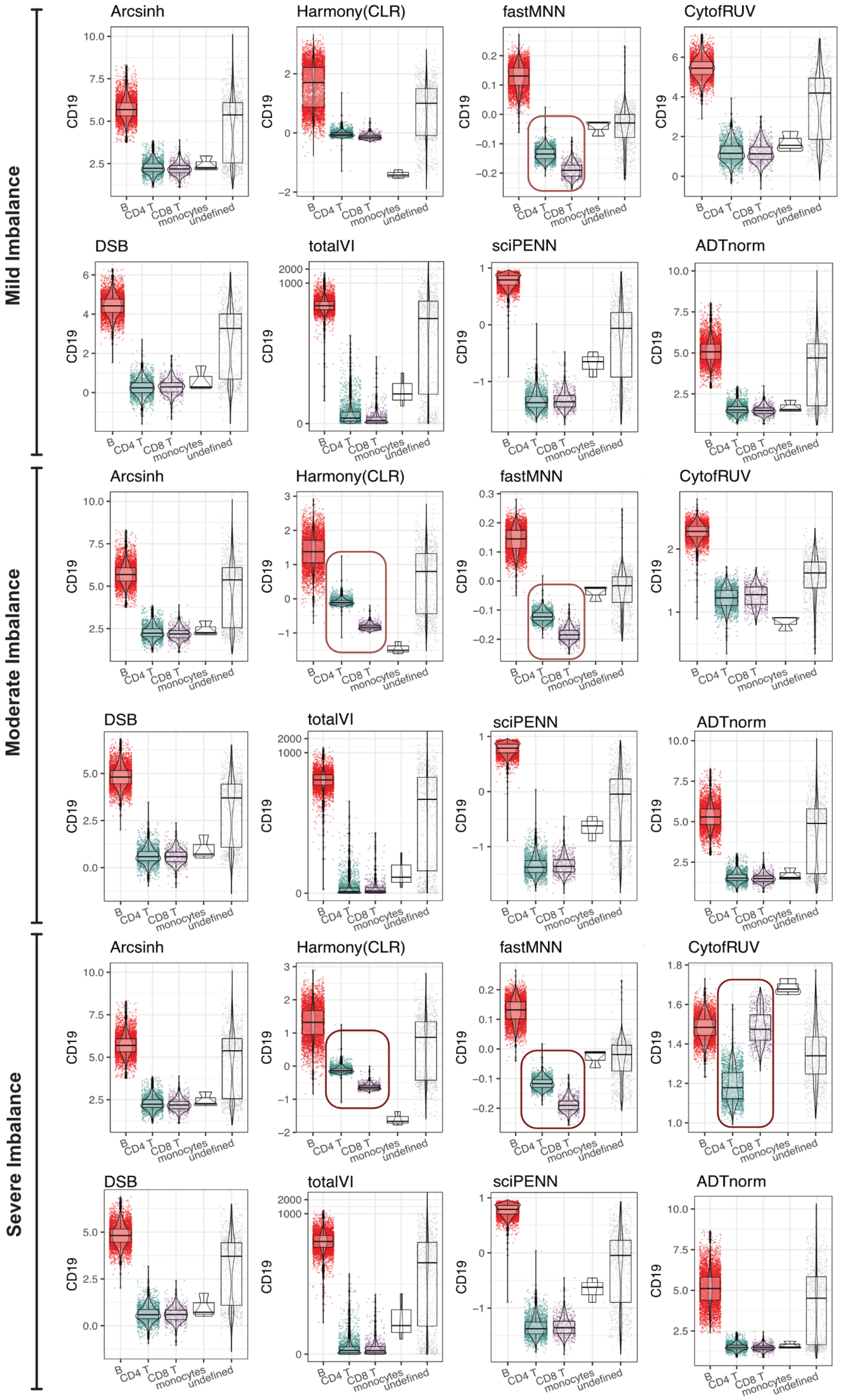


**Supplementary Figure 7 Artifact introduced during the normalization leading to abnormal CD19 values in 10X_malt_10k dataset.** 10X_malt_10k dataset, consisting of one sample, is used to demonstrate the unexpected CD19 normalization values in the CD4 T and CD8 T cells. Three cell-type imbalanced scenarios (Methods) are created to test the robustness of the normalization methods. Abnormal values are highlighted by the red circles.

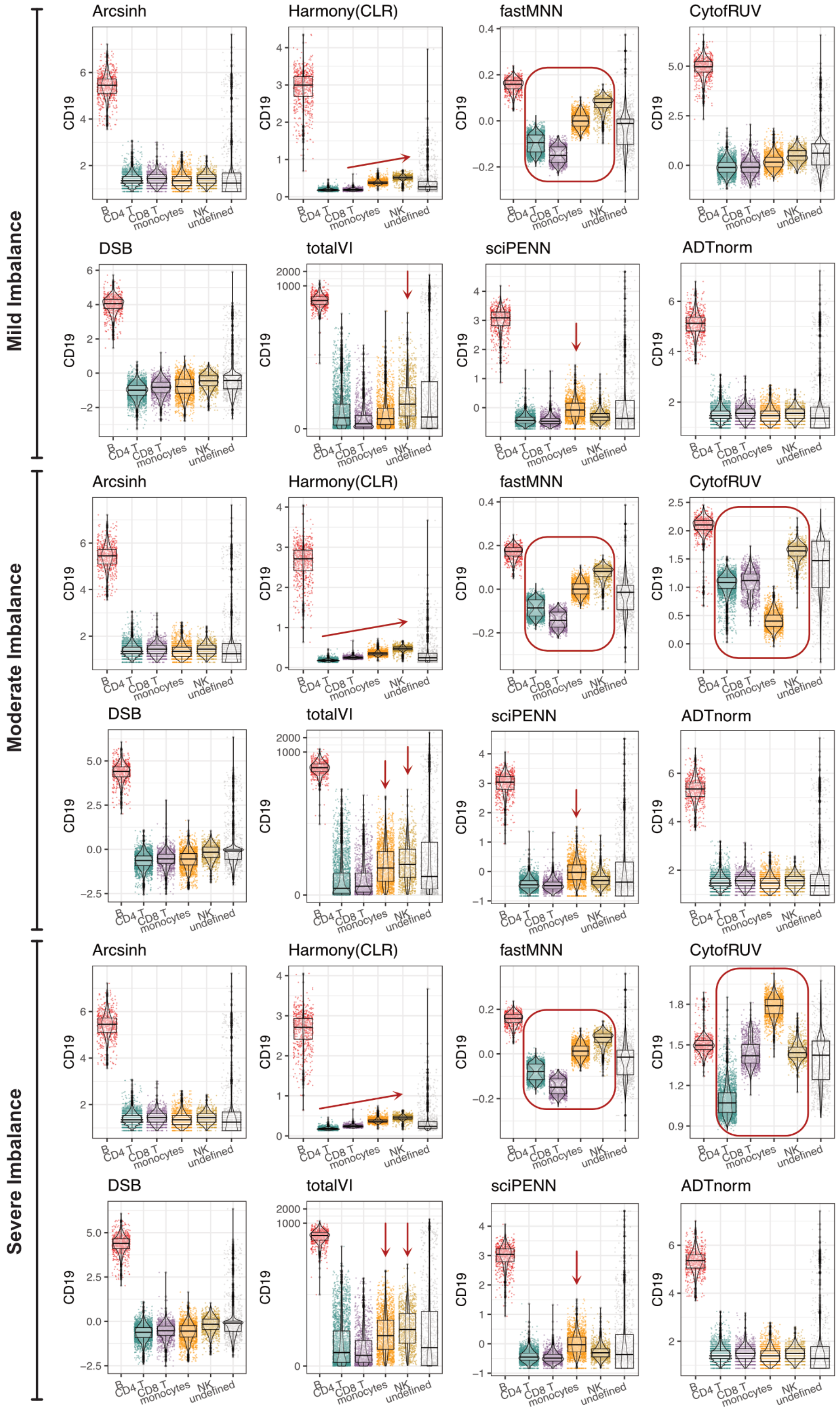


**Supplementary Figure 8 Artifact introduced during the normalization leading to abnormal CD19 values compared across cell types in 10X_pbmc_10k dataset.** 10X_pbmc_10k dataset, consisting of one sample, is used to demonstrate the unexpected CD19 normalization values in the T, Monocytes and NK cells. Three cell-type imbalanced scenarios (Methods) are created to test the robustness of the normalization methods. Abnormal values are highlighted by the red arrows and circles.

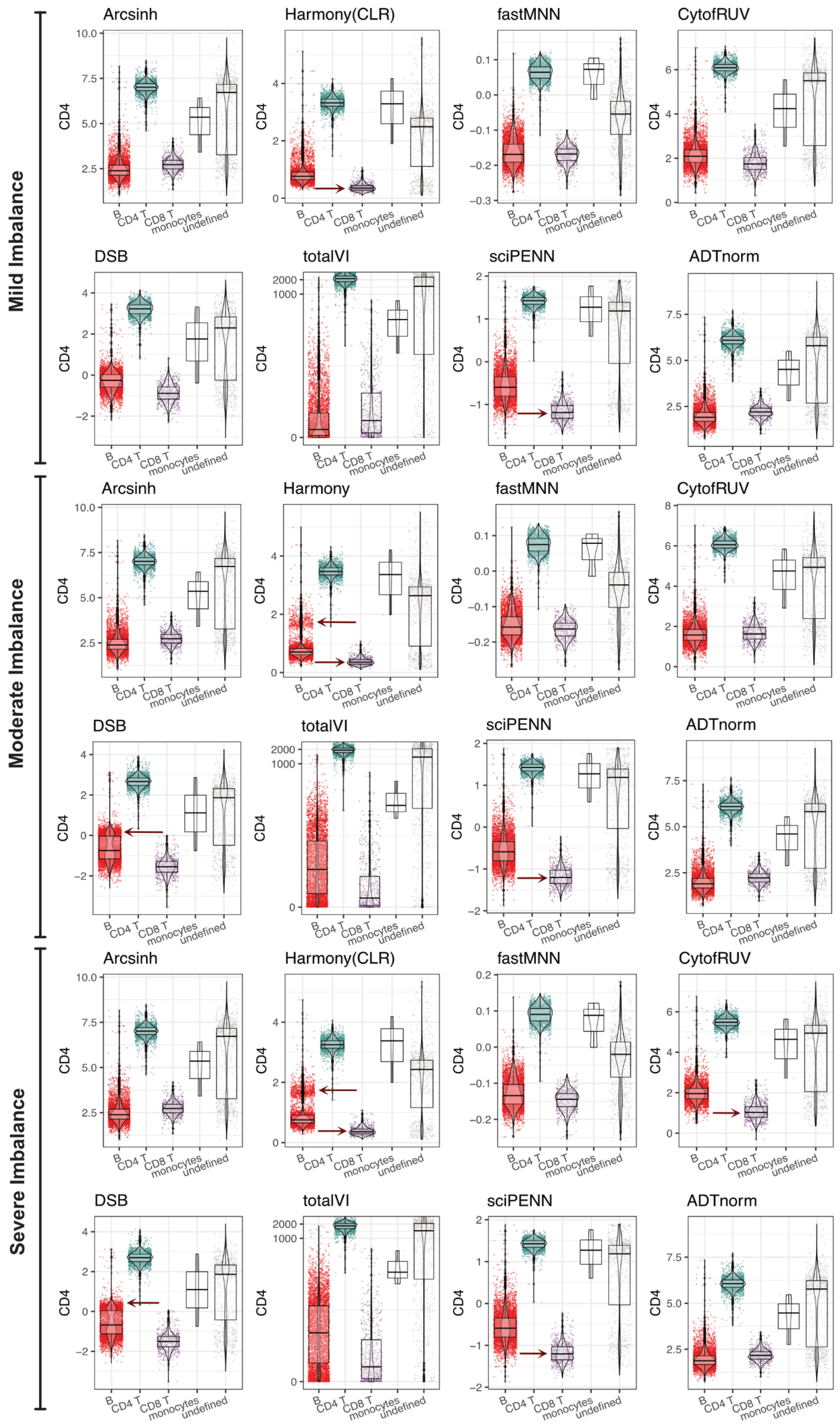


**Supplementary Figure 9 Artifact introduced during the normalization leading to abnormal CD4 values compared across cell types in 10X_malt_10k dataset.** 10X_malt_10k dataset, consisting of one sample, is used to demonstrate the unexpected CD4 normalization values in the B and CD8 T cells. Three cell-type imbalanced scenarios (Methods) are created to test the robustness of the normalization methods. Abnormal values are highlighted by the red arrows.

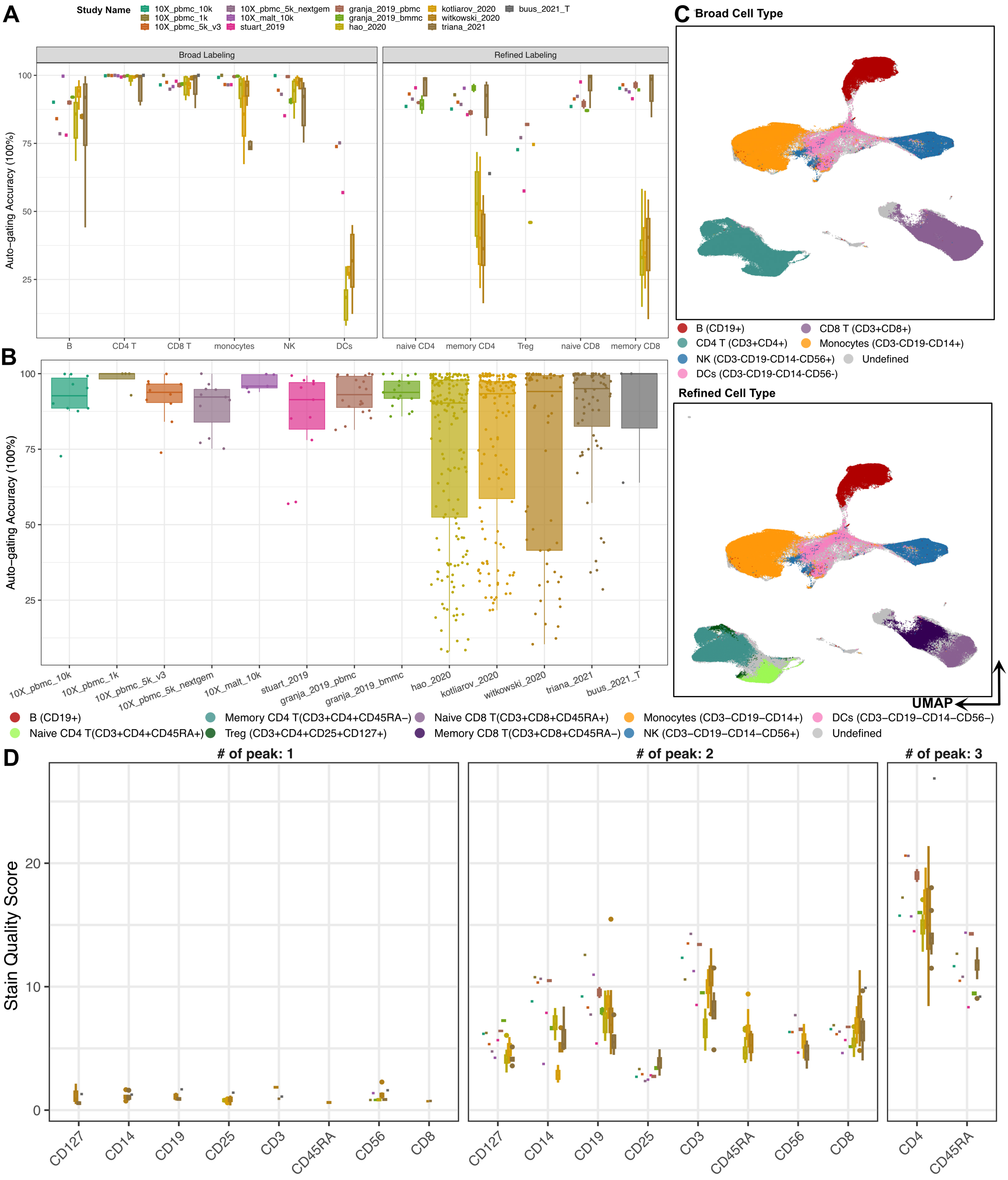


**Supplementary Figure 10 Auto-gating performance on the 13 public datasets.** **A.** Auto-gating accuracy compared to the manual gating cell type annotation by two immunologists. Auto-gating relies on the valley landmarks detected during the ADTnorm normalization. The auto-gating rules to define cell type are specified in the color legend at the bottom of panel B. Color corresponds to the study. **B.** Auto-gating accuracy across different studies. Each point on the boxplot represents a cell type per sample from the corresponding study. **C.** Auto-gating cell types annotation on the UMAP. **D.** Stain quality score (Methods) is used to quantify the positive and negative population separation power. The stain quality score is designed in a way so that protein markers with more peaks will have a higher stain quality score. Within the same number of peaks, markers with better separation between the negative peak and the right most positive peaks, namely longer distances between peak landmarks and sharper peaks, will be granted a higher score.

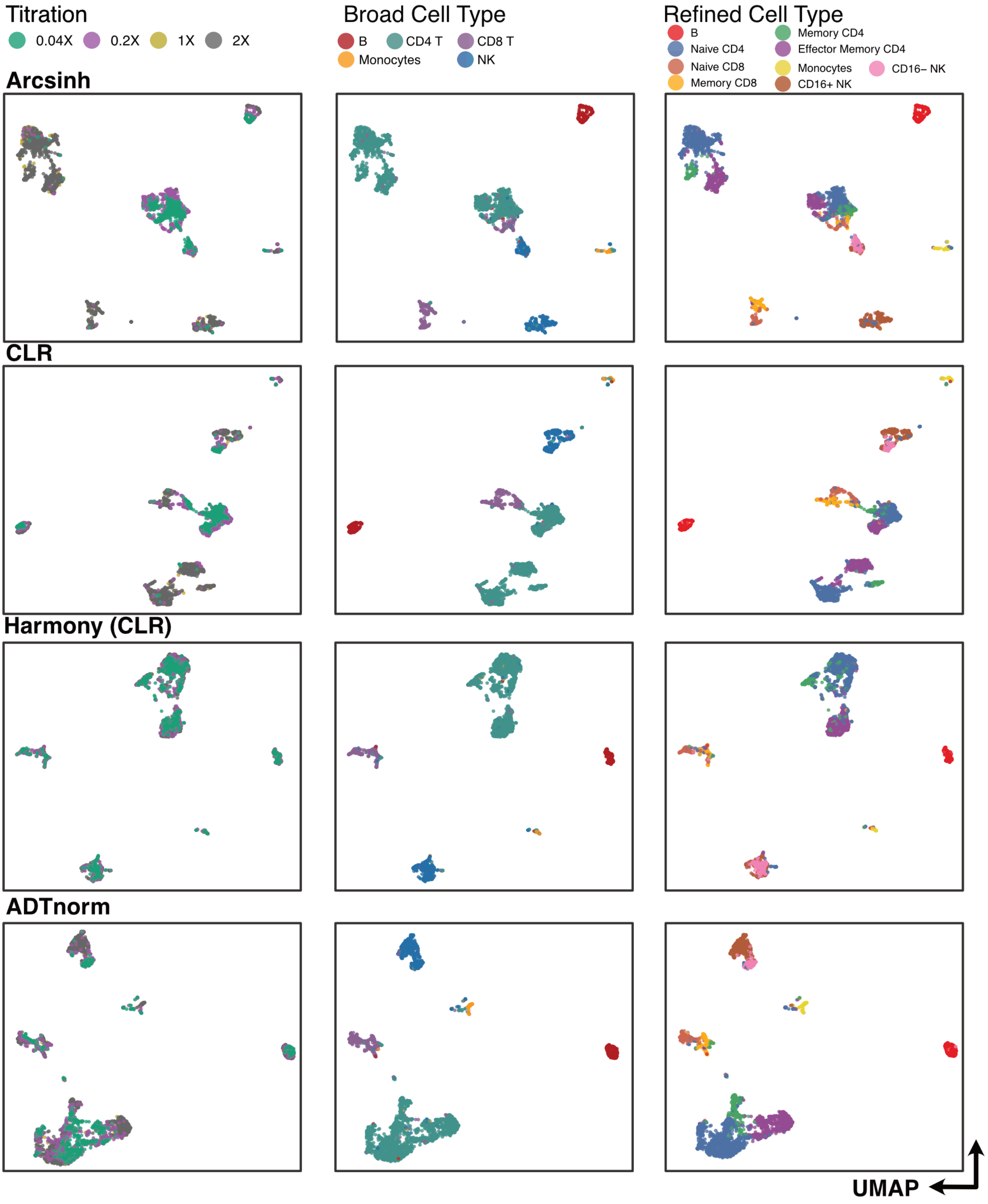


**Supplementary Figure 11 Titration experiments integration across four antibody concentration levels.** UMAP visualizes the integration of four levels of antibody concentrations, colored by the concentration levels, broad and refined cell types. Results from four normalization methods, Arcsinh, CLR, Harmony normalization on CLR transformed counts, and ADTnorm, are compared.

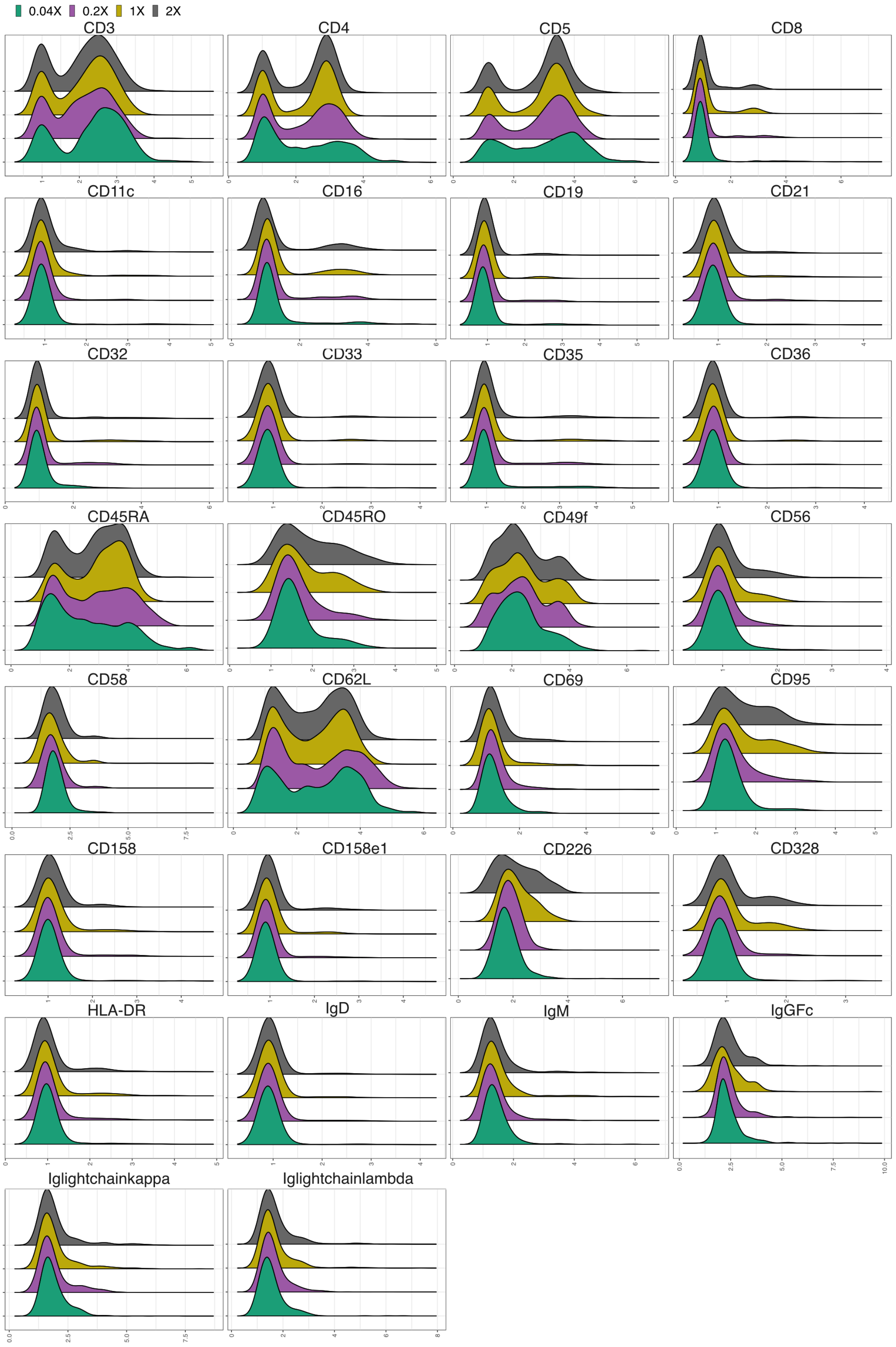


**Supplementary Figure 12 ADTnorm rescued the protein markers stained with low antibody concentration to reinforce the same positive and negative population separation power.**

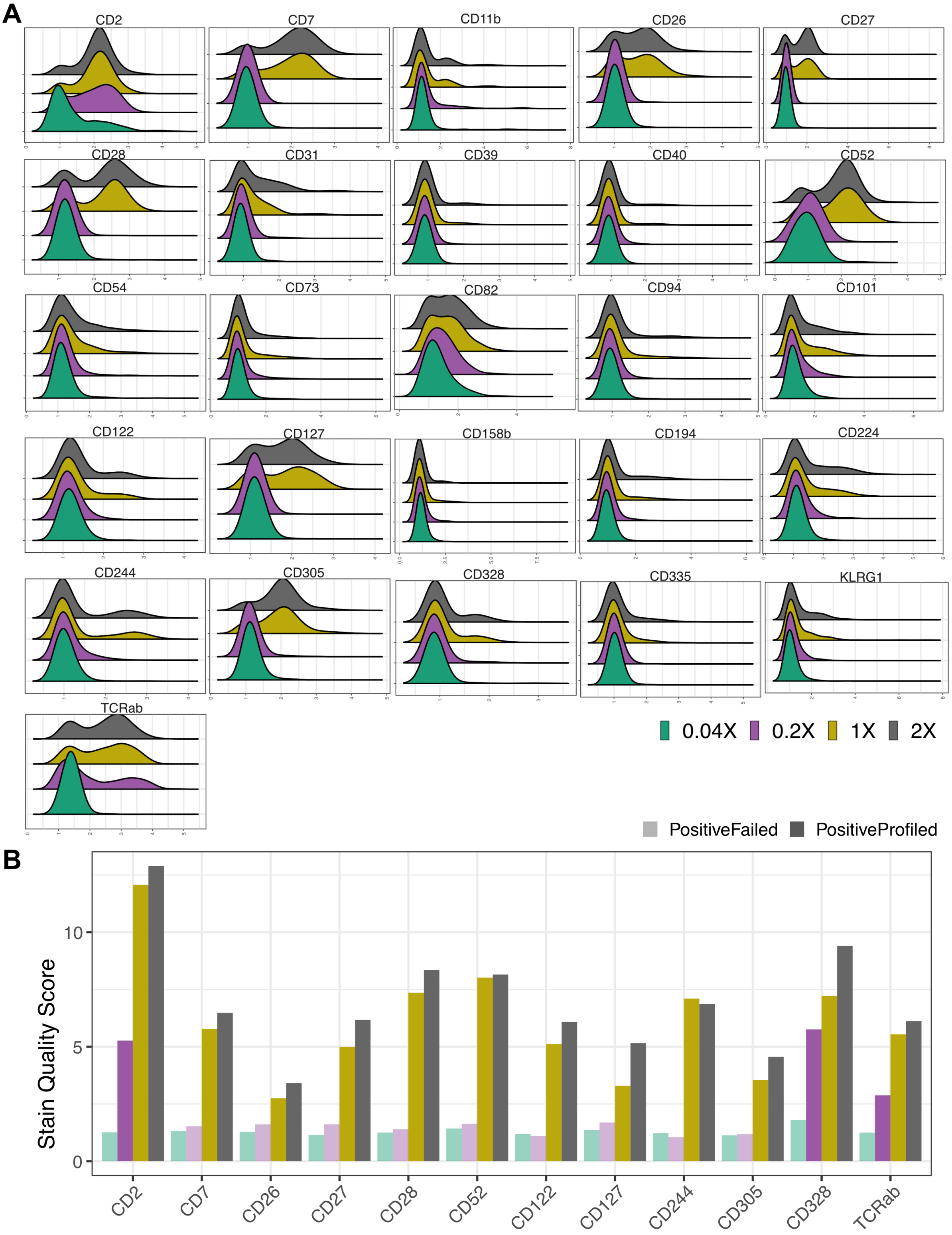


**Supplementary Figure 13 Density distribution and the staining quality of protein markers that failed to profile positive populations. A.** Lower antibody concentration can be insufficient to detect any positive population. Hence, those protein markers cannot be rescued by the ADTnorm normalization. **B.** The stain quality of the ADT marker profiled by lower antibody concentration is insufficient to detect any positive population.

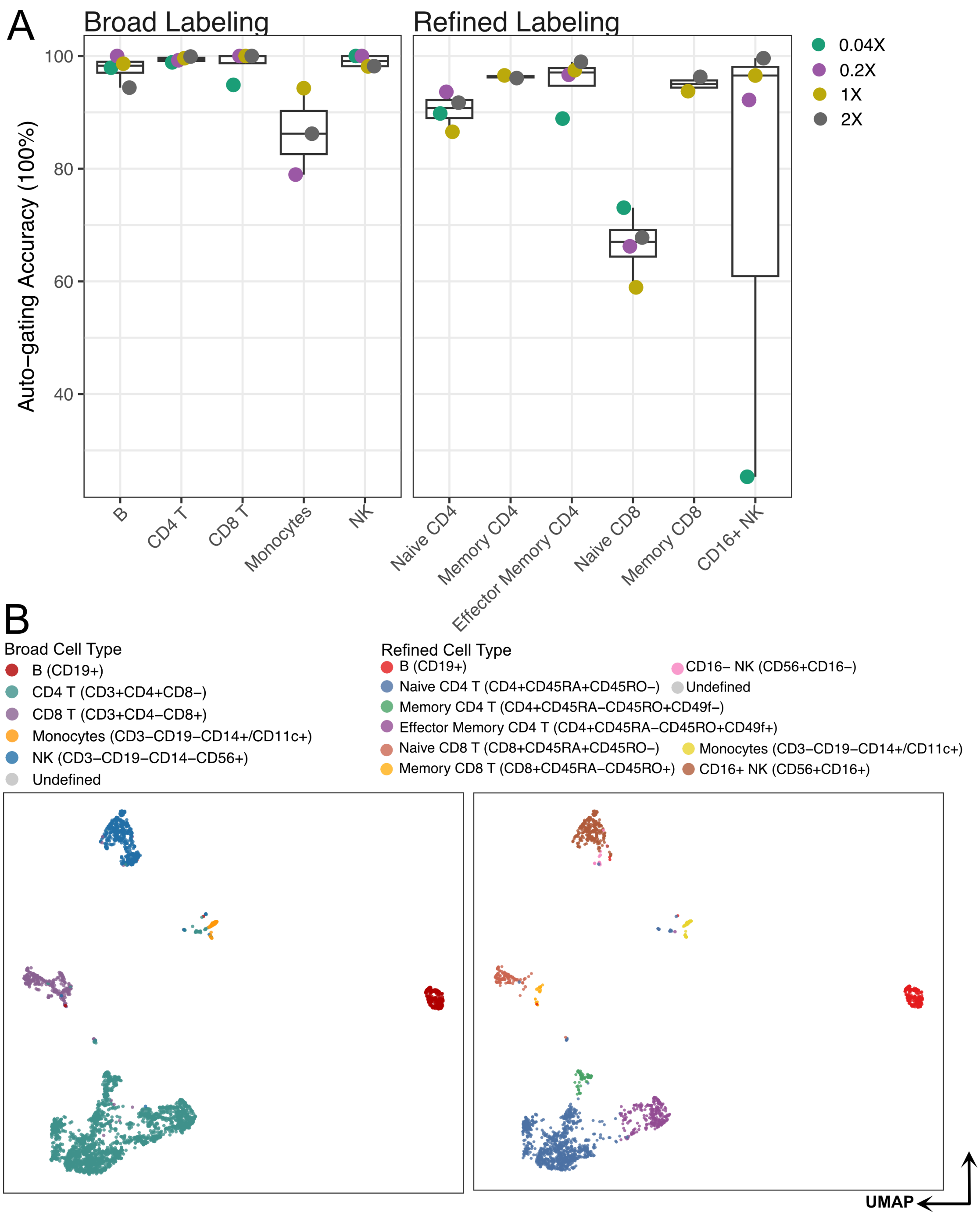


**Supplementary Figure 14 Auto-gating for cell type annotation on the titration experiments. A.** Auto-gating accuracy across broad and refined cell type annotations. 1/25 and/or 1/5 of the commercially recommended antibody concentration can be insufficient for profiling some lineage markers needed for defining certain cell types. Therefore, the dots, indicating samples, can be missing on the corresponding boxplot, such as Monocytes, Memory CD4, Memory CD8, and CD16+ NK cells. **B.** Auto-gating performance on the UMAP visualization of cells from four titration experiments. The gating strategy to define each cell type is specified in the parenthesis.

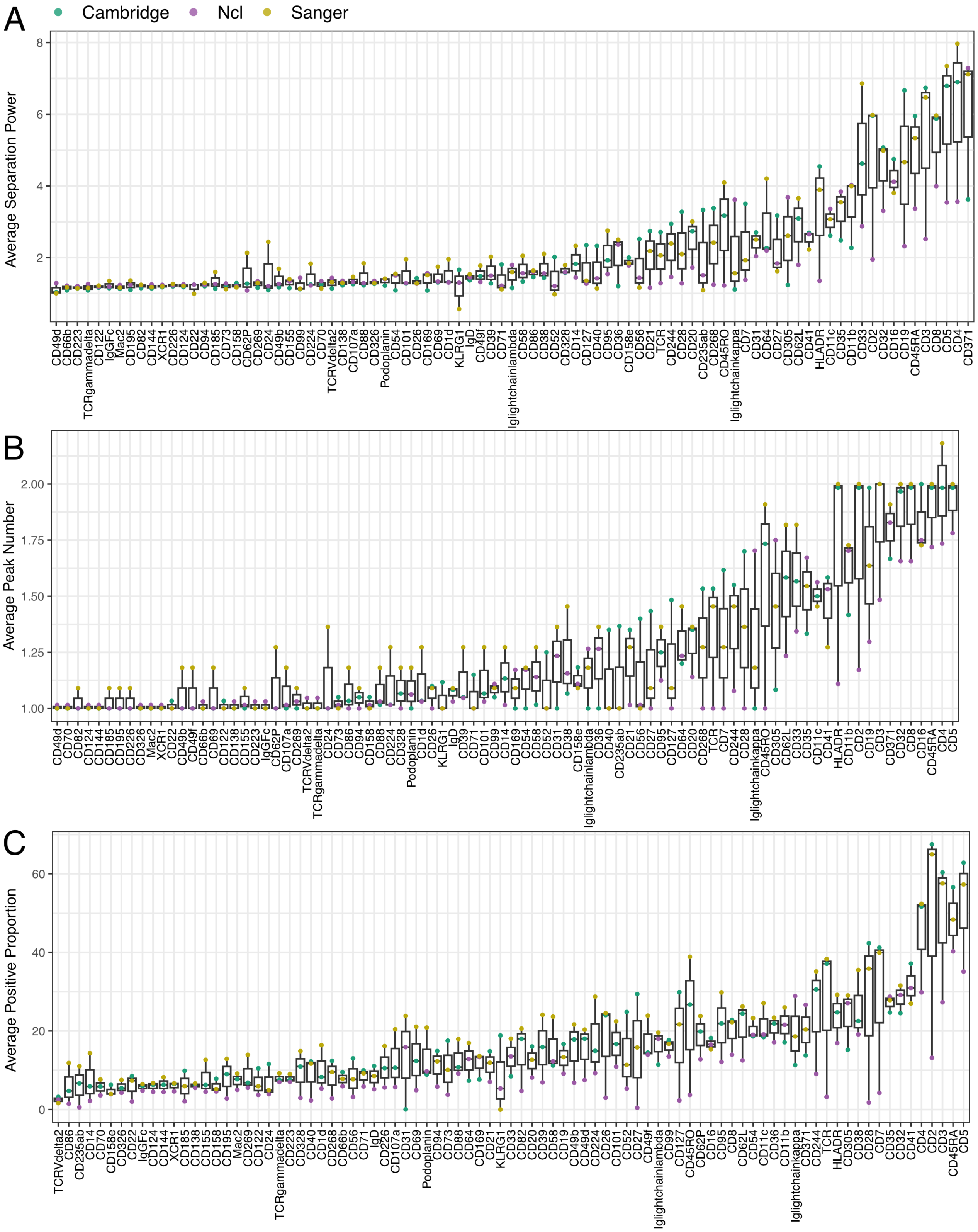


**Supplementary Figure 15 Comparison of protein staining qualities across three research centers for each protein marker.** The average protein marker stain quality (**A**), peak number (**B**) and proportion of positive cells (**C**) are compared across three research institutes for 192 protein markers profiled in the COVID-19 study.

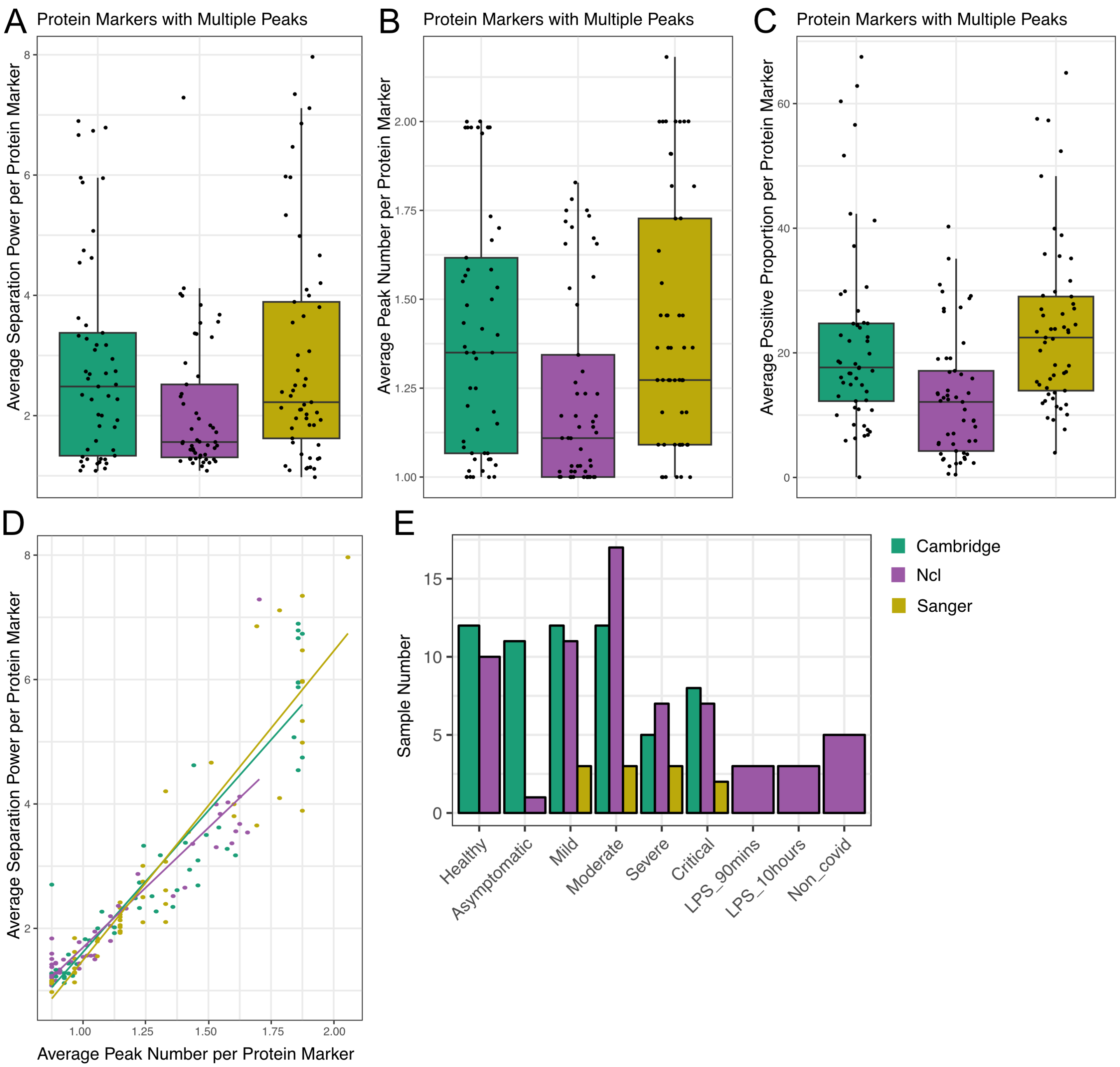


**Supplementary Figure 16 Protein marker stain quality comparison across three research institutes where COVID-19 data were collected. A.** The stain quality score is used to quantify the negative and positive population separation power for each protein marker (i.e., dots on the boxplot) averaged over samples from each research institute (color). **B.** Peak number comparison across three research institutes (color) for each protein marker. **C.** The proportion of positive cells (falling within the positive peak regions) per protein marker is compared across three research institutes (color). The average is taken across samples from the same institute. **D.** The average stain quality scores and the average peak number per protein marker demonstrate a linear relationship. **E.** The sample number from each research institute across different disease symptoms. To avoid bias where samples are from only one center, we excluded LPS and non-COVID samples from the disease-associated marker detection analysis.

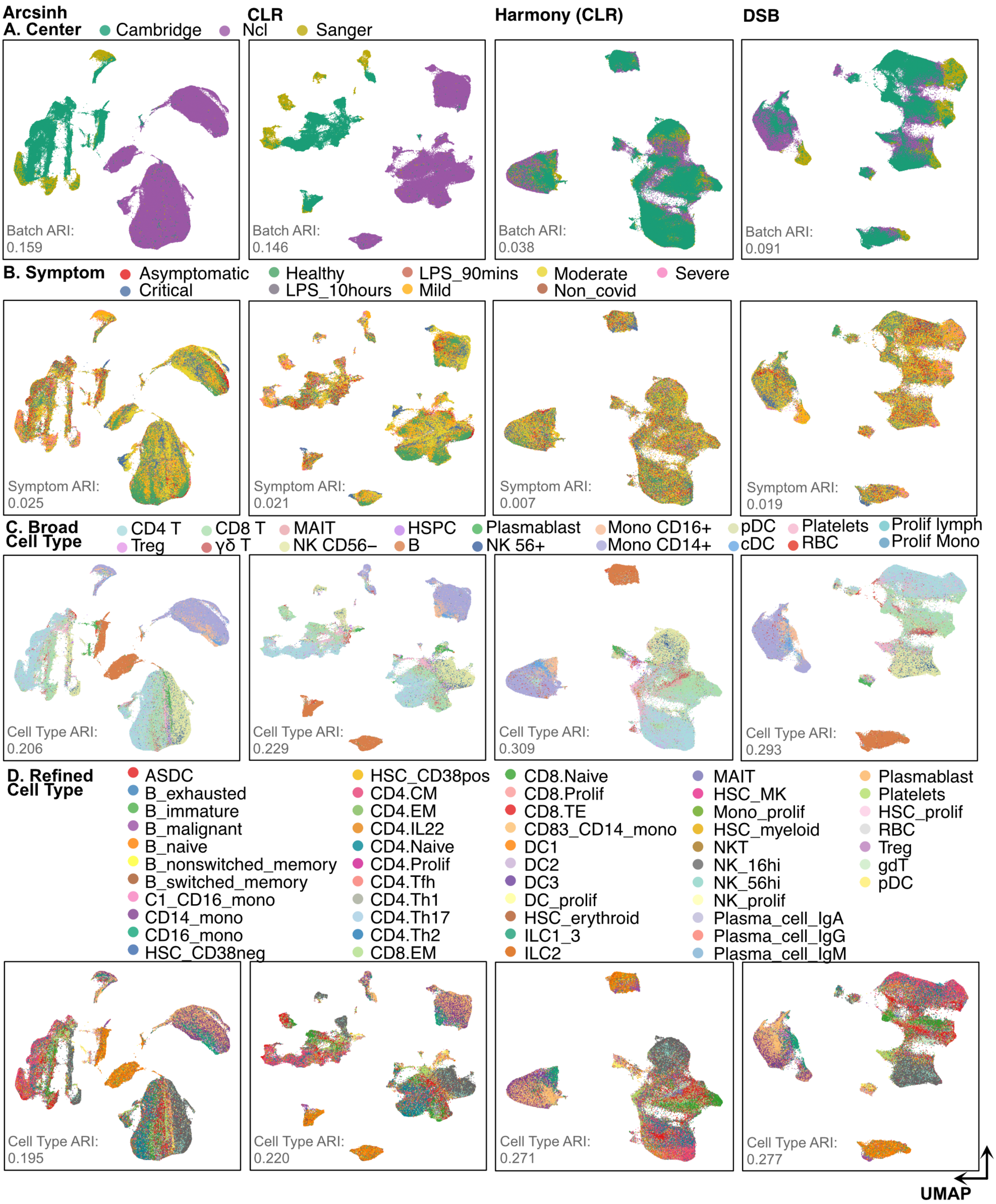


**Supplementary Figure 17 UMAP visualization of Arcsinh, CLR, Harmony and DSB normalization on the ADT counts from the COVID-19 study.** Color corresponds to (row 1) three research institutes where data were collected, (row 2) different disease symptoms, (row 3) broad cell type annotation from the original publication, and (row 4) refined cell type annotation from the original publication.

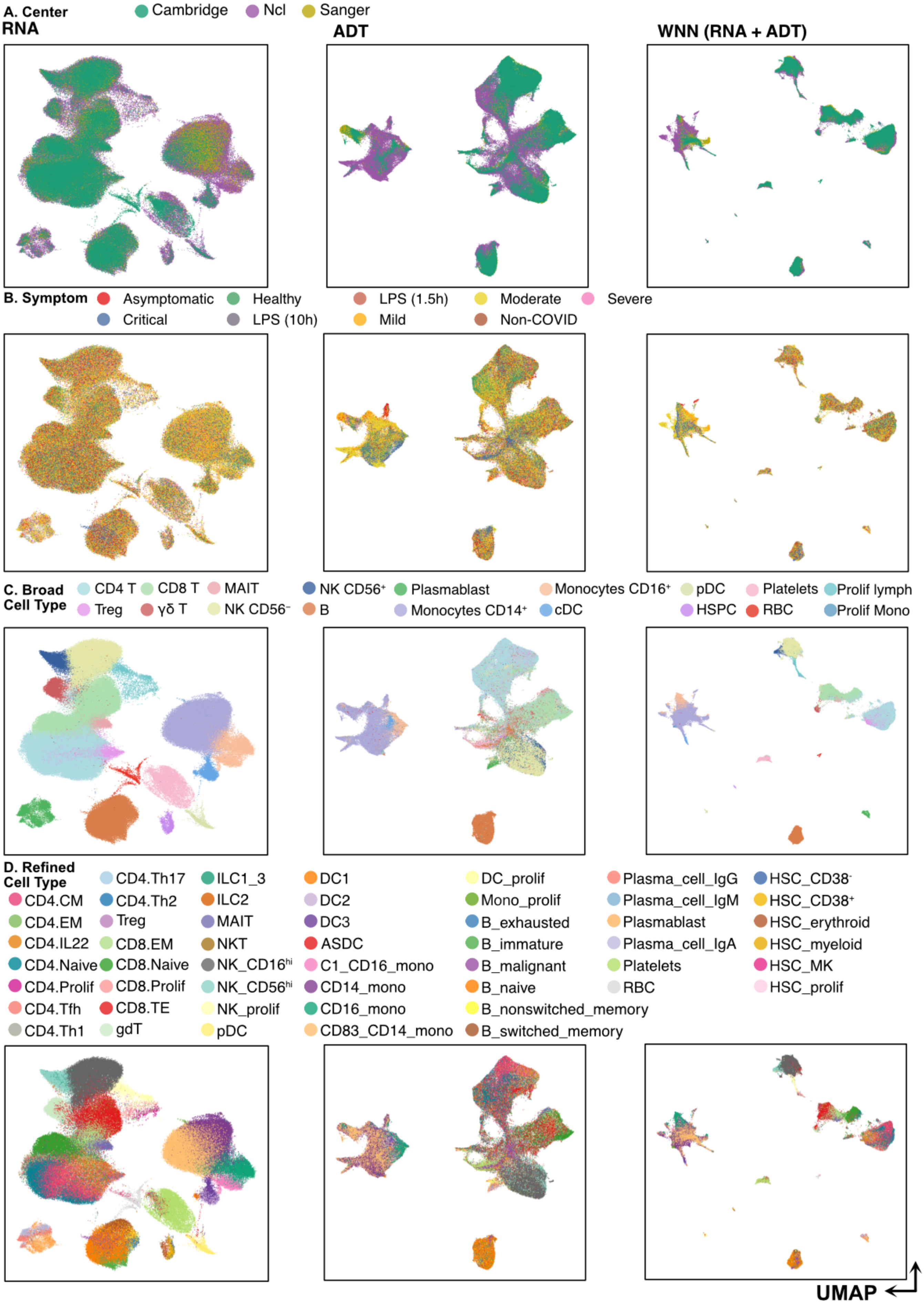
A. Center
Cambridge
Ncl
Sanger
RNA
ADT
WNN (RNA + ADT)
B. Symptom
Asymptomatic
Healthy
LPS (1.5h)
Moderate
Severe
Critical
LPS (10h)
Mild
Non-COVID
C. Broad Cell Type
CD4 T
CD8 T
MAIT
NK CD56+
Plasmablast
Monocytes CD16+
pDC
Platelets
Prolif lymph
Treg
γδ T
NK CD56-
B
Monocytes CD14+
cDC
HSPC
RBC
Prolif Mono
D. Refined Cell Type
CD4.Th17
ILC1_3
DC1
DC_prolif
Plasma_cell_IgG
HSC_CD38-
CD4.CM
CD4.Th2
ILC2
DC2
Mono_prolif
Plasma_cell_IgM
HSC_CD38+
CD4.EM
Treg
MAIT
DC3
B_exhausted
Plasmablast
HSC_erythroid
CD4.IL22
CD8.EM
NKT
ASDC
B_immature
Plasma_cell_IgA
HSC_myeloid
CD4.Naive
CD8.Naive
NK_CD16hi
C1_CD16_mono
B_malignant
Platelets
HSC_MK
CD4.Prolif
CD8.Prolif
NK_CD56hi
CD14_mono
B_naive
RBC
HSC_prolif
CD4.Tfh
CD8.TE
NK_prolif
CD16_mono
B_nonswitched_memory
CD4.Th1
gdT
pDC
CD83_CD14_mono
B_switched_memory
UMAP

**Supplementary Figure 18 Weighted nearest neighbor integration of RPCA aggregated RNA and ADTnorm normalized protein data.** The RNA component is first aggregated by reciprocal PCA (RPCA) across studies, and the protein component is normalized by ADTnorm for batch correction. The weighted nearest neighbor algorithm is used to integrate processed RNA and protein. The color corresponds to: **A.** the three research institutes from which the data were collected (row 1), **B.** disease symptoms (row 2), **C.** broad cell types from the original publication (row 3), and **D.** refined cell types from the original publication (row 4).

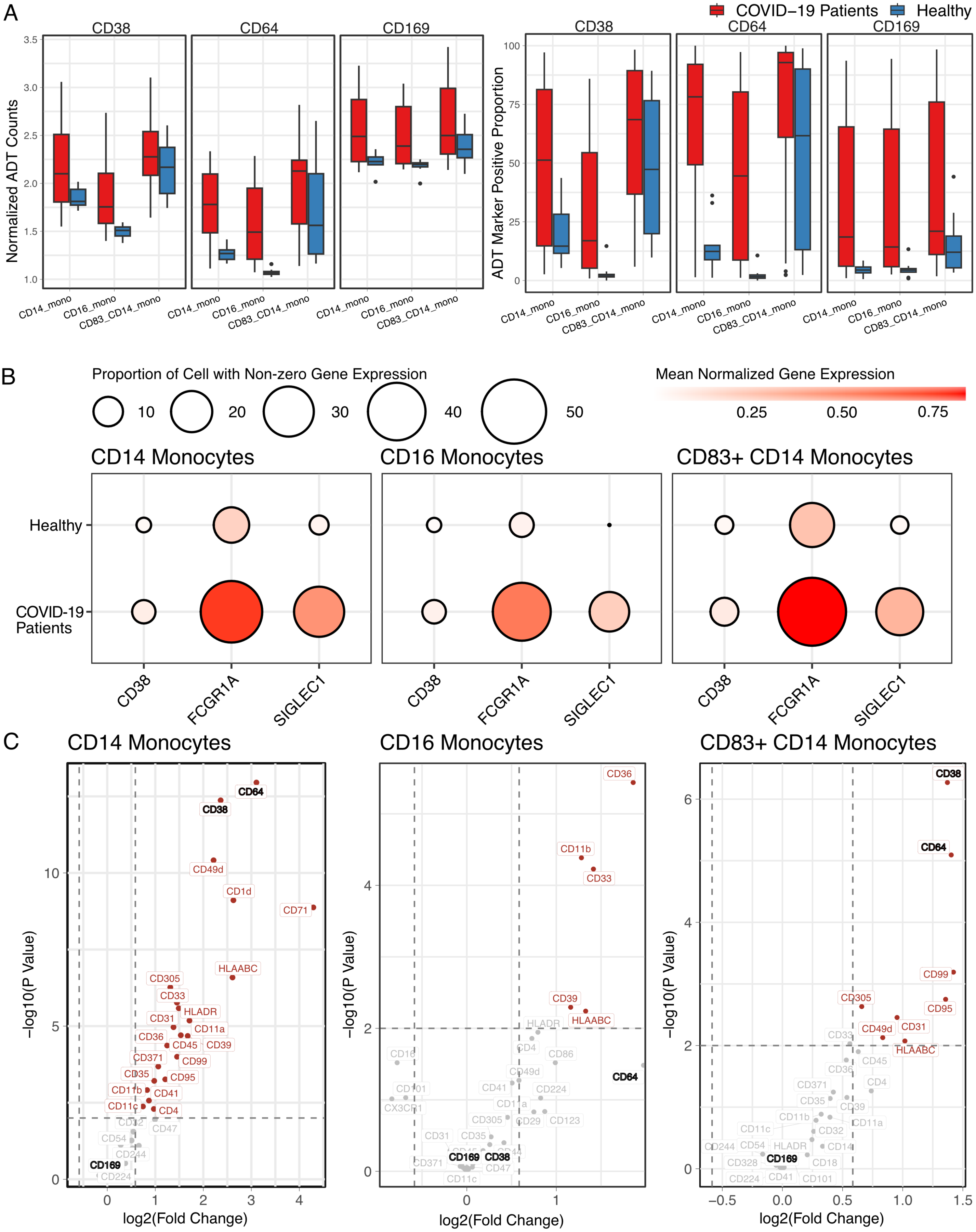


**Supplementary Figure 19 Differential detection compared between healthy donor**

**and COVID-19 patients. A.** ADTnorm normalized ADT counts and the proportion of the positive cells compared between the healthy donors and COVID-19 patients in terms of CD38, CD64 and CD169 in three monocytes subsets. **B.** Comparison of the scRNA-seq normalized expression and the proportion of cells regarding the protein CD38, CD64 and CD169's corresponding genes, i.e., CD38, FCGR1A and SIGLEC1, between healthy and COVID-19 patients across three monocyte subsets. **C.** Volcano plot of differential detection between healthy donors and COVID-19 patients using the positive cells after DSB normalization. The differential detection was done for cells annotated by CD14 Monocytes, CD16 Monocytes and $CD83^+$ CD14 monocytes, respectively.

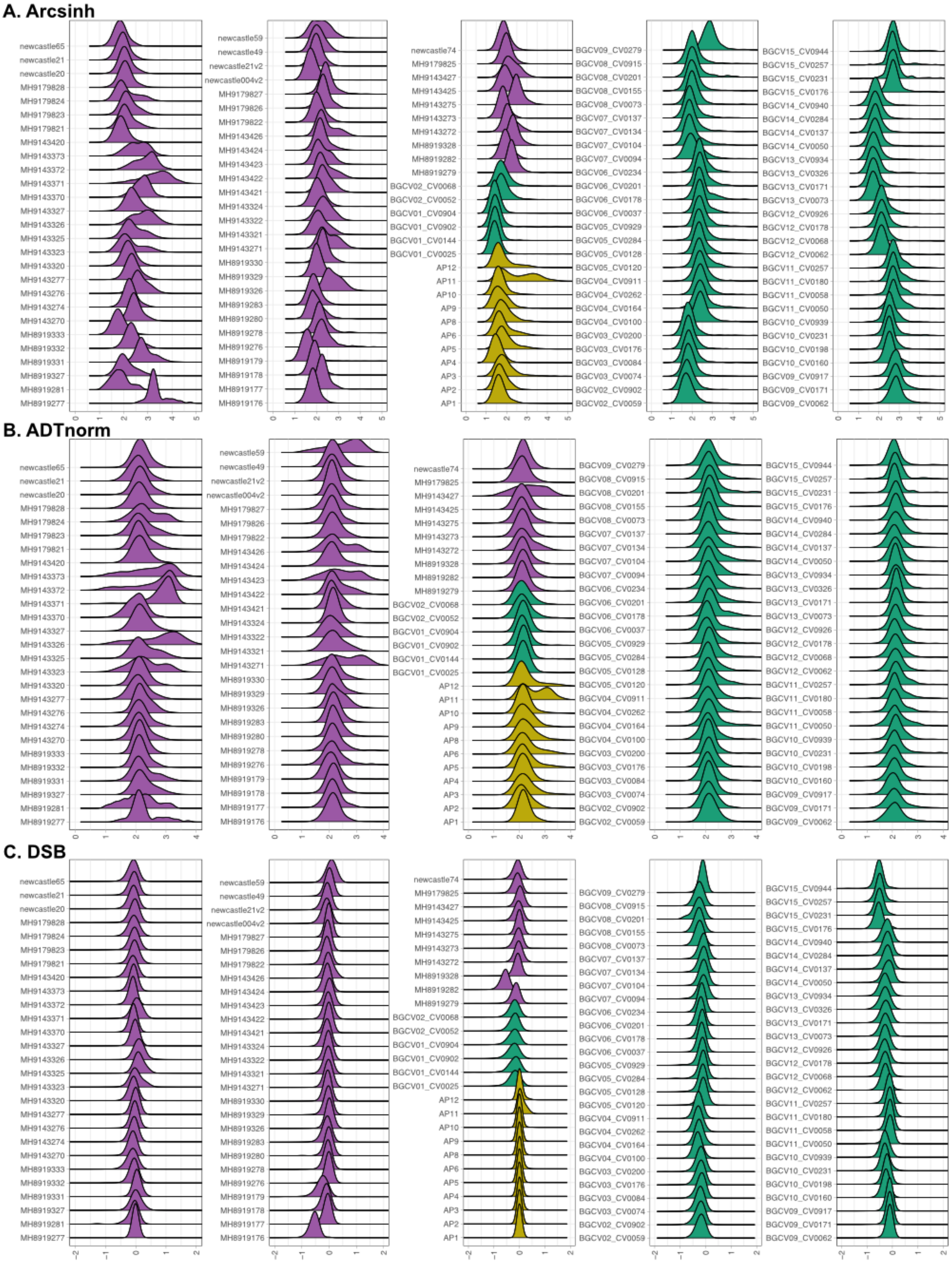


**Supplementary Figure 20 CD169 cell surface protein density distributions.** The density distribution of CD169 after Arcsinh transformation (used for better visualization of the raw protein count data, **A**), ADTnorm normalization (**B**), or DSB normalization used in the original publication (**C**). The positive population, observed in the Arcsinh and ADTnorm settings, is lost after DSB normalization.
