## Supplementary Notes for "ADTnorm: Robust Integration of Single-cell Protein Measurement across CITE-seq Datasets"

### Supplementary Note: Part 1 Protein Density Distribution

#### 1. 13 Public Datasets

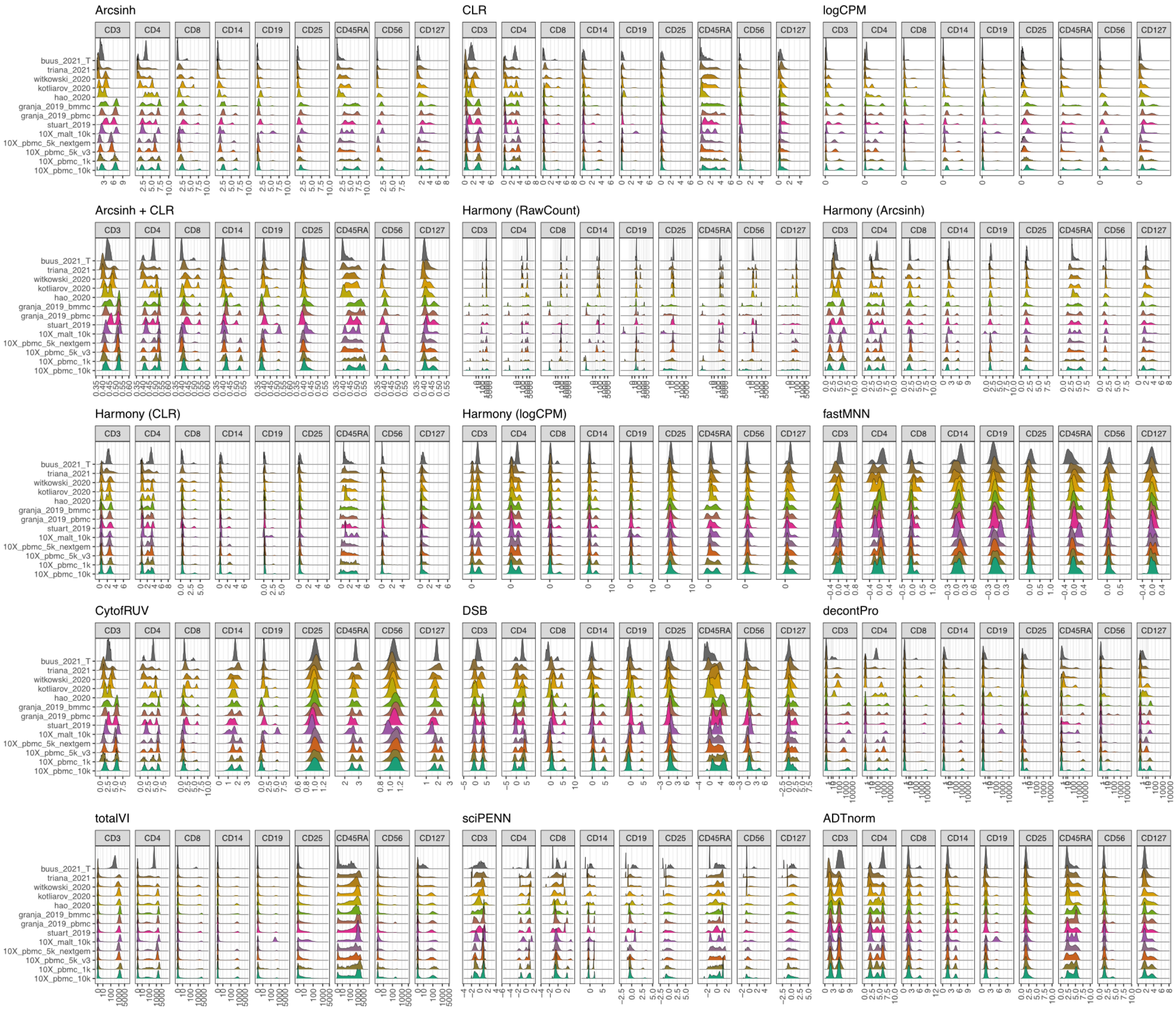


**Supplementary Note Figure 1. Normalization density distribution comparison of the nine shared protein markers across 13 public datasets and 15 normalization methods.** Among the 13 public datasets, buus_2021_T only contains T cells, and the CD3 marker is expected to have a uni-peak, which should be aligned to positive peaks.

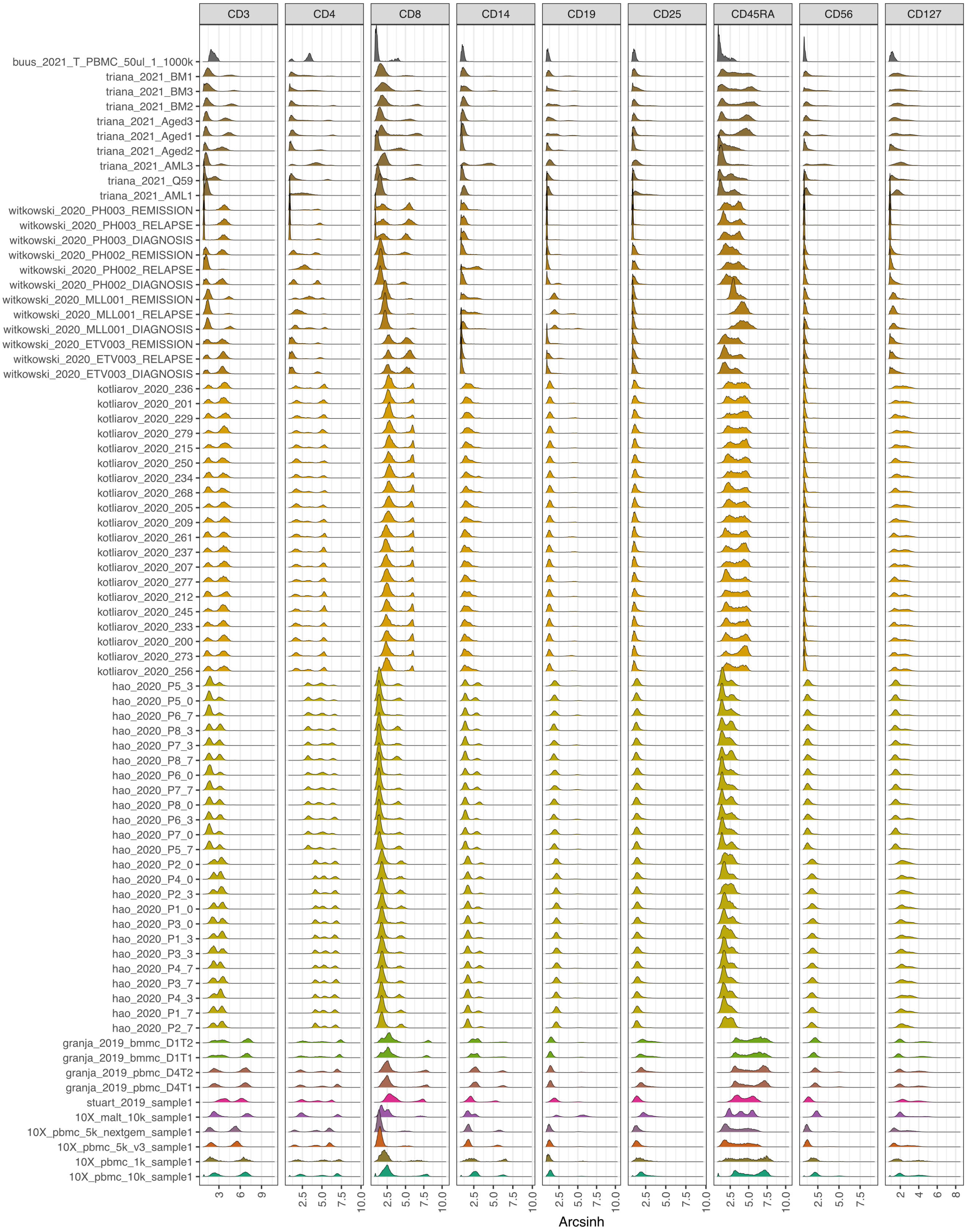


**Supplementary Note Figure 2. Arcsinh transformation of the 13 public datasets at the sample level.** Each row corresponds to a sample per study. Color corresponds to the study where the data were collected.

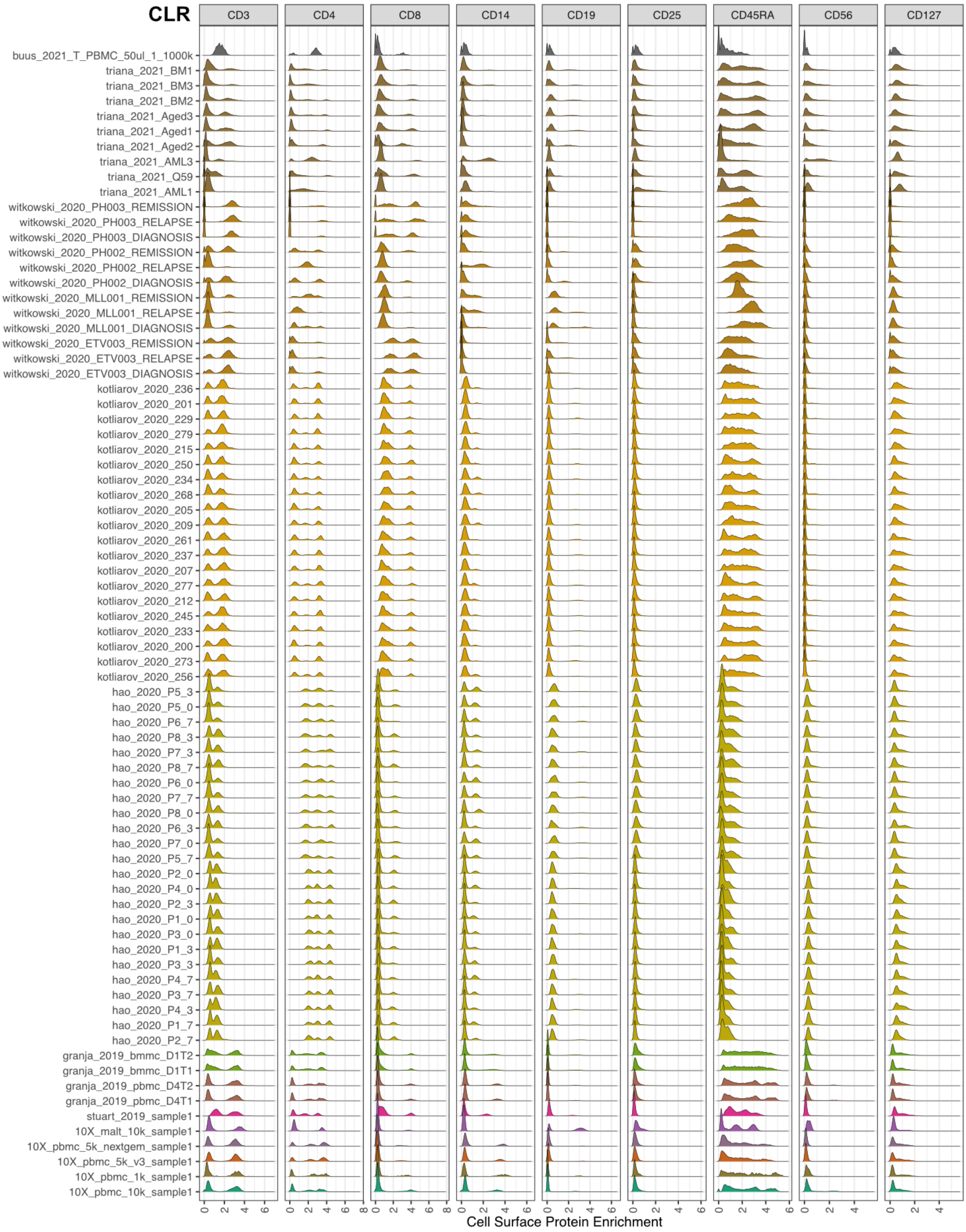


**Supplementary Note Figure 3. CLR transformation of the 13 public datasets at the sample level.** Each row corresponds to a sample per study. Color corresponds to the study where the data were collected.

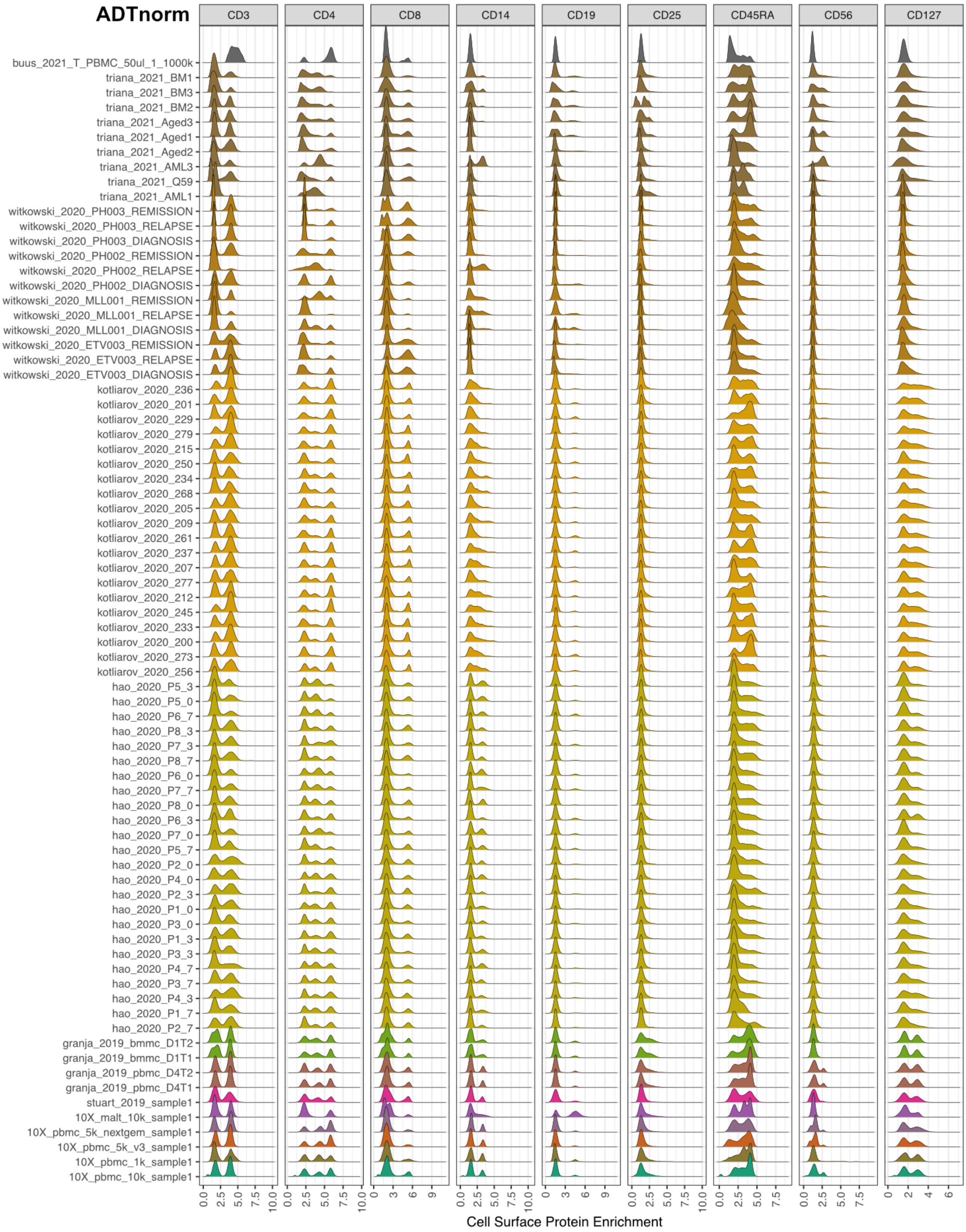


**Supplementary Note Figure 4. ADTnorm normalization on the sample unit from each study.** Each row corresponds to a sample per study. Color corresponds to the study where the data were collected. Sample from the buus_2021_T only contains T cells and CD3 marker expects to have uni-peak which should be aligned to positive peaks.

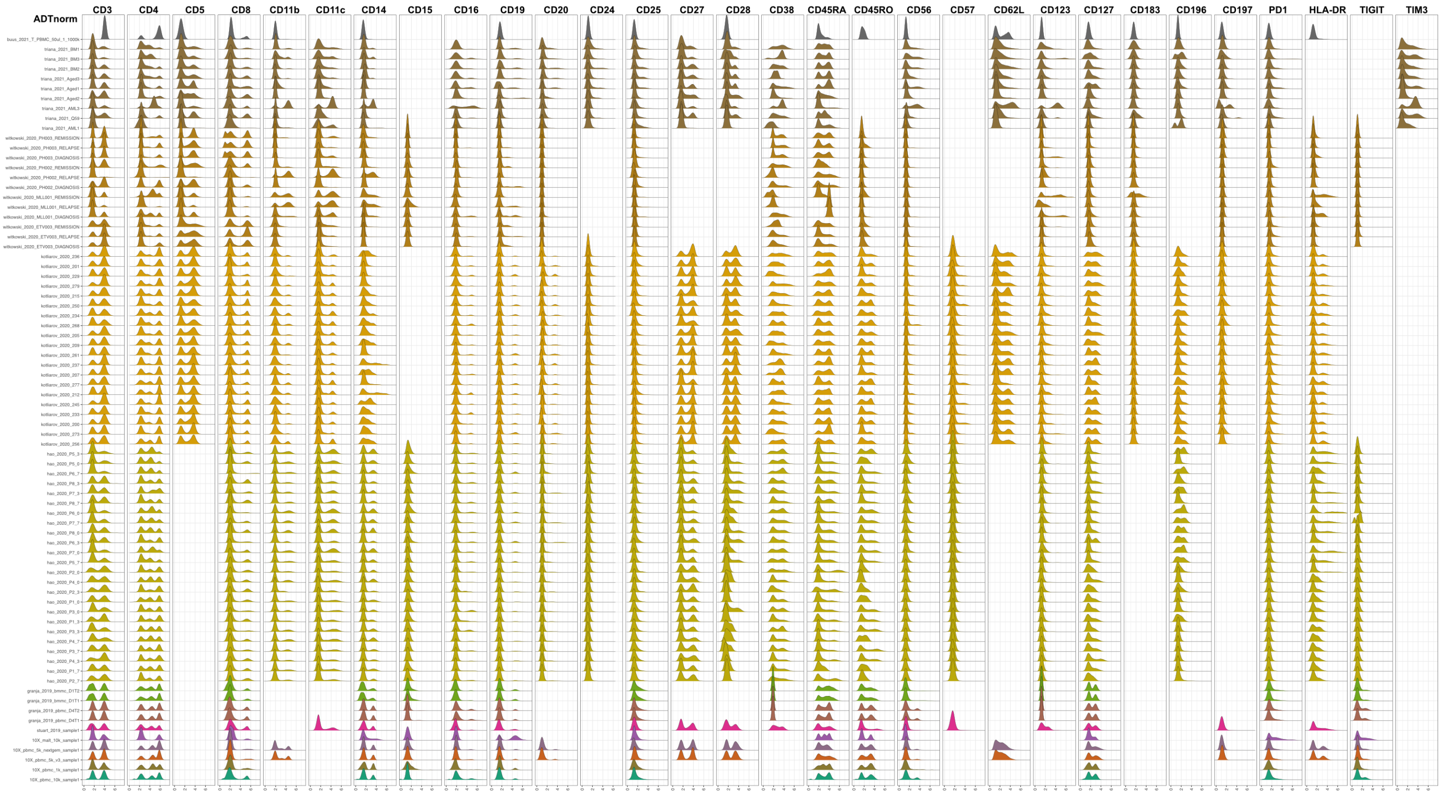


**Supplementary Note Figure 5. Density distribution of the ADTnorm normalized results across 30 protein markers with missing values.** Not all the protein markers are available across the 13 public datasets. ADTnorm can handle the missing values by aligning the peak and valley landmarks that are available.

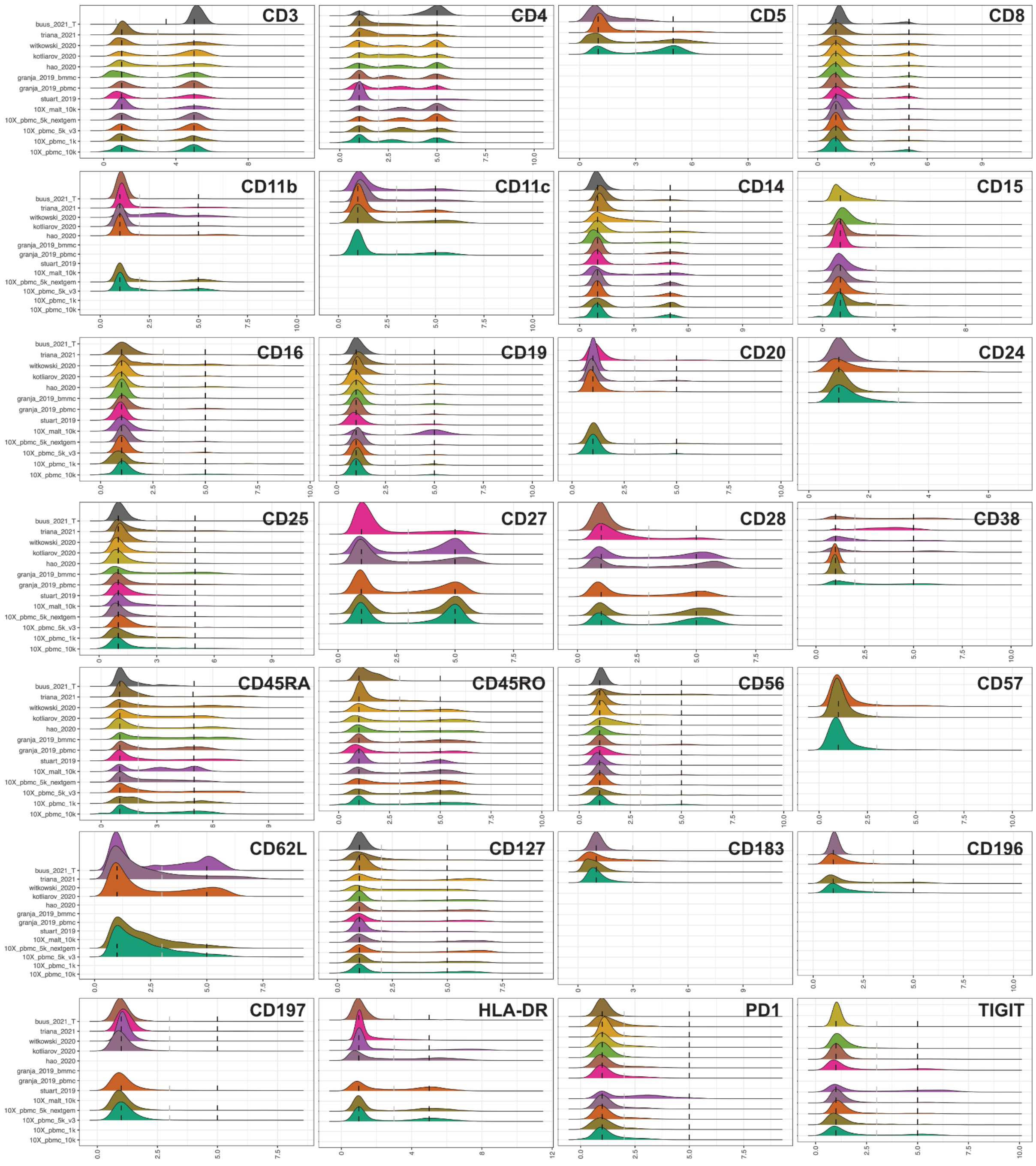


**Supplementary Note Figure 6. ADTnorm aligns the peak and valley landmarks to the fixed location.** The peak and valley landmarks were aligned to the same position across all the protein markers, with the negative peak aligned to 1, the first valley aligned to 3 and the right-most positive peak aligned to 5. Only the CD4's valley is aligned to 2 for better visualization of all three peaks. One study is considered as a batch.

### 2. Titration Study

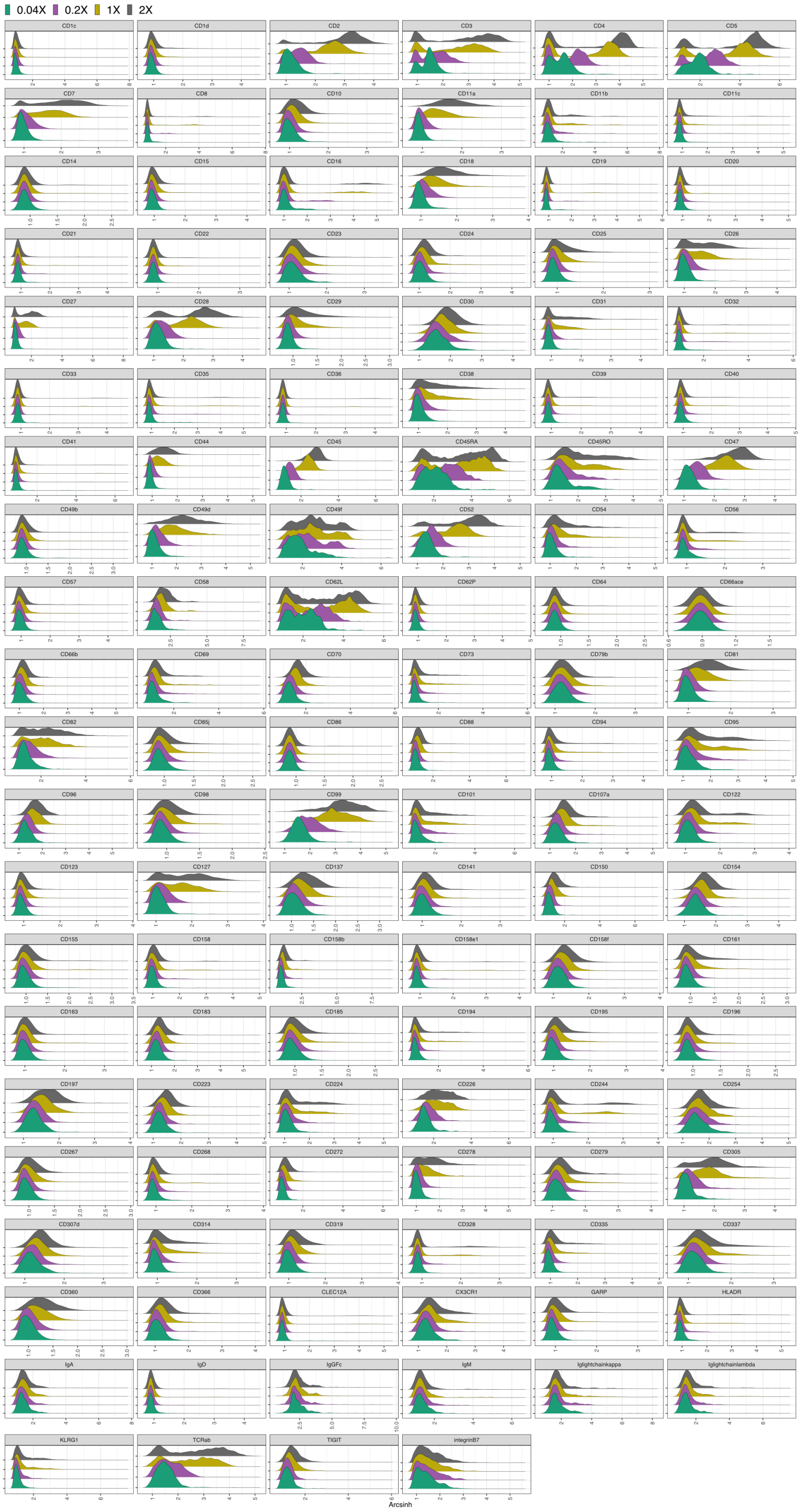


**Supplementary Note Figure 7. Arcsinh transformation of the protein expression for the titration study.** Color corresponds to the four antibody concentrations used in the titration experiment.

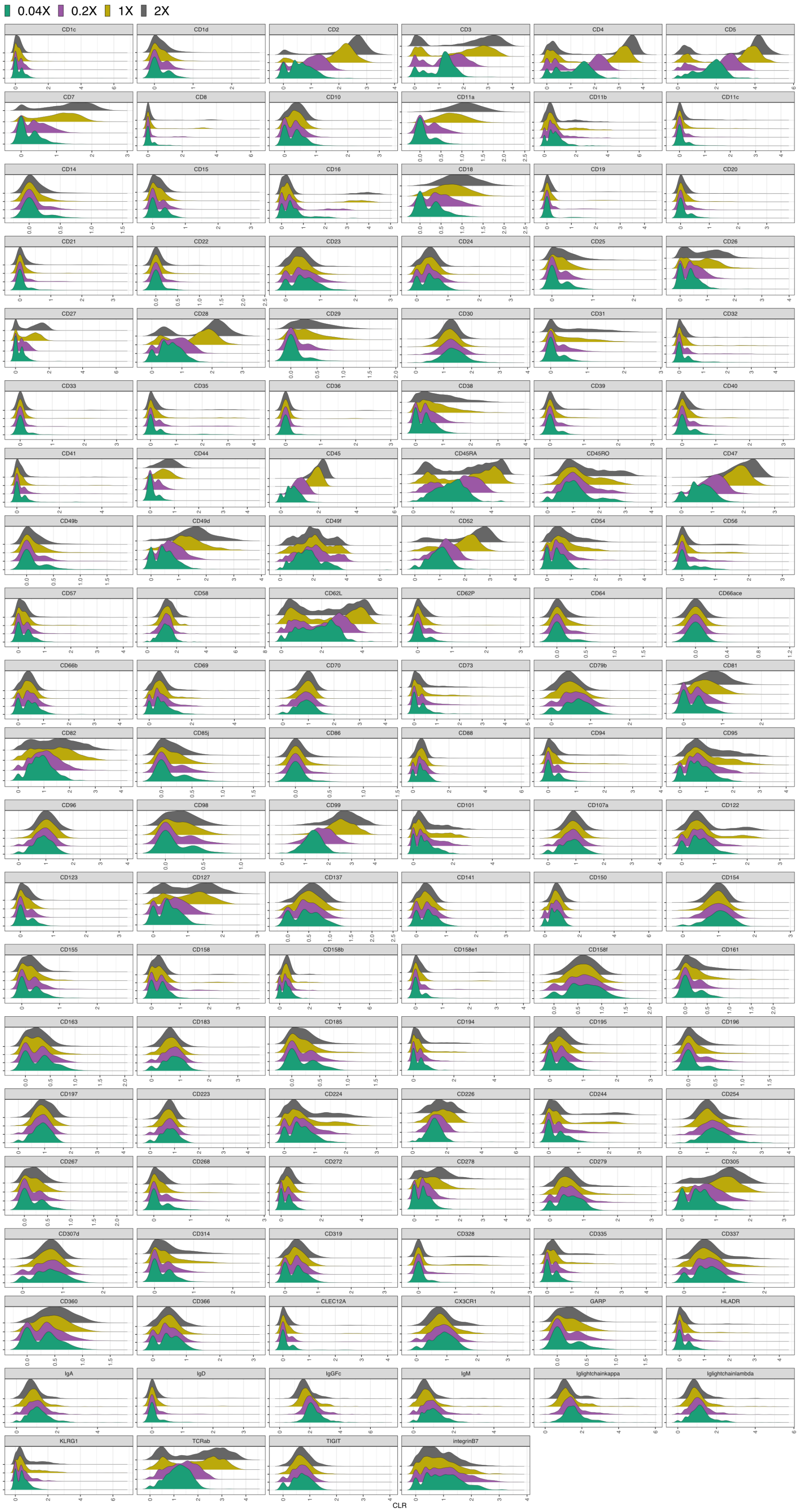


**Supplementary Note Figure 8. CLR transformation of the protein expression for the titration study.** Color corresponds to the four antibody concentrations used in each titration experiment.

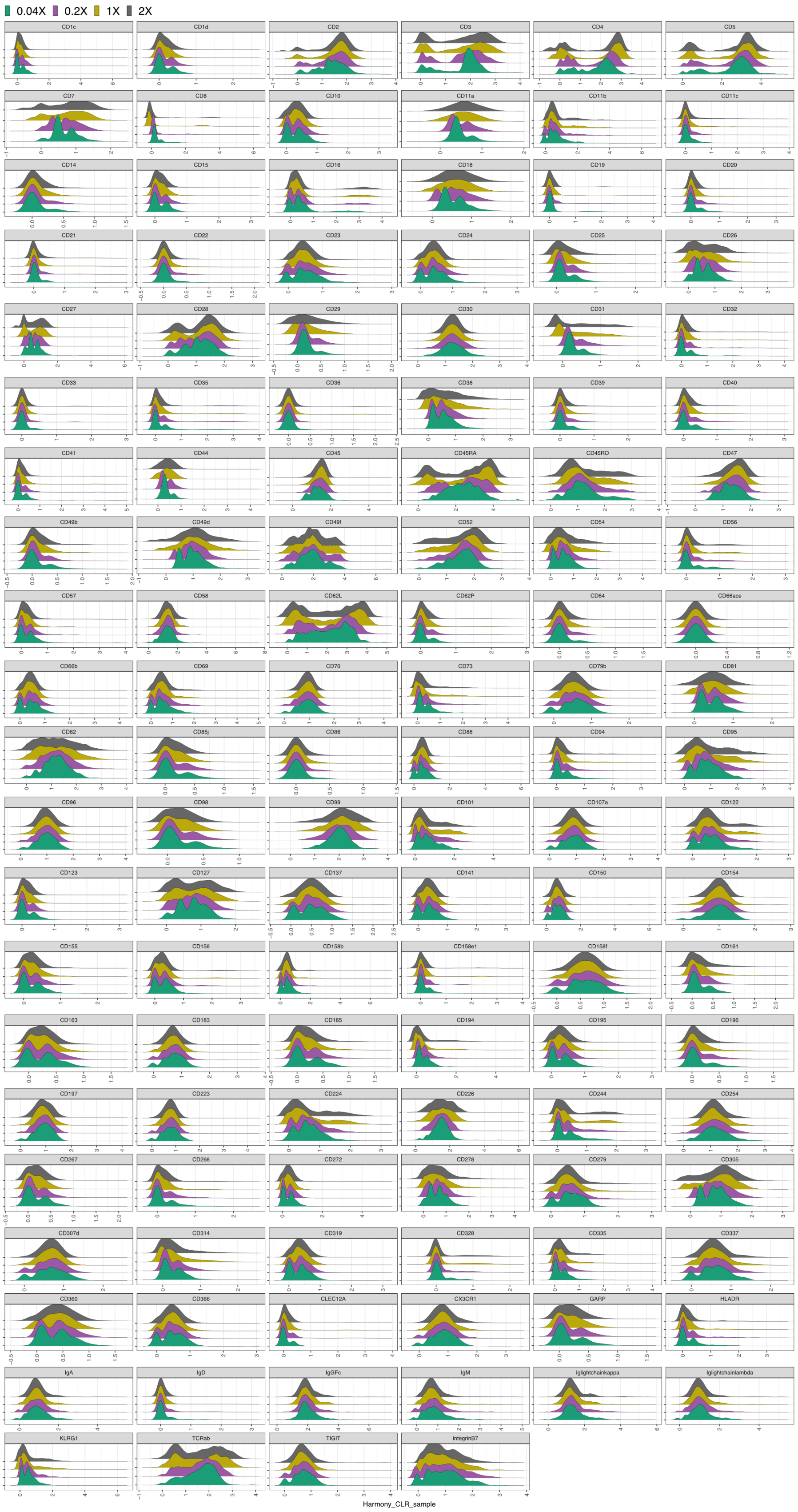


**Supplementary Note Figure 9. Density distribution of protein markers normalized by the Harmony on the CLR transformed counts for titration study.** Color corresponds to the four antibody concentrations used in each titration experiment.

### 3. COVID-19 Study

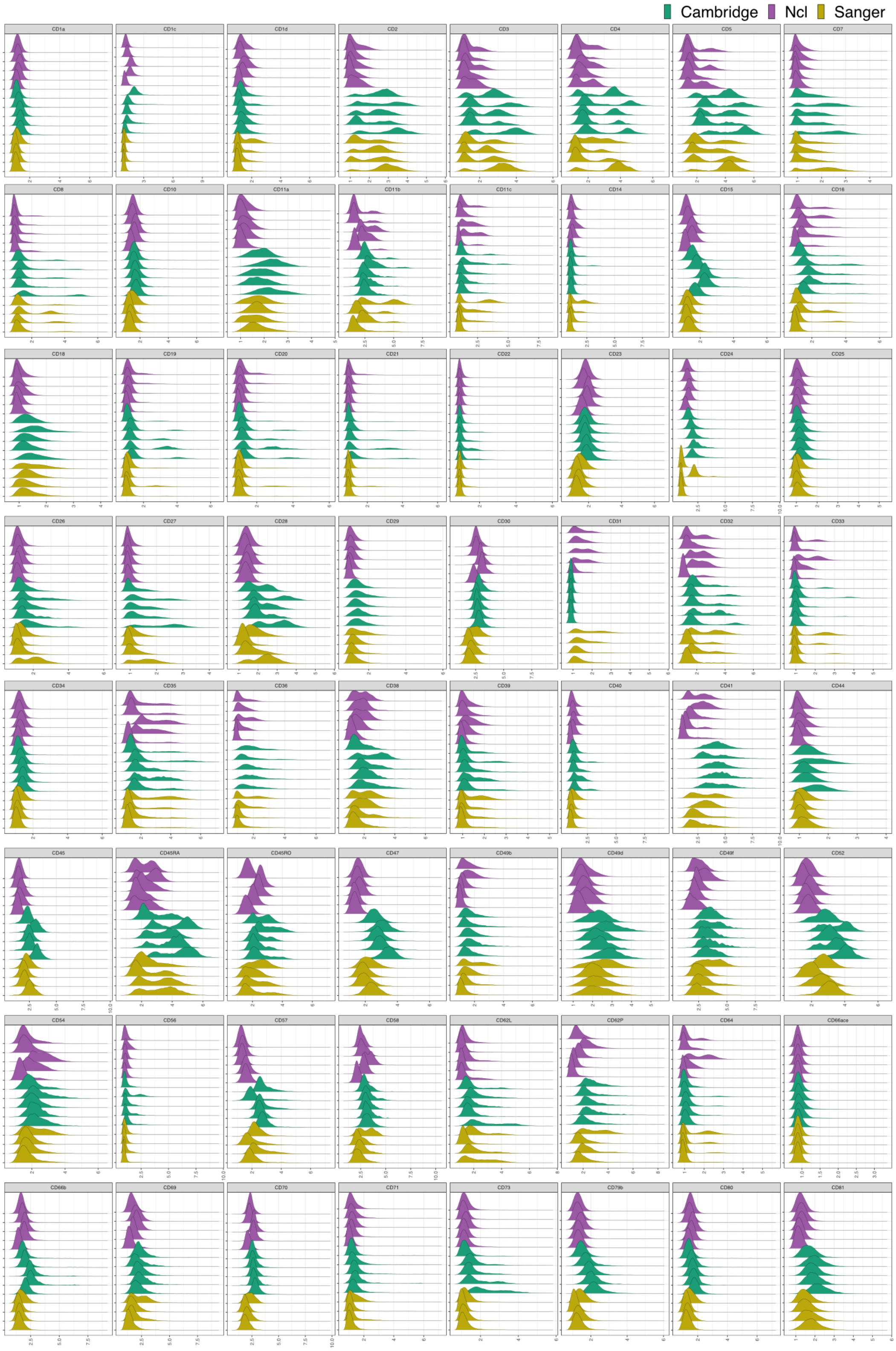


**Supplementary Note Figure 10. Arcsinh transformation of the protein data for the COVID-19 study.** Each sample is considered one batch. Color corresponds to the three research institutes where the experiments were conducted.

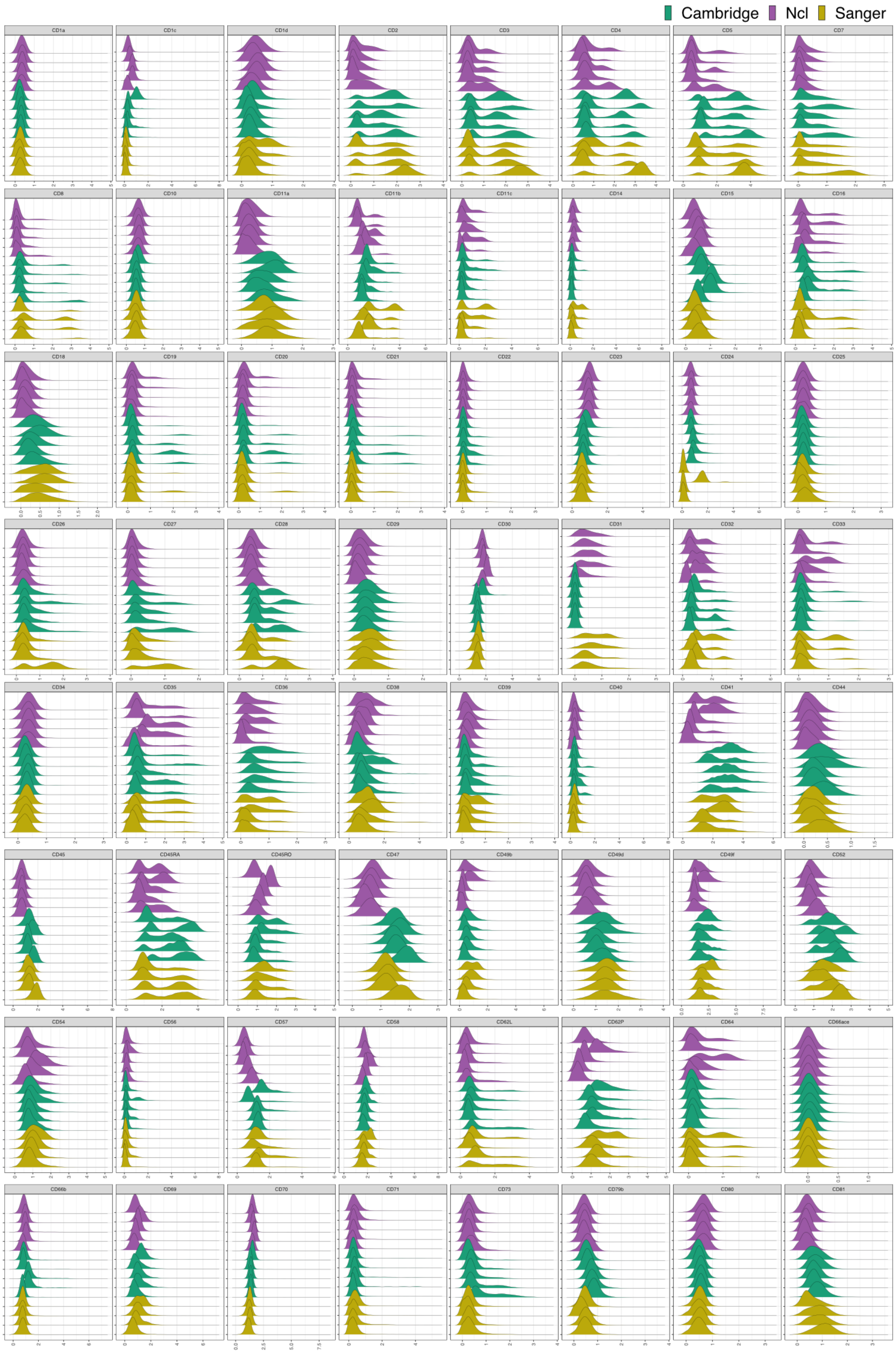


**Supplementary Note Figure 11. CLR transformation of the protein data for the COVID-19 study.** Each sample is considered one batch. Color corresponds to the three research institutes where the experiments were conducted.

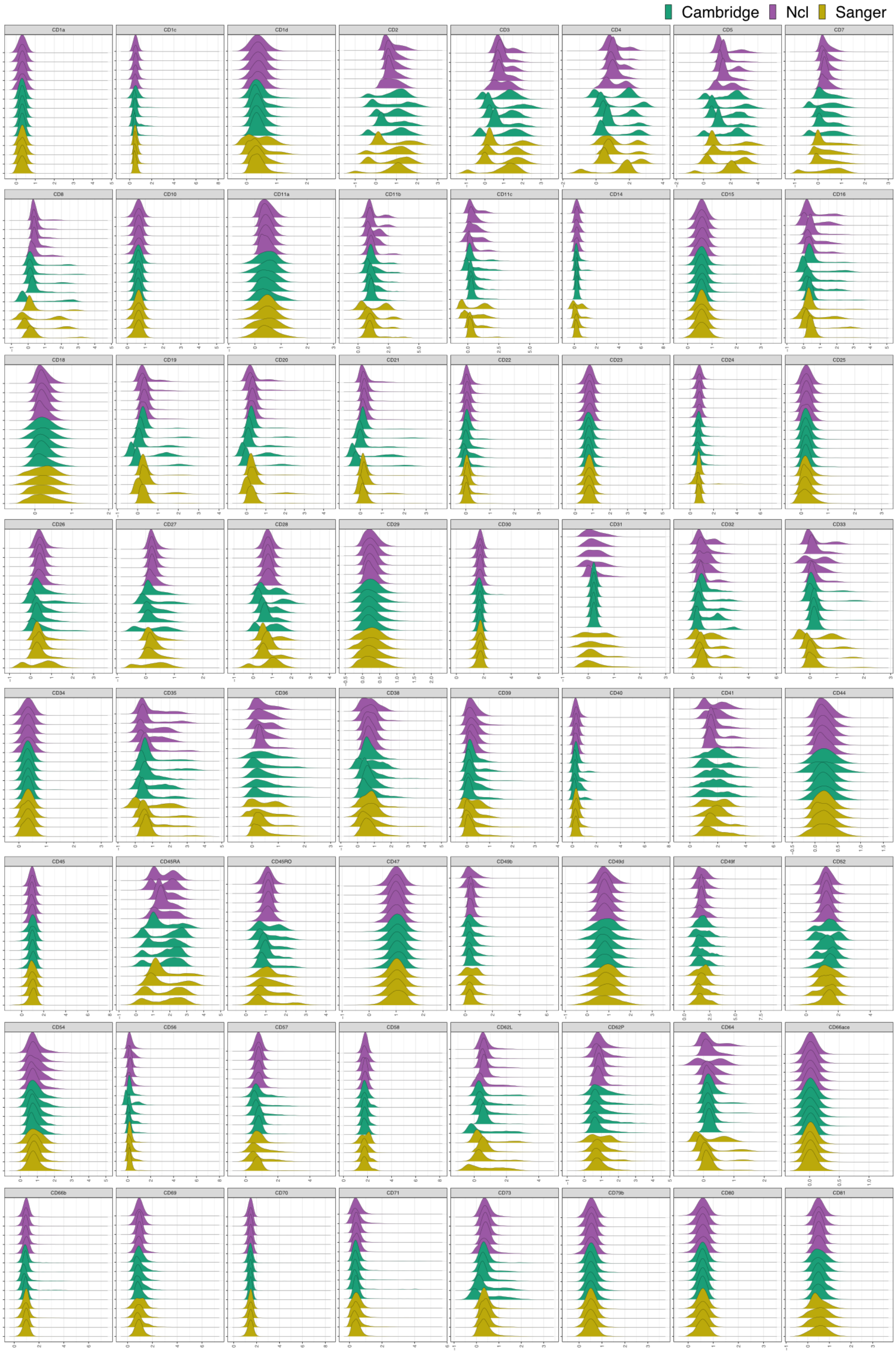


**Supplementary Note Figure 12. Harmony on the CLR transformed protein data for the COVID-19 study.** Each sample is considered one batch. Color corresponds to the three research institutes where the experiments were conducted.

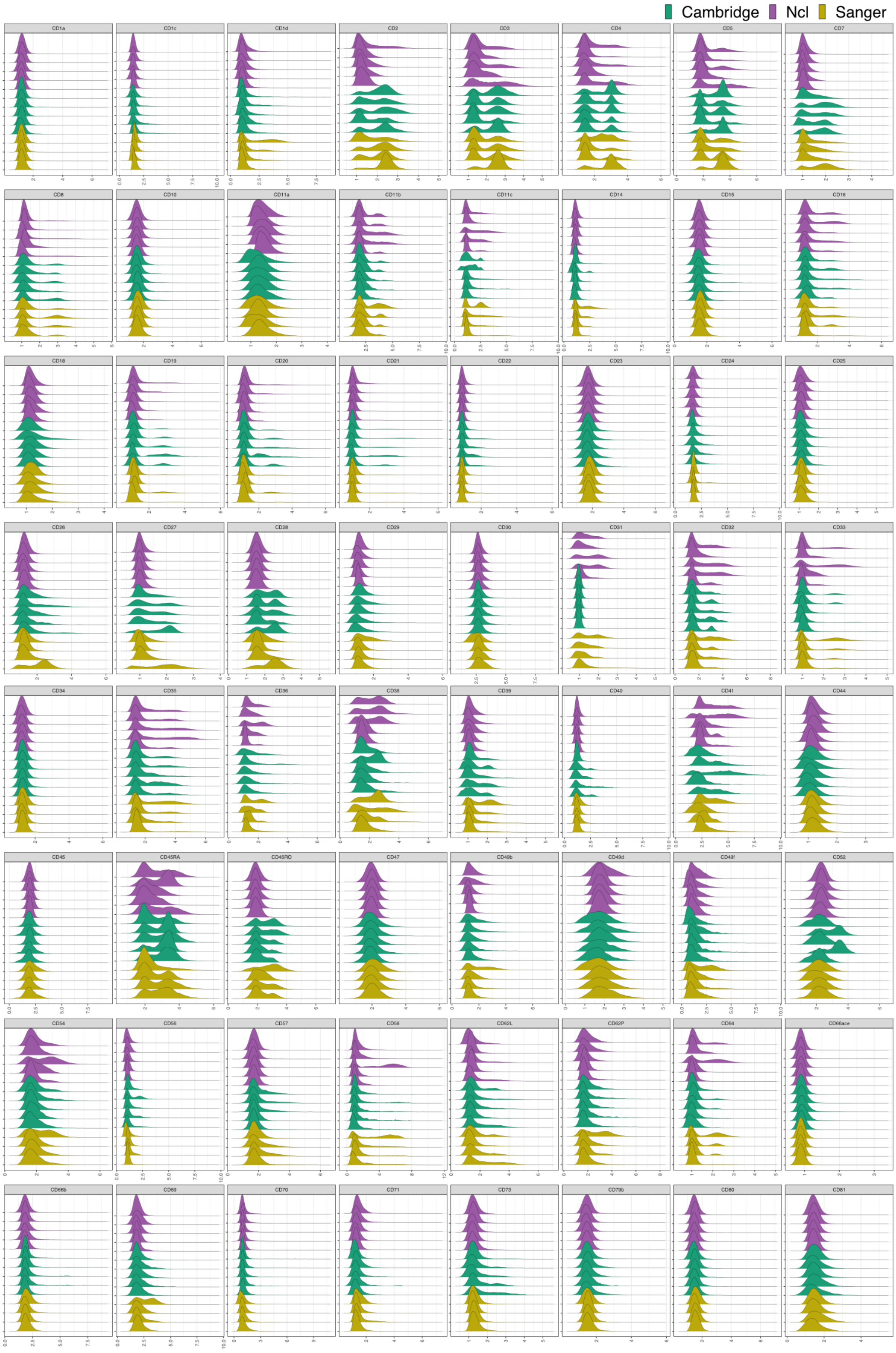


**Supplementary Note Figure 13. ADTnorm normalized the protein data for the COVID-19 study (part 1).** Each sample is considered one batch. Color corresponds to the three research institutes where the experiments were conducted.

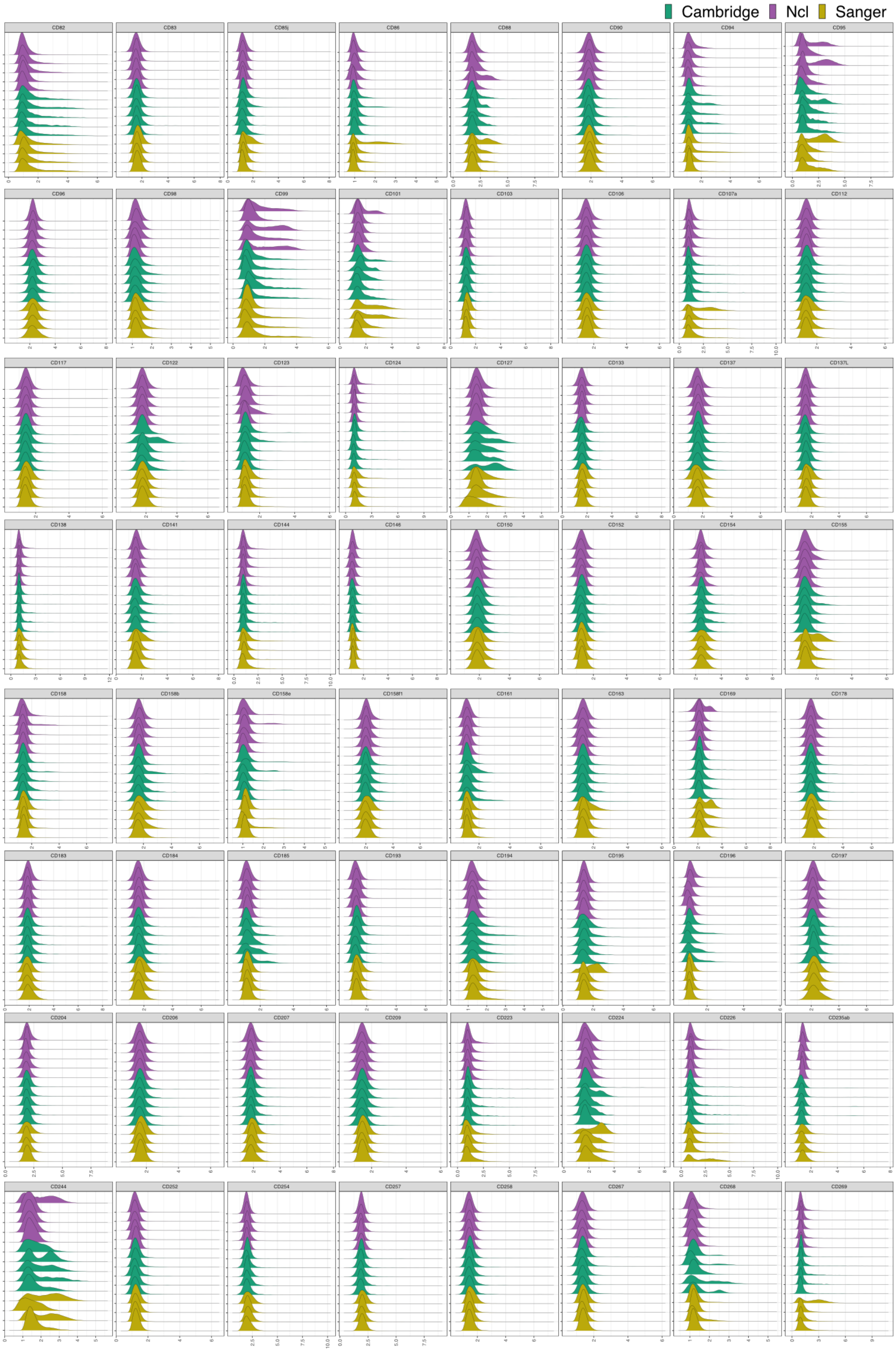


**Supplementary Note Figure 14. ADTnorm normalized the protein data for the COVID-19 study (part 2).** Each sample is considered one batch. Color corresponds to the three research institutes where the experiments were conducted.

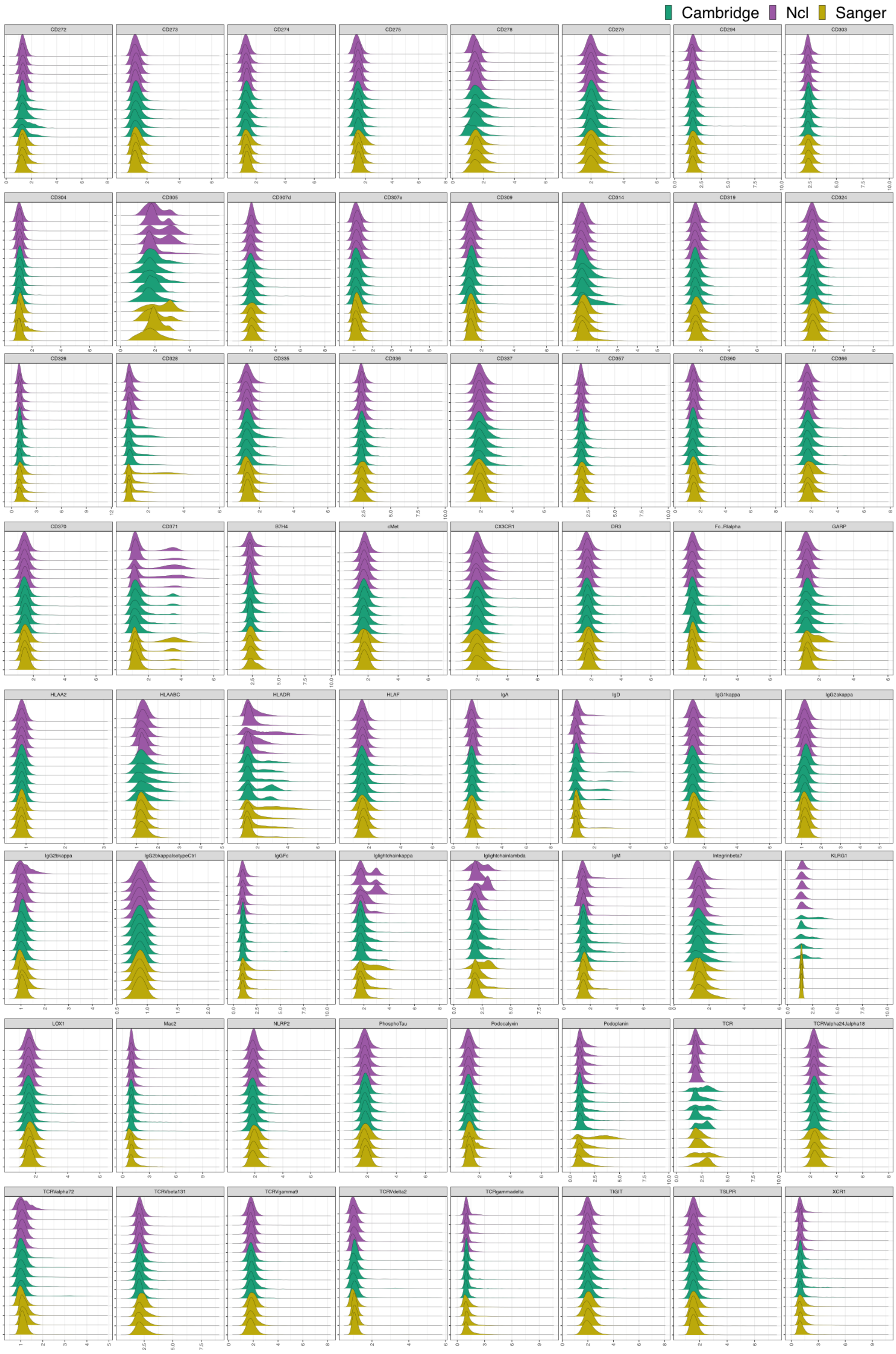


**Supplementary Note Figure 15. ADTnorm normalized the protein data for the COVID-19 study (part 3).** Each sample is considered one batch. Color corresponds to the three research institutes where the experiments were conducted.

### Supplementary Note: Part 2 ADTnorm Software Vignettes

#### 0. Introduction

CITE-seq technology enables the direct measurement of protein expression, known as antibody-derived tags (ADT), in addition to RNA expression. The increase in the copy number of protein molecules leads to a more robust detection of protein features compared to RNA, providing a deep definition of cell types. However, due to variation in antibody staining conditions, such as timing, staining concentrations and volumes, or antibody panel compositions, the batch effects of the ADT component of CITE-seq can dominate over biological variations, especially in across-study analyses. We present ADTnorm as a normalization and integration method designed explicitly for the ADT counts of CITE-seq data. Benchmarking with existing scaling and normalization methods, ADTnorm achieves a fast and accurate matching of the negative and positive peaks of the ADT counts across samples, efficiently removing technical variations across batches. Further quantitative evaluations confirm that ADTnorm achieves the best cell-type separation while maintaining the minimal batch effect. Therefore, ADTnorm facilitates the scalable ADT count integration of massive public CITE-seq datasets with distinguished experimental designs, which are essential for creating a corpus of well-annotated single-cell data with deep and standardized annotations.

#### 1. ADT Normalization Pipeline

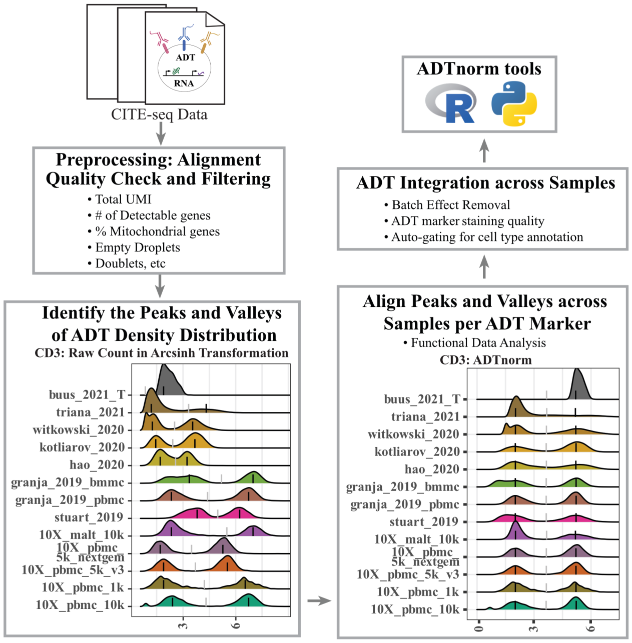


#### 2 Installation

##### 2.1 Install through GitHub

```r
# install.packages('remotes')
remotes::install_github("yezhengSTAT/ADTnorm", build_vignettes = FALSE)
```

##### 2.2 Using Docker

There are many dependencies in `ADTnorm`, so it takes a long time to install them all. Instead, you can use the Docker image of `ADTnorm`.

```
docker pull ghcr.io/yezhengstat/adtnorm:latest
docker run \
  -it \
  --user rstudio \
  --volume <yourDataDirectory>:/home/rstudio/data \
  yezhengstat/adtnorm:latest \
  R
```

Replace `<yourDataDirectory>` with the local directory path (absolute path) where you have the input data and would like to store the output files. For more information on using docker containers, please read this documentation by Bioconductor.

#### 3. Input Data

The 13 public datasets used in the manuscript are also included in the R package as a demo data set. They can be loaded by

```
data(cell_x_adt)
data(cell_x_feature)
```

- `cell_x_adt` contains raw counts for the ADT markers in each cell, as a data frame with 422682 cells (rows) and 9 ADT markers (columns): CD3, CD4, CD8, CD14, CD19, CD25, CD45RA, CD56, CD127.

```
  CD3  CD4 CD8 CD14 CD19 CD25 CD45RA CD56 CD127
1  18  138  13  491    3    9    110   17     7
2  30  119  19  472    3    5    125  248     8
3  18  207  10 1289    8   15   5268   26    12
4  18   11  17   20    5   15   4743  491    16
5   5   14  14   19    4   16   4108  458    17
6  21 1014  29 2428    7   52    227   29    15
```

- `cell_x_feature` is a data frame with 422682 cells (rows) and 7 feature variables (columns):
  - sample: Sample name used in original data of each study.
  - batch: Batch information provided from each study.
  - sample_status: Sample status, i.e., Healthy, MALTtumor, HIV Vaccine, Lupus, B-ALL, AML.
  - study_name: Name of the data set/study.
  - ADTseqDepth: Total UMI per cell.
  - cell_type_l1: Broad level of cell type annotation using manual gating.
  - cell_type_l2: Fine level of cell type annotation using manual gating.

```
              sample               batch sample_status   study_name
1 10X_pbmc_10k_sample1 10X_pbmc_10k_batch1       healthy 10X_pbmc_10k
2 10X_pbmc_10k_sample1 10X_pbmc_10k_batch1       healthy 10X_pbmc_10k
3 10X_pbmc_10k_sample1 10X_pbmc_10k_batch1       healthy 10X_pbmc_10k
4 10X_pbmc_10k_sample1 10X_pbmc_10k_batch1       healthy 10X_pbmc_10k
5 10X_pbmc_10k_sample1 10X_pbmc_10k_batch1       healthy 10X_pbmc_10k
6 10X_pbmc_10k_sample1 10X_pbmc_10k_batch1       healthy 10X_pbmc_10k
  ADTseqDepth cell_type_l1       cell_type_l2
1         981    monocytes classical monocyte
```

```
2        1475    monocytes classical monocyte
3        7149    monocytes classical monocyte
4        6831           NK           CD16+ NK
5        6839           NK           CD16+ NK
6        4720    monocytes classical monocyte
```

#### 4. Quick start

For a quick introduction to using ADTnorm, we will treat all the cells from one study as one batch and normalize across studies.

##### 4.1 ADTnorm general usage

```r
library(ADTnorm)
save_outpath <- "/path/to/output/location"
run_name <- "ADTnorm_demoRun"
data(cell_x_adt)
data(cell_x_feature)

cell_x_feature$sample = factor(cell_x_feature$study_name)
cell_x_feature$batch = factor(cell_x_feature$study_name)

cell_x_adt_norm = ADTnorm(
    cell_x_adt = cell_x_adt,
    cell_x_feature = cell_x_feature,
    save_outpath = save_outpath,
    study_name = run_name,
    marker_to_process = NULL, # setting it to NULL by default will process all available markers in cell_x_adt.
    bimodal_marker = NULL, # setting it to NULL will trigger ADTnorm to try different settings to find bimodal peaks for all the markers.
    trimodal_marker = c("CD4", "CD45RA"), # CD4 and CD45RA tend to have three peaks.
    positive_peak = list(ADT = "CD3", sample = "buus_2021_T"), # setting the CD3 uni-peak of buus_2021_T study to positive peak if only one peak is detected for CD3 marker.
    brewer_palettes = "Dark2", # color brewer palettes setting for the density plot
    save_fig = TRUE
)
```

##### 4.2 ADTnorm basic parameter explanation

```
cell_x_adt:         Matrix of ADT raw counts in cells (rows) by ADT markers (columns)
↪ format.

cell_x_feature:     Matrix of cells (rows) by cell features (columns) such as sample,
↪ batch, and cell type-related information. Please note "sample" column is mandatory
↪ and should be the smallest unit to group the cells. At this resolution, ADTnorm will
↪ identify peaks and valleys to implement normalization. Please make sure the samples
↪ will have different names across batches/conditions/studies. "batch" column is
↪ optional. It can be batches/conditions/studies/etc that group the samples based on
↪ whether the samples are collected from the same batch run or experiment. This column
↪ is needed if the ```multi_sample_per_batch``` parameter is turned on to remove
↪ outlier positive peaks per batch. If the "batch" column is not provided, it will be
↪ set as the same as the "sample" column. In the intermediate density plots that
↪ ADTnorm provides, density plots will be colored by the "batch" column.
```

```
save_outpath:       The path to save the results.

study_name:         Name of this run.

marker_to_process:  Markers to normalize. Leave empty to process all the ADT markers in
↪ the cell_x_adt matrix.

bimodal_marker:     Specify ADT markers that are likely to have two peaks based on
↪ researchers' prior knowledge or preliminary observation of the particular data to be
↪ processed. Leaving it as default, ADTnorm will try to find the bimodal peak in all
↪ markers that are not listed in `trimodal_marker.`

trimodal_marker:    Index of the ADT markers that tend to have three peaks based on
↪ researchers' prior knowledge (e.g., CD4) or preliminary observation on particular
↪ data to be processed.

positive_peak:      A list variable containing a vector of ADT marker(s) and a
↪ corresponding vector of sample name(s) in matching order to specify that the uni-peak
↪ detected should be aligned to positive peaks. For example, for samples that only
↪ contain T cells, the only CD3 peak should be aligned to the positive peaks of other
↪ samples.

save_fig:  Save the density plot figure for checking the peak and valley location
↪ detection.
```

The full parameter explanation for the `ADTnorm` function can be found at Reference - ADTnorm. In the next section, we will show the usage of the rest of the parameters and go through some typical parameter tuning examples.

##### 4.3 Results

`cell_x_adt_norm` is the normalized ADT counts of the same dimension of `cell_x_adt`.

```
       CD3      CD4      CD8     CD14     CD19     CD25   CD45RA     CD56
1 1.969350 3.292308 1.637615 3.885946 1.296276 1.328292 2.728777 1.510205
2 2.285588 3.179321 1.890224 3.854359 1.296276 1.060717 2.825304 3.285286
3 1.969350 3.612930 1.482134 4.704417 1.713140 1.634128 5.858261 1.735593
4 1.969350 1.620001 1.812941 1.603789 1.484578 1.634128 5.753854 3.926593
5 1.321316 1.753275 1.684441 1.573910 1.394763 1.677503 5.615075 3.857970
6 2.062768 5.069278 2.206177 5.306176 1.642843 2.628830 3.278655 1.796711
     CD127
1 1.335782
2 1.394575
3 1.594514
4 1.754638
5 1.790186
6 1.717475
```

Additionally, in the `save_outpath` specified by the users, there will be subfolders: `figures` (if setting `save_fig to TRUE``) and `RDS` (if setting`save_landmarktoTRUE‘`). Thefigures`subfolder contains intermediate figures to check if peaks and valleys are accurately identified. Therefore, users can decide if further parameter tuning or manual adjustment on the landmark locations is needed for certain ADT markers. The`RDS‘ subfolder contains the peaks and valleys location objects before and after ADTnorm.

#### 5. ADTnorm parameter tuning examples

##### 5.1 Clean ADT marker name

ADTnorm provides a quick and naive ADT marker cleaning and unifying function, `clean_adt_name`, which will be able to remove unwanted suffixes such as “_TotalSeqB”, “*PROT” or prefixes such as ”ADT*”. Or, if there is no “CD8” marker detected, look for “CD8A” or “CD8a” to work as “CD8”. Users need to clean and unify the ADT marker names first before running ADTnorm (for example, “PDL1” may also be named as “CD274”. Choose one and use it consistently) and please do not rely too much on this function to clean ADT marker names across the study. Users may also refer to AbNames package to match antibody names to gene or protein identifiers and to help standardize names across data sets.

##### 5.2 Align the negative peak and the right-most positive peak to a fixed location

`target_landmark_location`: Align the landmarks to a fixed location or, by default, align to the mean across samples for each landmark. The default value is NULL. Setting it to “fixed” will align the negative peak to 1 and the right-most positive peak to 5. Users can also assign a two-element vector indicating the location of the negative and most positive peaks to be aligned.

For example, in the “Quick start” example, setting `target_landmark_location` to be fixed will align the negative peak to 1 and the right-most positive peak to 5.

```
library(ADTnorm)
save_outpath <- "/path/to/output/location"
run_name <- "ADTnorm_demoRun_fixedAlignment"
data(cell_x_adt)
data(cell_x_feature)

cell_x_feature$sample = factor(cell_x_feature$study_name)
cell_x_feature$batch = factor(cell_x_feature$study_name)

cell_x_adt_norm = ADTnorm(
    cell_x_adt = cell_x_adt,
    cell_x_feature = cell_x_feature,
    save_outpath = save_outpath,
    study_name = run_name,
    marker_to_process = NULL, # setting it to NULL by default will process all available
    markers in cell_x_adt.
    bimodal_marker = NULL, # setting it to NULL will trigger ADTnorm to try different
    settings to find bimodal peaks for all the markers.
    trimodal_marker = c("CD4"), # CD4 and CD45RA tend to have three peaks.
    positive_peak = list(ADT = "CD3", sample = "buus_2021_T"), # setting the CD3 uni-peak
    of buus_2021_T study to positive peak if only one peak is detected for CD3 marker.
    brewer_palettes = "Dark2", # color brewer palettes setting for the density plot
    save_fig = TRUE,
    target_landmark_location = "fixed"
)
```

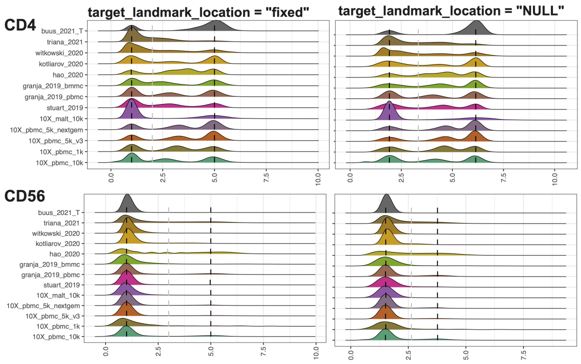


Remarkably, setting to fixed locations can eliminate the value range discrepancies across ADT markers. Also, when additional new samples are added, users can only run ADTnorm on the new samples and align to the same target locations for negative peaks and positives.

##### 5.3 Different peak alignment option

`landmark_align_type`: Algin the peak and valleys using one of the “negPeak”, “negPeak_valley”, “negPeak_valley_posPeak”, and “valley” alignment modes.

By default, ADTnorm will align the negative peaks across samples, align first valleys across samples and align the right-most positive peaks simultaneously. However, ADTnorm does have options to only align negative peaks across samples (landmark_align_type = “negpeak”), or only align negative peaks and first valleys (landmark_align_type = “negPeak_valley”), or only align the first valleys across samples (landmark_align_type = “valley”). We strongly recommend using the default option, “negPeak_valley_posPeak”, unless user have a strong reason to choose another option.

##### 5.4 Definition of the “peak”

`peak_type`: The type of peak to be detected. Select from “midpoint” for setting the peak landmark to the midpoint of the peak region being detected or “mode” for setting the peak landmark to the mode location of the peak. “midpoint” can generally be more robust across samples and less impacted by the bandwidth. “mode” can be more accurate in determining the peak location if the bandwidth is generally ideal for the target marker.

Generally, we recommend using “midpoint”. However, suppose users observe that the negative peaks modes have obvious misalignment after ADTnorm (see below). In that case, users can consider using “mode” and see if a better peak alignment result can be obtained.

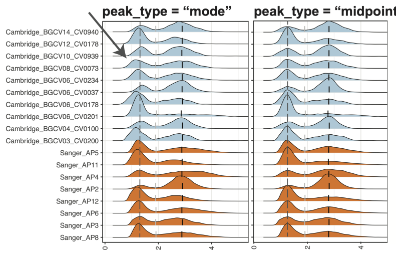


##### 5.5 Ignore the only positive peak per batch

`multi_sample_per_batch`: Set it to TRUE to discard the positive peak that only appears in one sample per batch (sample number is >=3 per batch).

There are cases where there is only one sample that has a detected positive peak. Therefore, there is no need to align the only positive peak across samples and imputing the rest of the positive peaks is not proper. Setting `multi_sample_per_batch` to TRUE will remove the only detected positive peak and only align the negative peak and valley.

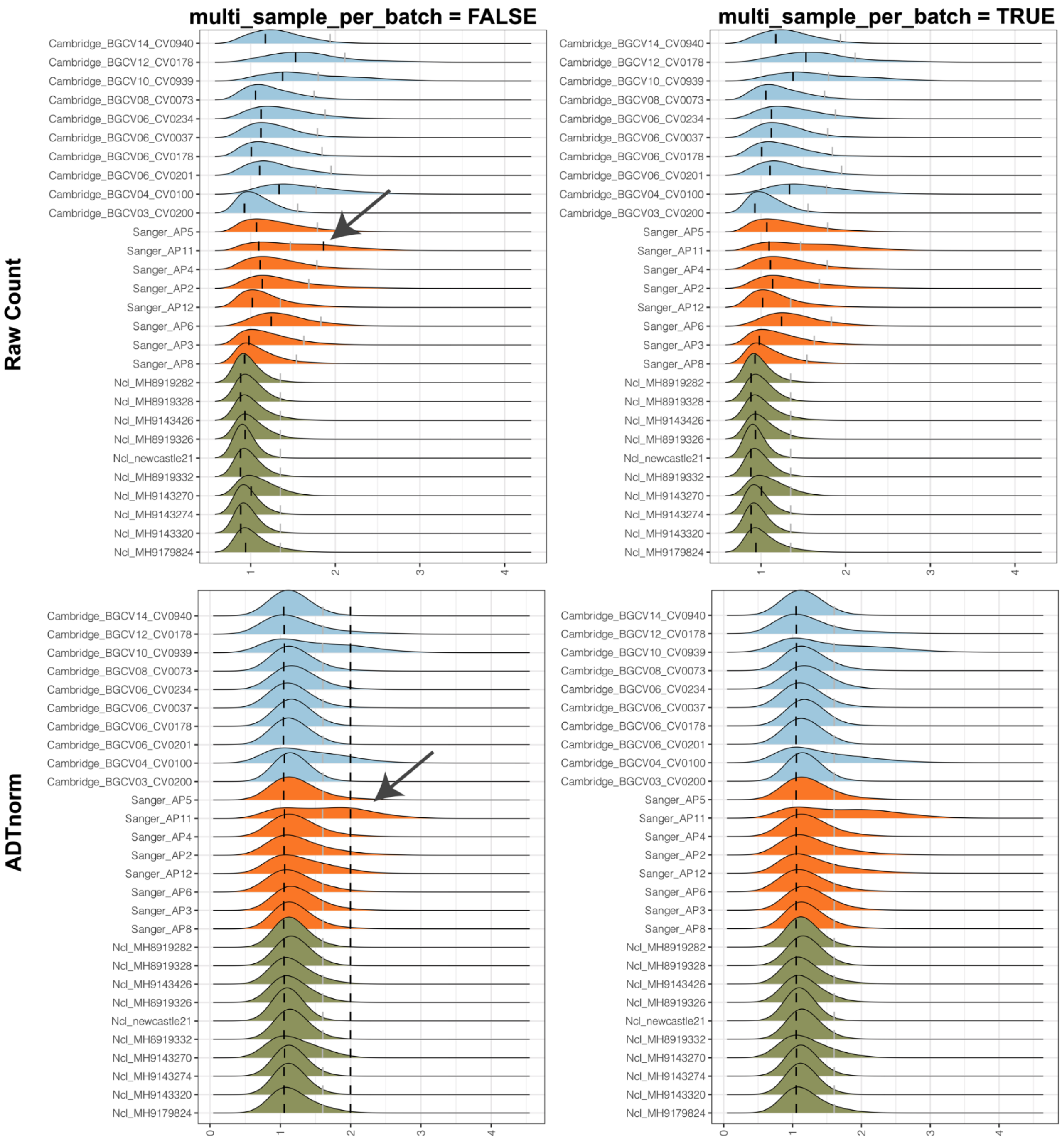


##### 5.6 Remove suspicious low signal positive peaks

`lower_peak_thres`: The minimal ADT marker density height of calling it a real peak. Set it to 0.01 to avoid a suspicious positive peak. Set it to 0.001 or smaller to include some small but usually real positive peaks, especially for markers like CD19.

Sometimes, the only positive peak can be artificial and has a very weak peak signal. Users can use this parameter to remove unwanted low signal positive peaks. The minimal default threshold to call a peak is 0.001. For example, for the COVID-19 data set used in the manuscript, to exclude the unwanted low signal peak of CD15, `lower_peak_thres` can be set to 0.01.

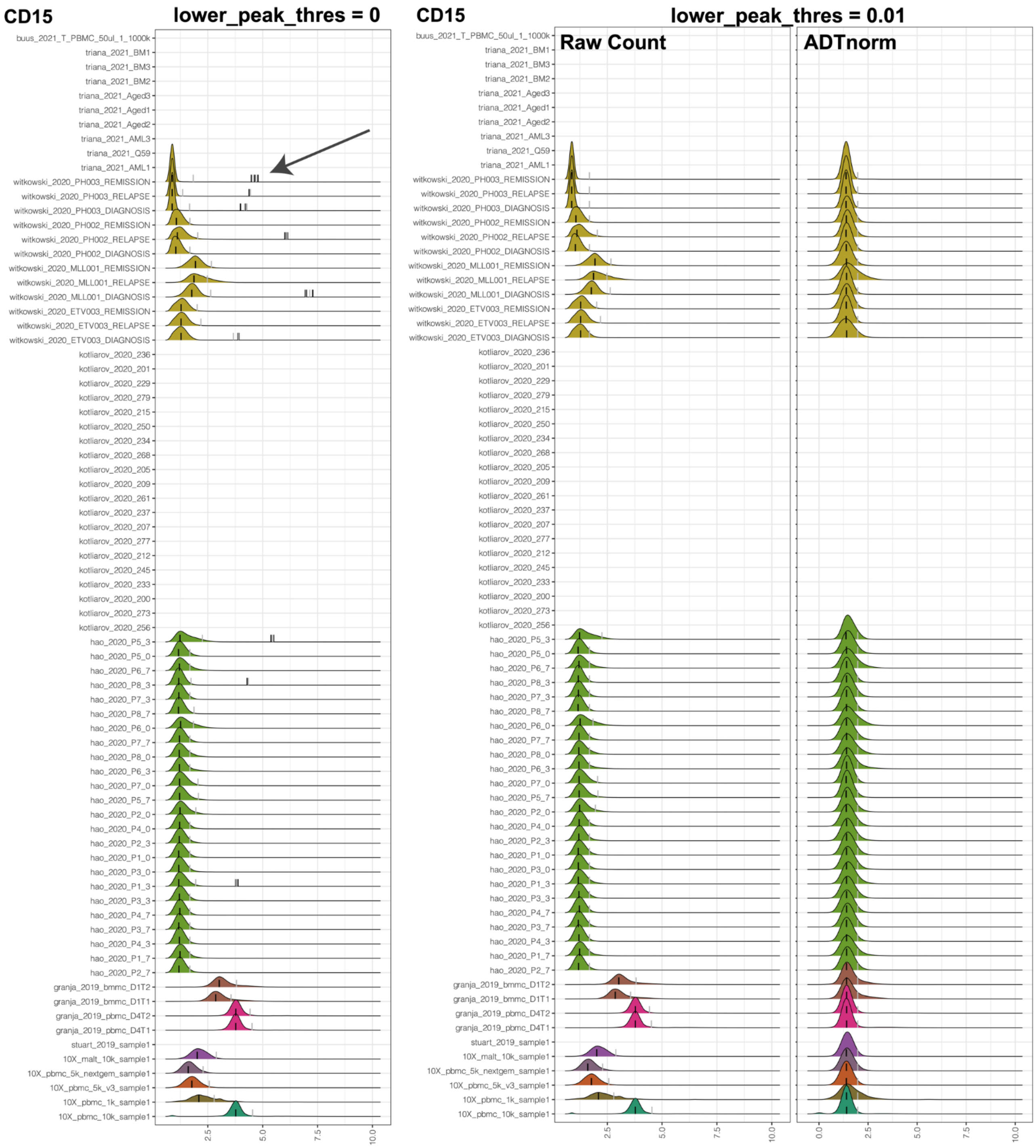

However, for markers such as CD19, we may expect to have a low signal-positive peak. `lower_peak_thres` can be decreased to 0.001 or a much smaller value to include those low signal peaks.

##### 5.7 Detect shoulder peak as the positive population.

`shoulder_valley`: Indicator to specify whether a shoulder valley is expected in case of the heavy right tail where the population of cells should be considered as a positive population.

`shoulder_valley_slope` The slope on the ADT marker density distribution to call shoulder valley.

Due to technical variations, the positive population of cells may not have a very clear separation from the negative population. Instead of having a separable positive peak, those positive populations may overlap largely with the negative ones, leading to a shoulder peak or heavy right tail pattern on the ADT count density distribution. `shoulder_valley` will allow users to turn on the detection of valley location around the "shoulder" location of the negative peak, and `shoulder_valley_slope` will allow users to obtain a relatively more stringent shoulder valley or more relaxing shoulder valley. Setting `shoulder_valley_slope` to a more negative value, such as -1, -2, ADTnorm will set the valley to a more sharply decreasing location (slope is sharper) hence closer to the negative peak mode. Setting `shoulder_valley_slope` to a less negative value, such as -0.5, -0.1, and ADTnorm will set the valley towards the tail region of the negative peak. Users can set `save_fig` to TRUE to see the ADT count density plot and the location of valleys and peaks. Those figures will help diagnose.

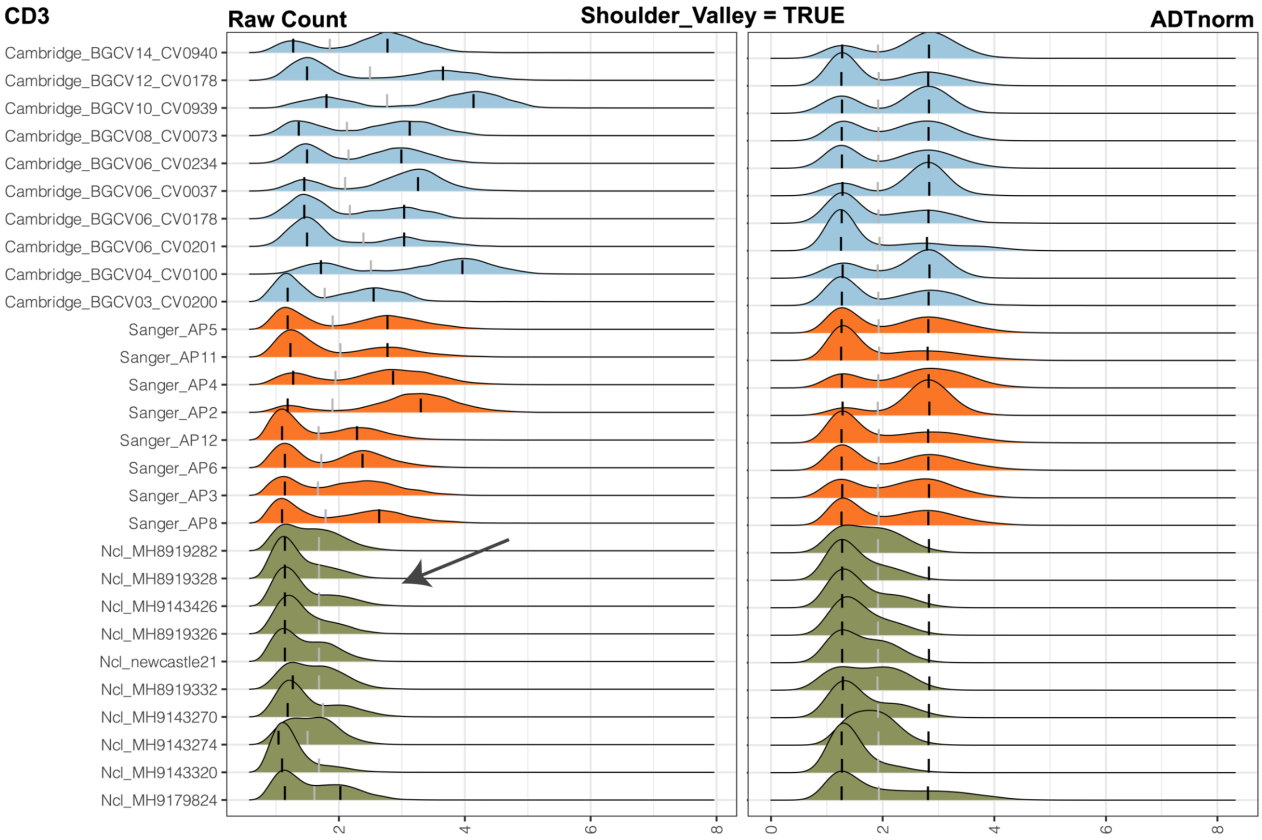


`shoulder_valley` mode will be especially useful when the valley locations vary dramatically. For example, in the following example, CD4 middle peak can be hard to detect for some samples. If we set the valley to be locally minimal between the negative peak and positive peak, it may vary dramatically. Some samples may have the valley before or after the middle peak. Setting `shoulder_valley` to be TRUE with a less negative `shoulder_valley_slope` (such as -0.1) will guarantee that the valley is detected before the middle peak.

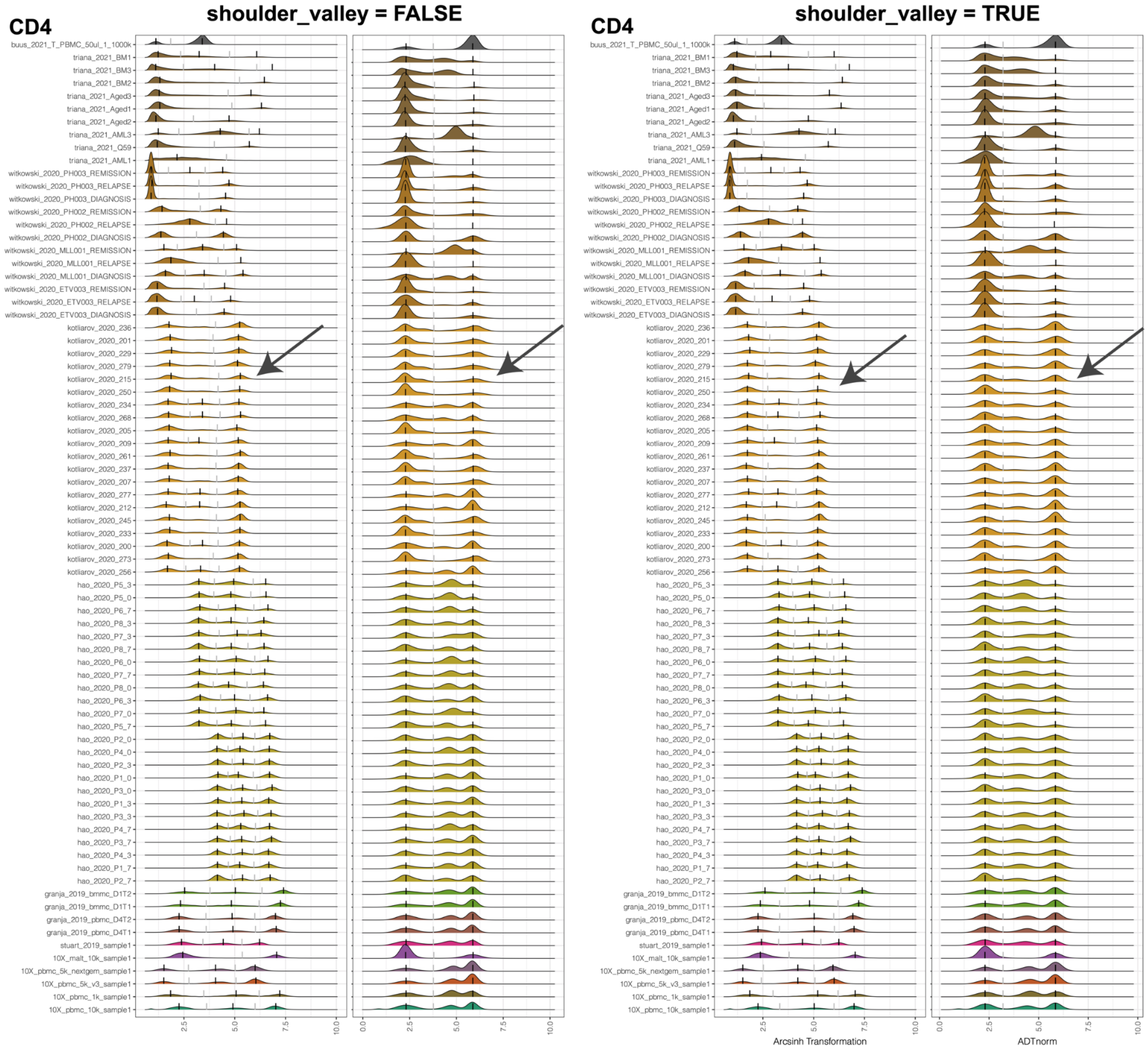


##### 5.8 Bandwidth for the ADT count density distribution that affects the valley location detection

`valley_density_adjust`: Parameter for `density` function: bandwidth used is adjust*bw. This makes it easy to specify values like half the default bandwidth. A lower value, such as 0.5 or 1, can lead to a more accurate valley location but is more prone to minimal local variations. Larger values, such as 3 or 5, can make the valley more consistent across samples.

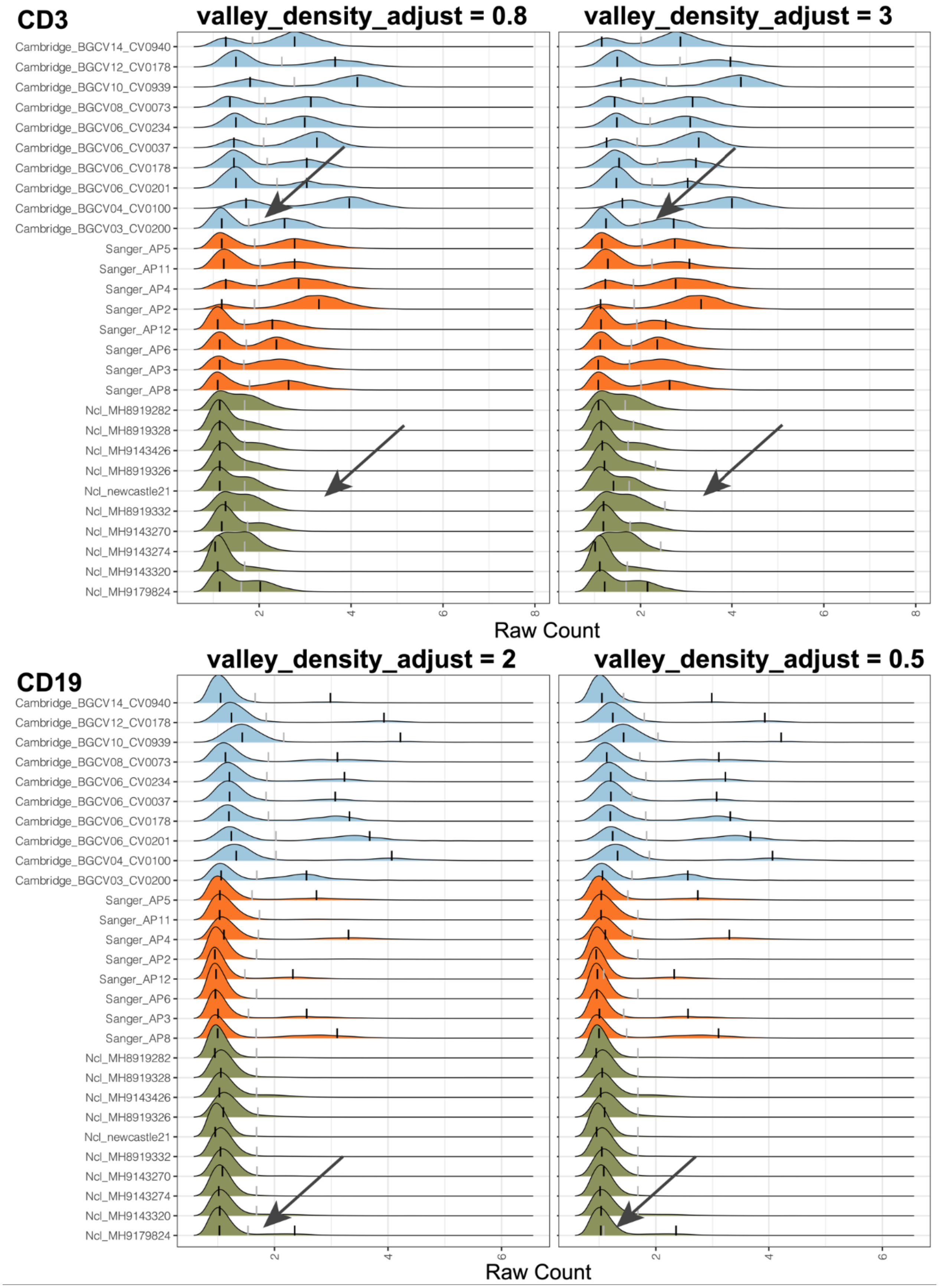
CD3
valley_density_adjust = 0.8
valley_density_adjust = 3
Cambridge_BGCV14_CV0940
Cambridge_BGCV12_CV0178
Cambridge_BGCV10_CV0939
Cambridge_BGCV08_CV0073
Cambridge_BGCV06_CV0234
Cambridge_BGCV06_CV0037
Cambridge_BGCV06_CV0178
Cambridge_BGCV06_CV0201
Cambridge_BGCV04_CV0100
Cambridge_BGCV03_CV0200
Sanger_AP5
Sanger_AP11
Sanger_AP4
Sanger_AP2
Sanger_AP12
Sanger_AP6
Sanger_AP3
Sanger_AP8
Ncl_MH8919282
Ncl_MH8919328
Ncl_MH9143426
Ncl_MH8919326
Ncl_newcastle21
Ncl_MH8919332
Ncl_MH9143270
Ncl_MH9143274
Ncl_MH9143320
Ncl_MH9179824
2
4
6
8
Raw Count
CD19
valley_density_adjust = 2
valley_density_adjust = 0.5
2
4
6
Raw Count

##### 5.9 Imputation method for imputing the missing first valley

`midpoint_type`: Fill in the missing first valley by the midpoint of two positive peaks ("midpoint") or impute by other valleys ("valley").

##### 5.10 A general threshold to determine negative (background) population

`neg_candidate_thres`: The upper bound for the negative peak. Users can refer to their IgG samples to obtain the minimal upper bound of the IgG sample peak. It can be one of the values of asinh(4/5+1), asinh(6/5+1), or asinh(8/5+1) if the right 95% quantile of IgG samples is large. The default is asinh(8/5+1) for raw count input. This filtering function will be disabled if the input is not raw count data and relies on the user to decide whether to give a negative peak upper bound to avoid too many suspicious positive peaks closer to low ADT count.

##### 5.11 Detect outlier valley and impute by the neighbor samples

`detect_outlier_valley`: Detect outlier valley and impute by the neighbor samples. For outlier detection methods, choose from "MAD" (Median Absolute Deviation) or "IQR" (InterQuartile Range). Recommend trying "MAD" first if needed.

##### 5.12 Zero ADT count exclusion and other scaling or transformation as input

`exclude_zeroes`: Indicator to consider zeros as NA, i.e., missing values. Recommend TRUE if zeroes in the data represent dropout, likely for large ADT panels, big datasets, or under-sequenced data. Additionally, if one marker (possibly IgG) only has zero value, this marker will be excluded from the downstream processing.

`cell_x_adt`: Matrix of ADT raw counts in cells (rows) by ADT markers (columns) format. By default, ADTnorm expects raw counts to be provided and arcsinh transformation to be performed by ADTnorm internally. If ADTnorm detects that the input count matrix is a non-integer, it will skip the arcsinh transformation. Also, please note that users need to tune the parameters, such as `neg_candidate_thres`, `bw_smallest_*`, `quantile_clip`, etc., to fit their input transformation.

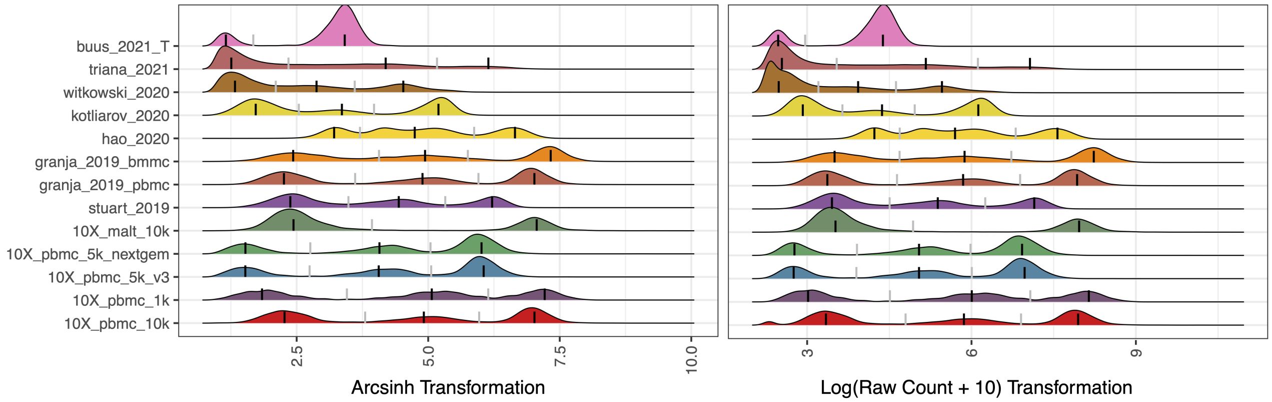


##### 5.13 Ignore the cells with the extremely large ADT count

`quantile_clip`: Implement an upper quantile clipping to avoid warping function errors caused by extremely high expression outlier measurements. Provide the quantile threshold to remove outlier points above such a quantile. The default is 1, meaning no filtering. 0.99 means 99th quantile and points above 99th quantile will be discard. This filtering can be very helpful when the user observes a very long right tail that causes the positive peaks and negative peaks to be squeezed into one narrow peak or just a line in the density plot.

##### 5.14 Manual Adjustment of Landmark Locations by R Shiny

`customize_landmark`: By setting it to be TRUE, ADTnorm will trigger the interactive landmark tuning function and pop out a shiny application for the user's manual setting of the peaks and valleys location. The procedure to adjust the landmarks (peaks and valleys) is indicated below.

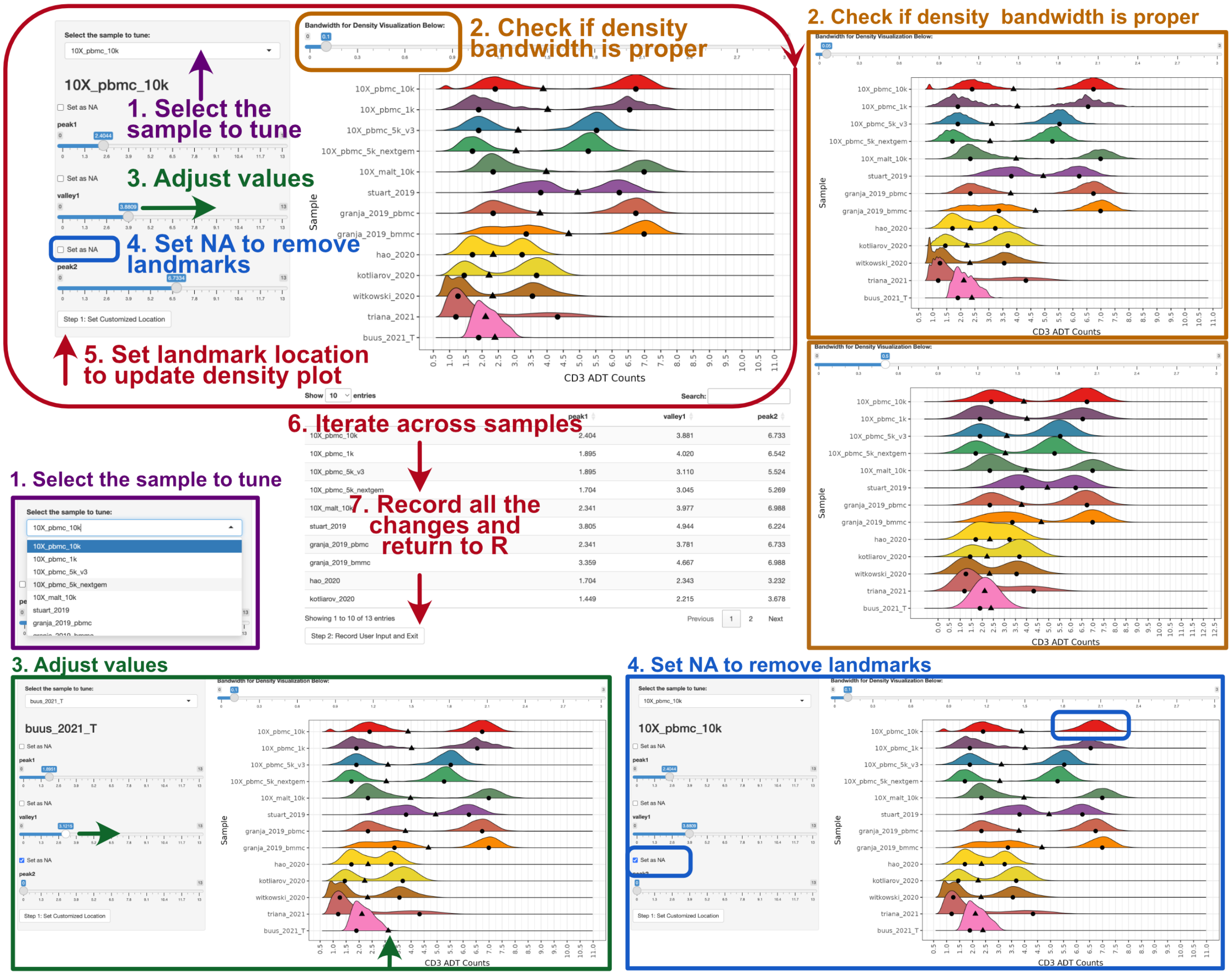


Please note:

- We recommend using this function after initial rounds of ADTnorm normalization with a few parameter tuning attempts. It is better to narrow down a few ADT markers that do need manual tuning and provide the list to `marker_to_process` as the interactive function will pop out for every marker being processed.
- If zigzag discrete negative peaks are observed, user can first increase the "Bandwidth for Density Visualization" at the top of the right panel to smooth out the discrete negative peaks before setting the landmarks.
- Currently, the shiny browser support setting any landmark (peaks or valleys) to NA as missing. However, it does not support inserting new landmark(s). For example, if the marker density distribution shows a triple peak pattern but ADTnorm only detects two peaks across all the samples. Shiny browser

does not allow manual insertion of a new peak and valley, but user can tune the other parameters to push ADTnorm to detect three peaks: specify the target marker as `trimodal_marker`, reducing the `bw_smallest_tri` or setting smaller bandwidth value and specify for the target ADT marker through `bw_smallest_adjustments`.

##### 5.15 Override the peak and valley locations

`override_landmark`: Override the peak and valley locations if prior information is available or the user wants to manually adjust the peak and valley locations for certain markers. The input value of `override_landmark` is in the same format of the intermediate landmark location saved in the rds file if `save_landmark` is set to TRUE. More specifically, the user can overwrite the landmark values by providing a matrix of new landmark values:

```
customized_peak_landmark_list = matrix(rnorm(26), nrow = 13,
    ncol = 2)
rownames(customized_peak_landmark_list) = rownames(peak_mode_res)
customized_valley_landmark_list = matrix(rnorm(26), nrow = 13,
    ncol = 1)
rownames(customized_peak_landmark_list) = rownames(valley_location_res)

override_landmark = list(CD3 = list(peak_landmark_list = customized_peak_landmark_list,
    valley_landmark_list = customized_valley_landmark_list),
    CD4 = list(peak_landmark_list = customized_peak_landmark_list,
        valley_landmark_list = customized_valley_landmark_list))
```

“customized__peak__landmark__list” and “customized__valley__landmark__list” are matrices of customized landmark locations with matching sample names as the rownames.
